# Optimization of pyrazolo[1,5-a]pyrimidines lead to the identification of a highly selective casein kinase 2 inhibitor

**DOI:** 10.1101/2020.06.27.175109

**Authors:** Andreas Krämer, Christian Georg Kurz, Benedict-Tilman Berger, Ibrahim Ethem Celik, Stefan Knapp, Thomas Hanke

## Abstract

Casein kinase 2 (CK2) is a constitutively expressed serine/threonine kinase that has a large diversity of cellular substrates. Thus, CK2 has been associated with a plethora of regulatory functions and dysregulation of CK2 has been linked to disease development in particular to cancer. The broad implications in disease pathology makes CK2 an attractive target. To date, the most advanced CK2 inhibitor is silmitasertib, which has been investigated in clinical trials for treatment of various cancers, albeit several off-targets for silmitasertib have been described. To ascertain the role of CK2 inhibition in cancer, other disease and normal physiology the development of a selective CK2 inhibitor would be highly desirable. In this study we explored the pyrazolo[1,5-a]pyrimidine hinge-binding moiety for the development of selective CK2 inhibitors. Optimization of this scaffold, which included macrocyclization, led to **IC20** (**31**) a compound that displayed high *in vitro* potency for CK2 (*K*_D_ = 12 nM) and exclusive selectivity for CK2. X-ray analysis revealed a canonical type-I binding mode for **IC20**. However, the polar carboxylic acid moiety that is shared by many CK2 inhibitors including silmitasertib was required for potency and reduced somewhat cellular activity. In summary, **IC20** represents a highly selective and potent inhibitor of CK2, which can be used as a tool compound to study CK2 biology and potential new applications for the treatment of diseases.

**Notes:** The authors declare no conflict of interest.

**TOC Figure / Graphical Abstract:** 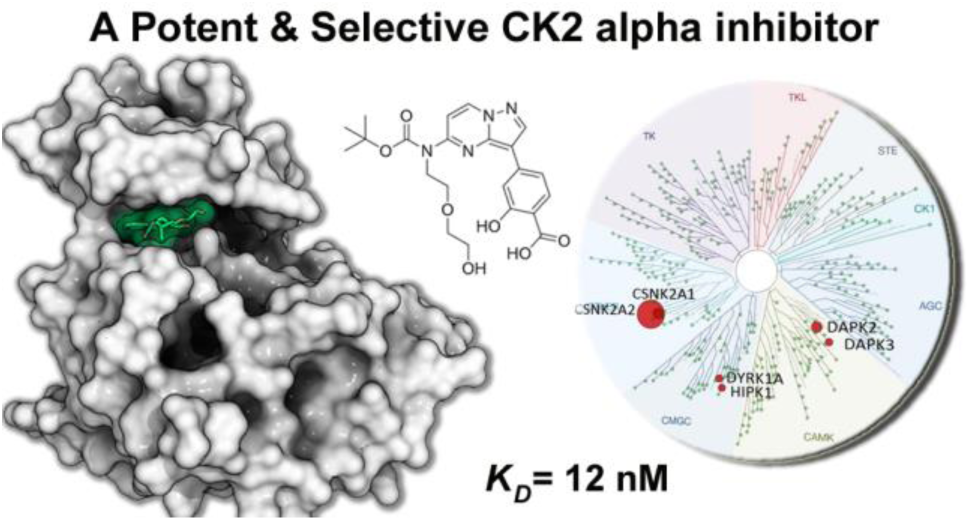

## INTRODUCTION

Protein kinases have emerged as one of the most prominent drug target classes due to their key regulatory role in cell signalling and their deregulation in a large diversity of diseases particular in cancer and inflammatory/autoimmune disorder[1, 2] [3]. Despite the increasing knowledge of their physiology and pathophysiology, the development of selective protein kinase inhibitors is still challenging but necessary for unravelling the molecular functions of kinases and disease applications outside oncology[4]. For the development of selective kinase inhibitors, different strategies have been pursued including the development of covalent inhibitors, allosteric inhibitors or inhibitors using unique binding modes[5, 6].

The most successful way targeting a protein kinase is by addressing the ATP-binding pocket. However, high sequence conservation within the kinase ATP binding pocket and the large size of the kinase family renders selectivity a major challenge[7]. ATP competitive compounds usually rely on a hinge binding motif that anchors the inhibitor to the hinge backbone. A large diversity of hinge binding moieties have been developed that typically employ aromatic heterocycles as hydrogen bond donors and acceptors serving as rigid scaffolds for the formation of strong hydrogen bonds with hinge carbonyls and amid nitrogens. Pyrazolo[1,5-a]pyrimidines represent a privileged and widely explored scaffold that has been used for the development of kinase inhibitors for diverse targets with good selectivity profiles including PIM, RET, JAK1 (pseudokinase domain) and ALK2[8-11] (see **Figure 1A**). Strategies developing selective inhibitors based on this scaffold include unusual binding modes as well as macrocyclization that resulted in the development of selective ALK2 and FLT3 inhibitors[8, 12].

**Figure 1.**
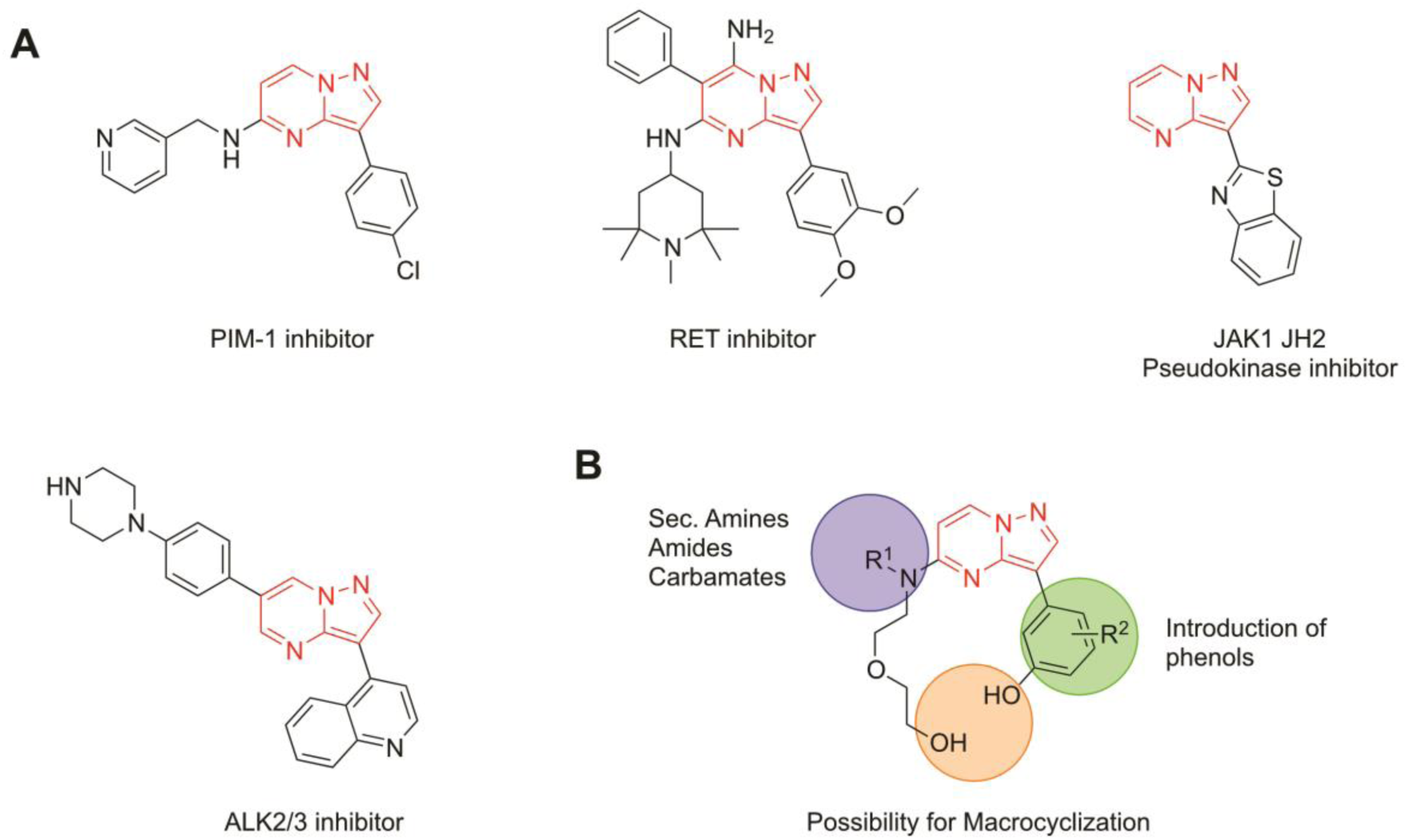
**A** Published pyrazolo[1,5-a]pyrimidines targeting different kinases. **B** Design and modification on pyrazolo[1,5-a]pyrimidine scaffold for targeting CK2.

Protein Kinase CK2 former “Casein kinase 2” is an ubiquitously expressed serine/threonine protein kinase. It plays a role in a wide range of cellular processes such as cell cycle progression, apoptosis, transcription, and viral infections[13]. The transferase is able to utilize both ATP and GTP as a cofactor on a large number of substrates in various pathways[14]. Structural studies revealed that the holoenzyme exists as an *α*_*2*_*β*_*2*_ heterotetramer composed of two interchangeable and highly homologous *α* subunits (*α* and / or *α’*), and two β subunits[15].Unlike many kinases, CK2 is a constitutively active protein kinase whose activity is not directly affected by phosphorylation. Usually kinases can be activated by phosphorylation at specific residues within their activation segments[7]. However CK2 has an N-terminal extension which is able to form stable interactions with the activation segment resulting in a constitutively active enzyme[16]. Thus, the kinase is regulated by expression levels and localisation rather than phosphorylation. Upregulation of CK2 activity has been linked to tumour progression in multiple cancers such as breast, lung, colon, and prostate, making it an attractive target in cancer therapy[13, 17]. Multiple endeavours to target CK2 with small molecules have been explored [18-25]. The most promising inhibitor for translational studies so far is silmitasertib (CX-4945, **1**)[26, 27], which has been granted orphan drug status by the U.S. Food and Drug Administration for treatment of advanced cholangiocarcinoma[28], highlighting the role of CK2 as a therapeutic target. However, despite being used as a chemical probe in a large number of basic research studies, a significant number of off-targets have been reported for this inhibitor. A recent selectivity screen against 238 kinases revealed - besides CK2α and CK2α’ (each with an *IC*_50_ of 1 nM) - potent inhibition of DAPK3 (17 nM), FLT3 (35 nM), TBK1 (35 nM), CLK3 (41 nM), HIPK3 (45 nM), PIM1 (46 nM) and CDK1 (56 nM) limiting its application as a tool compound for mechanistic studies[29]. A pyrazolo[1,5-a]pyrimidine lead compound has been developed by AstraZeneca, which has also been associated with significant off-target activity including low nanomolar activity on DYRK kinases, DAPKs, and HIPKs[30]. In addition, 4,5,6,7-tetrabromo-1*H*-benzimidazole[31] has been an important early tool compound stabilized by halogen bonds to the hinge back bone which, however, also showed considerable off-target activity similar to several natural products such as flavones and flavonols with weak CK2 activity and poor pharmacological properties[32]. Thus, despite intensive research on CK2 and their prominent role in the development of cancer and other diseases no selective chemical probes for these closely related kinases have been reported so far.

In this study, we optimized pyrazolo[1,5-a]pyrimidines to yield highly selective inhibitors of CK2. The expansion of this scaffold yielded the macrocyclic compound IC19 (**32**) and its ring opened analogue IC20 (**31**) that showed excellent potency and selectivity on kinome wide selectivity screens (DiscoverX/Eurofins). NanoBRET™ target engagement assays demonstrated a robust cellular activity for both IC19 (**32**) and IC20 (**31**) in the single digit micromolar range. However, while IC20 (**31**) represents a good cellular tool compound, the comparison with a NanoBRET™ lysed-cells assay revealed a significant drop in cellular activity that would need to be addressed in further optimization of this inhibitor class for *in vivo* use.

## RESULTS

In order to develop a series of CK2 targeting inhibitors, the pyrazolo[1,5-a]pyrimidine hinge binding scaffold was modified at the 3 position by differently decorated phenyl moieties and at the 5 position where an ether linker was coupled via an amine to the pyrimidine ring resulting in secondary amines, which were modified to amides or carbamates. Ring-closure was achieved via a Mitsunobu reaction yielding macrocyclic compounds (see **Figure 1B**).

The synthesis of the double protected hinge-binding moiety (**8**) was performed as described in **Scheme 1**. For the synthesis of pyrazolo[1,5-a]pyrimidinones Gavrin *et al*. reported the observation of two different regioisomers pyrimidin-7-one and pyrimdin-5-one, depending on the synthesis conditions[33]. The synthesis of 4*H*,5*H*-pyrazolo[1,5-a]pyrimidin-5-one (**3**) was performed by starting of commercially available 3-aminopyrazole (**2**) heating with ethyl 3-ethoxyprop-2-enoate in DMF according to the procedure of Gavrin *et al*.[33]. Pyrazolo[1,5-a]pyrimidin-5-one (**3**) was chlorinated with phosphorus oxychloride to the 5-chloropyrazolo[1,5-a]pyrimidine (**4**) and subsequently brominated by using NBS to the brominated hinge-binder 3-bromo-5-chloropyrazolo[1,5-a]pyrimidine (**5**). The linker was introduced by a nucleophilic aromatic substitution of (**5**) with 2-(2-aminoethoxy)ethan-1-ol to (**6**), the hydroxy group was protected with TBDMS and the secondary amine with a BOC-protecting group leading to intermediate (**8**).

In parallel pinacol boronic esters (**12, 14** and **16**) were prepared according to **Scheme 2**. For **12** we started with esterification of 2,4-dihydroxybenzoic acid (**9**) to the methyl ester (**10**). The hydroxy group in para-position to the methyl ester was activated by triflation with Tf_2_O to the corresponding triflate (**11**), which was subsequently converted to the pinacol boronic ester **12** in a Miyaura borylation reaction by using bis(pinacolato)diboron, Pd_2_(dba)_3_ and XPhos in 1,4-dioxane. For the brominated starting material (**13**) the Miyaura borylation was performed with PdCl_2_(dppf) and KOAc in 1,4-dioaxne and for the chlorinated starting material (**15**) Miyaura borylation was performed with Pd_2_(dba)_3_, tricyclohexylphosphine and KOAc in 1,2-dimethoxyethane under microwave conditions at 150 °C to obtain the corresponding pinacol boronic esters (**14** and **16**).

Afterwards both building blocks (**8** and pinacol boronic esters (**12, 14, 16** and **17**)) were coupled using Suzuki coupling to the arylated hinge-binding motifs (**18**–**21**). The cleavage of the silyl ether protecting group was performed with TBAF to the acyclic derivatives (**22**–**25**) and the ring-closure to the macrocyclic compounds (**26**–**29**) were done by a Mitsunobu reaction under high dilutions. The acyclic derivative (**22**) was partially deprotected, either by TFA to obtain the acyclic derivative without the BOC group (**30**), or with LiOH to the acyclic derivative without the methyl ester (**31**).

The same procedure was performed for the macrocyclic compound **23**, which partially deprotected by LiOH to the carboxylic acid derivative **28** or by TFA to the methyl ester **29**, were the methyl ester was cleaved afterwards with LiOH to the double deprotected derivative **30**. Both macrocyclic derivatives **29** and **30** were used for further derivatization. This derivatization was conducted either on the secondary amine by using different acyl chlorides to the corresponding tertiary amides **31** and **32** or by an amide coupling on the carboxylic acid to the secondary amide **33**.

### Pyrazolo[1,5-a]pyrimidines proved to be potent CK2 inhibitors *in vitro*

In order to investigate the selectivity of the acyclic and macrocyclic pyrazolo[1,5-a]pyrimidines, we screened compounds (**23**–**37**) against a panel of 56 kinases using differential scanning fluorimetry (DSF, see **Figure 2** and **Table S1**)[34] The panel included several know off-targets such as PIM, DAPK, DYRK and CLKs as well as kinases important for cell proliferation. This rapid and sensitive assay-format identified several of the synthesized pyrazolo[1,5-a]pyrimidines as potent CK2 inhibitors *in vitro*. Modifications of the macrocyclic and open scaffold led to improved selectivity within the *T*_m_-Panel. Highest *T*_m_-shifts for CK2*α* and CK2*α’* were observed for IC20 (**31**) and its macrocyclic derivative IC19 (**32**). Both compounds shared a carboxylic acid group at the pendant aromatic ring as well as a BOC-group located at the amine attached to the 5-position of the pyrazolo[1,5-a]pyrimidine. Variations at these two positions revealed that both the carboxylic acid and substitution on the 5-amine were essential for potent binding to CK2*α*. However, a nitrile group substituting the carboxylic acid group (**23**) retained some activity in the DSF assay. To better understand the binding mode and the observed SAR of this series of acyclic and macrocyclic pyrazolo[1,5-a]pyrimidines we crystallized cyclic and acyclic derivatives of this inhibitors class with CK2*α*.

**Figure 2.**
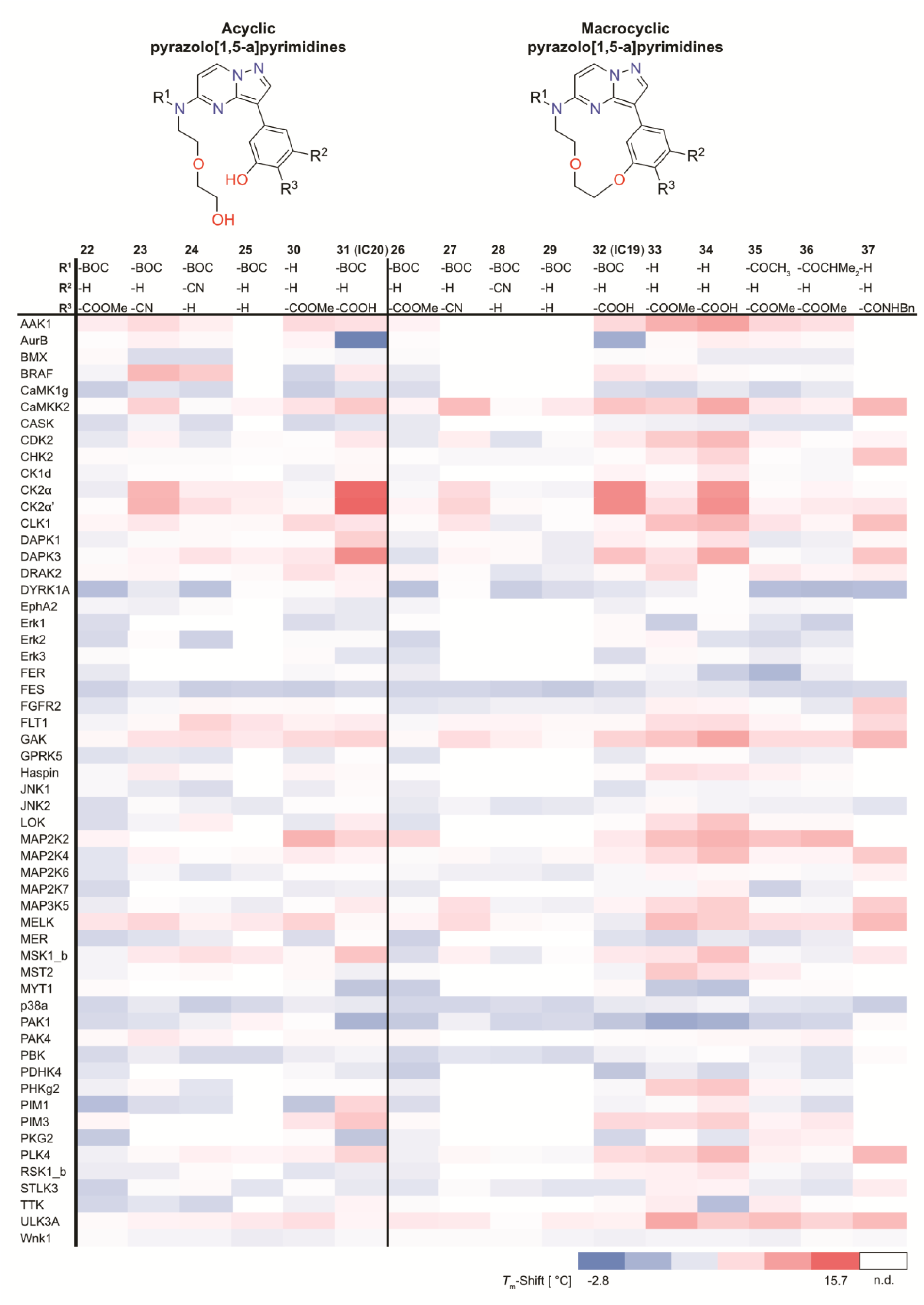
Structure selectivity relationship of acyclic and macrocyclic pyrazolo[1,5-a]pyrimidines. Compounds **22**–**37** were screened in a differential scanning fluorimetry (DSF) assay against a panel of 56 kinases. The results are represented as a heat map ranging from −2.8 °C (colour blue), indicating no stabilization of the kinase, to 15.7 °C (colour red) indicating a high stabilization of the kinase by a compound.

### Crystal structures revealed canonical type-I binding modes

Human CK2*α* (residues 1–337) crystallized in the tetragonal space group P4_3_2_1_2 with two monomers per asymmetric unit. The electron density was clearly interpretable for nearly all residues with the exception of the first and last three amino acids at the N- and C-termini, respectively. The overall conformation of the kinase domain and the ligand binding was identical in each of the two monomers which is highlighted by a RMSD difference of only 0.45 Å over 298 Cα-atoms between the two monomers in the asymmetric unit. As expected, the crystal structure of **34** and IC20 (**31**) revealed that both compounds bound to the ATP binding site. The aromatic pyrazolo[1,5-a]pyrimidine showed a hinge interaction to the backbone nitrogen of residue V116. (see **Figure 3**).

**Figure 3.**
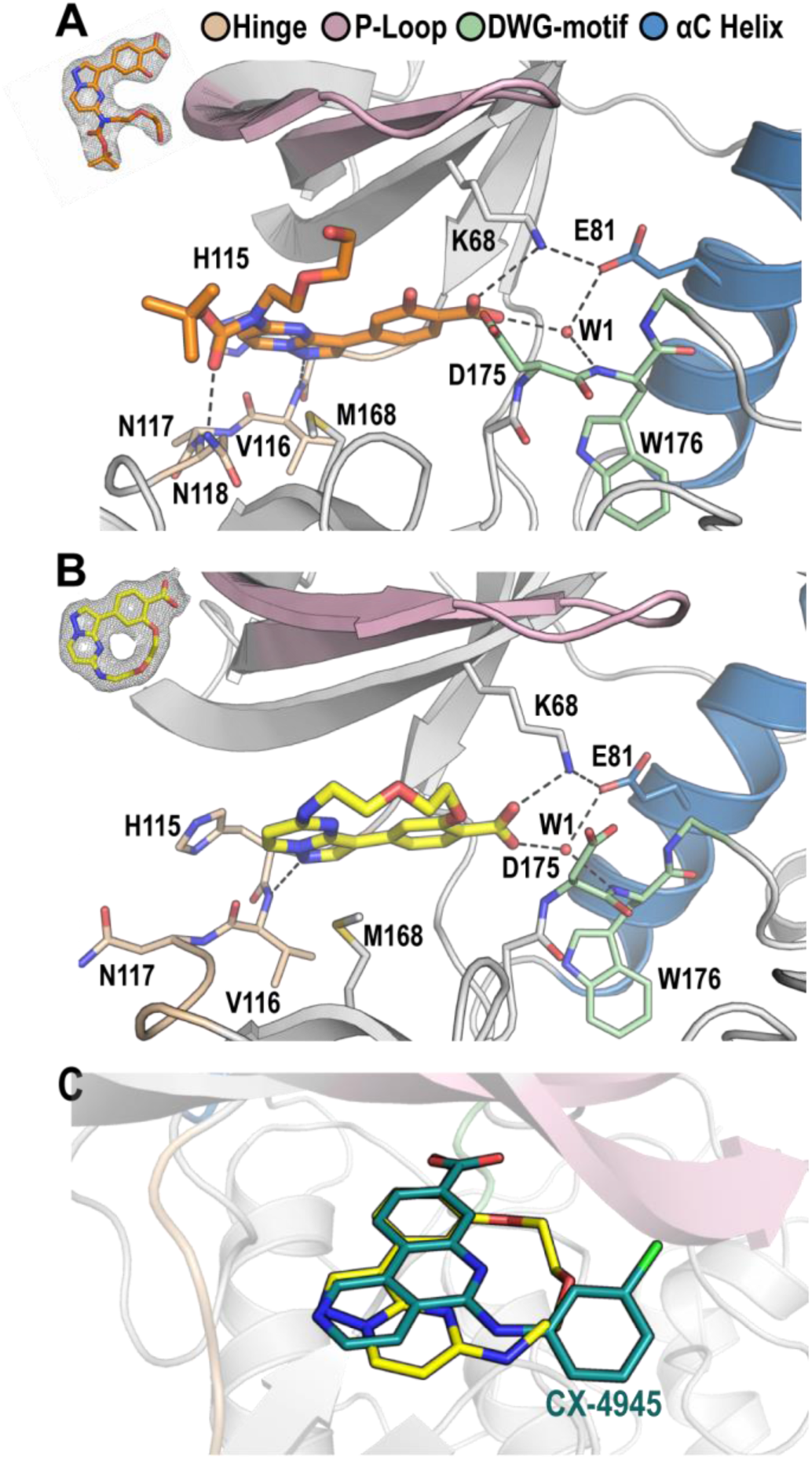
Binding mode of acyclic IC20 (A) and macrocyclic compound 34 (B) and comparison with CX-4945 (C). IC20 (**31**) is shown in orange and **34** in yellow stick representation, respectively. Water molecules are shown as red spheres and important structural motifs are indicated and the same in each panel. Potential hydrogen bonds are shown as black dashed lines. The ligands act as ATP competitive (type I) inhibitors which bind to the backbone of the CK2 hinge region. The overall binding mode of the acyclic and macrocyclic compound is identical. The carboxylate anion of the inhibitors forms a salt bridges to the cationic ammonium sidechain of K68, and a water mediated bridge to E81. All mentioned amino acids are catalytically relevant, underlining the importance of the carboxyl group in this position. The acyclic compound IC20 (**31**) harbours an additional BOC-group attached at the amine on the pyrimidine ring. Its carboxyl group can form an additional hydrogen bond to the side chain N118, providing an explanation for the higher affinity observed for compounds possessing this functional group. The insert on the upper left corner of figure **A** and **B** shows the electron density map of each ligand (2F_o_–F_c_), contoured at 1s. Panel **C** shows an overlay of the macrocycle **34** with the CX-4945. Despite different scaffolds of the compounds, the interaction with the hinge region and the position of the carboxylate anion is essentially the same.

Crucial for CK2 potency was the carboxylic acid group pointing towards the kinase back pocket. The importance of the free acid for binding was demonstrated by the inactivity of methyl esters, which abolished activity completely (see **Figure 2** and **Table S1**). The dramatic effect on inhibitor activity can be explained by a network of polar interactions formed with conserved CK2 back pocket residues including a salt bridge with the VIAK motif lysine K68, a water mediated hydrogen bond to the conserved αC glutamate (E81), and a hydrogen bond to the backbone nitrogen of D175, which is part of the degenerated DFG (DWG)-motif in CK2. It is worth noting that CK2*α* and CK2*α’* are the only human kinases in which the typical DFG motif is replaced by a DWG motif (**Figure 3**). Compound IC20 harbours a BOC-group attached to the amine connected to the 5-position on the pyrazolo-pyrimidine ring system. The carbonyl group of this moiety formed an additional hydrogen bond to the hinge region via the side chain nitrogen of N118. This additional interaction explains the higher *T*_m_-shifts for compounds containing this functional group (see **Figure 2** and **Table S1**). Importantly, the pyrazolo[1,5-a]pyrimidine ring system and the pendant 3-phenyl ring system showed similar orientations in the open and cyclized form explaining that both inhibitors - IC20 (**31**) and IC19 - have similar bioactive binding modes and hence similar selectivity profiles.

### Isothermal titration calorimetry revealed similar binding thermodynamics for IC19 and IC20

The constrained structure of macrocycles suggests that cyclised inhibitors have favourable binding entropy changes (Δ*S*) due to their conformational restrictions, which limit the number of conformations of the unbound inhibitor in comparison to acyclic counterparts[35]. We therefore assessed the thermodynamic properties of binding comparing the macrocyclic and acyclic inhibitors IC19 (**32**) and IC20 (**31**) using isothermal titration calorimetry (ITC) (see **Figure 4**). ITC data revealed that both compounds were highly potent with *K*_D_ values of 12 nM for IC20 (**31**) and 82 nM for IC19 (**32**), respectively. These data correlated well with determined *T*_m_-shifts data (**Table S1**). As expected from their similar binding modes, the enthalpic and entropic contribution to binding differed only marginally. The binding of both compounds was driven by a highly favourable binding enthalpy change driven and opposed by entropy (*T*Δ*S*). Thus, the ring opened compound already rests in a favourable co-planar orientation of both ring systems and the additional constrains of cyclization resulted only in less favourable interactions with the kinase domain. The acyclic IC20 (**31**) therefore represents a better and more potent inhibitor for CK2α.

**Figure 4.**
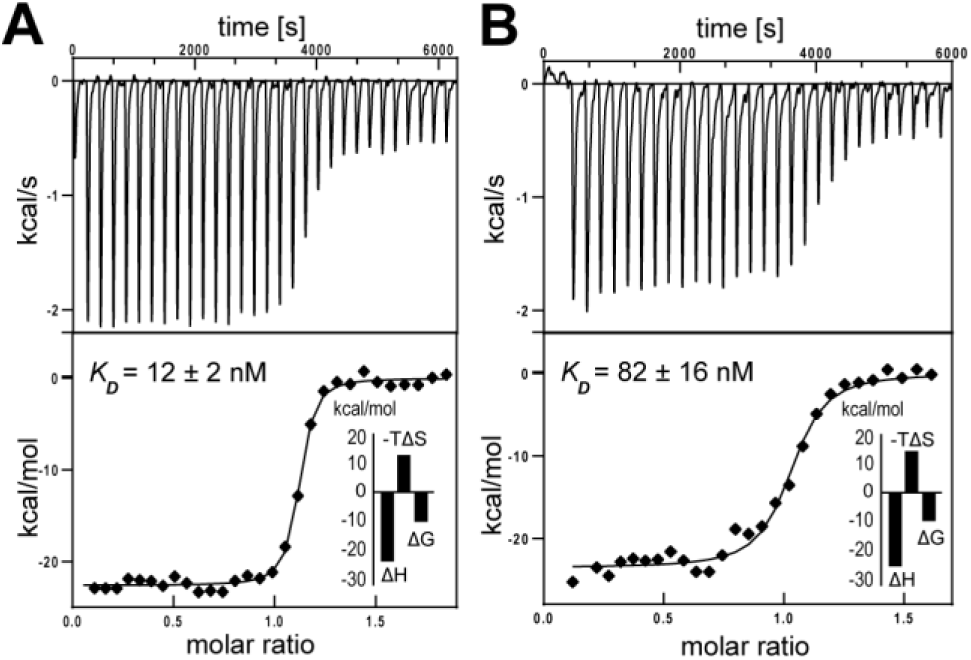
Binding affinity of IC20 (**31, Panel A**) and IC19 (**32, Panel B**) determined by isothermal titration calorimetry reveals two-digit nanomolar binding affinity of these compounds towards CK2. Minor differences are identifiable in their overall Gibbs free energy, as well as in their enthalpic and entropic contribution to CK2 binding.

### IC19 and IC20 were two highly selective CK2 inhibitors

Encouraged by the favourable selectivity data in our DSF experiment, the macrocycle IC19 (**32**) and the acyclic counterpart IC20 (**31**) were assessed on their selectivity against a larger screening panel. The KINOME*scan*® is an assay platform covering 469 kinases (including disease-relevant mutants). Both compounds were screened at a concentration of 1 µM in order to determine their selectivity against the human kinome (see **Figure 5A** and **Table S5**/**S6**). The selectivity of the compounds can be defined by a selectivity score (S-Score), which represents the number of hits divided by the number of assays:

**Figure 5.**
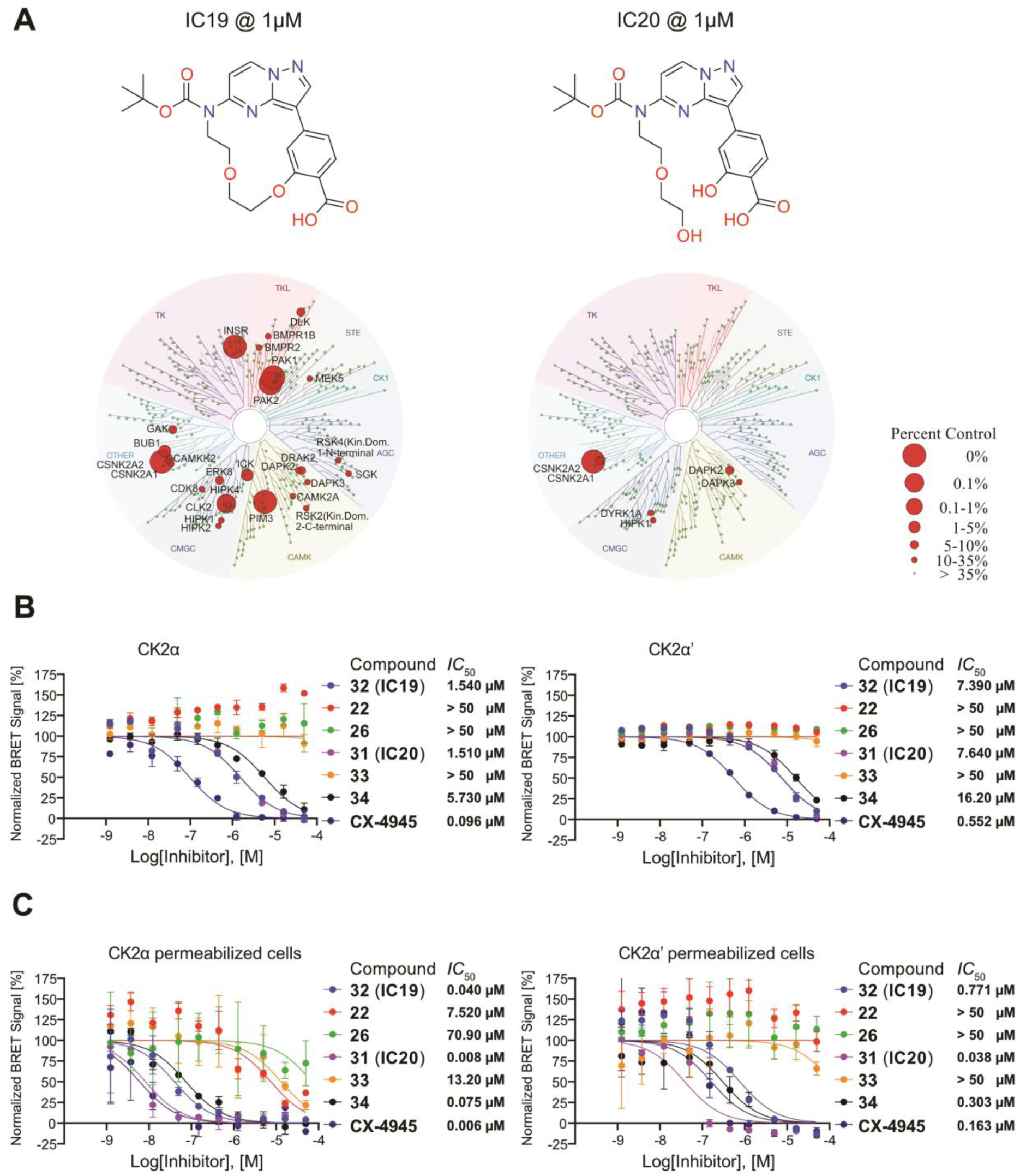
Selectivity and cellular potency of acyclic and macrocyclic pyrazolo[1,5-a]pyrimidines. **A**) **IC19** and **IC20** were screened in a KinomeScan by DiscoverX (now belonging to Eurofins) against a panel of 469 kinases (including disease relevant mutants). Red circles identify potential kinases that are affected by these compounds. **B**) Cellular potency of pyrazolo[1,5-a]pyrimidines on CK2α/CK2α’ determined by NanoBRET™ in HEK293T cells. The respective *IC*_50_ is indicated in the figure legend. **C**) Potency of pyrazolo[1,5-a]pyrimidines on CK2α/CK2α’ determined by NanoBRET™ after permeabilizing the cells. The respective *IC*_50_ is indicated in the figure legend.

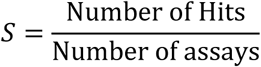

With S_35_ = (number of non-mutant kinases with %Ctrl <35)/(number of non-mutant kinases tested) [Ref: https://www.discoverx.com/services/drug-discovery-development-services/kinase-profiling/kinomescan/scanmax]. The macrocyclic compound IC19 (**32**) had a good selectivity (S_35_ = 0.07) albeit the most favourable selectivity was achieved by the acyclic compound IC20 (**31**) (S_35_ = 0.02). However, we typically observed a false positive rate of approximately 1% in kinome wide screens using KINOME*scan*® technology, which makes follow-up verification necessary. Some targets were already verified in the DSF panel, which led us to believe that PAK1, PIM3, DAPK2 and GAK represented false positive hits for IC19 (**32**) and DYRK1A most likely being a false positive hit for IC20 (**31**). False positives are usually easily identified when highly related kinases isoforms are not inhibited by the investigated compound. It is therefore likely that even though not tested experimentally, also INSR and PAK2 represent false positive hits for IC19 (**32**) because related kinases are missing in the hitlist. For IC20 (**31**), binding to DAPK3 was confirmed by DSF. Gratifyingly, **IC20** (**31**), had an excellent selectivity profile where KINOME*scan*® data revealed exclusive selectivity for the two CK2 isoforms with only minor activity detected for DYRK1A and HIPK1.

### Target engagement by NanoBRET™ revealed low µM potency for IC19 and IC20 in cells

After the encouraging kinome-wide selectivity data for IC19 (**32**) and IC20 (**31**) we were interested in the cellular activity of IC19 (**32**) and IC20 (**31**). To get insight on cellular activity we used NanoBRET™ target engagement assays that are now available for most kinases[36]. NanoBRET™ is a tracer displacement assay that uses bioluminescent resonance energy transfer and allows to measure molecular proximity. The target of interest is fused to a NanoLuc® luciferase, transiently transfected into and expressed in HEK293T cells and after binding of a fluorescent tracer to this target of interest, a BRET signal can be measured. By adding an inhibitor competing with the tracer molecule, the displacement of the fluorescent tracer leads to a dose-dependent decrease of the NanoBRET signal. This allows the determination of an *IC*_50_ in a cellular system under physiological ATP concentrations for full-length kinases. For the CK2α and CK2α’ NanoBRET™ assays we used IC19 (**32**), IC20 (**31**), the non-selective macrocycle **34** and silmitasertib (**1**) as a reference compound. The *IC*_50_s for the macrocycle IC19 (**32**) and acyclic counterpart IC20 (**31**) were a magnitude higher than the binding affinity determined by ITC, with *IC*_50_-values ranging from 1.5 µM to approximately 7.5 µM for both compounds on CK2α and CK2α’, respectively (see **Figure 5B** and **Table S3/S4**). The *IC*_50_ for **34** was slightly higher (5.7 µM/16.2 µM), which was in good agreement with the results from DSF assays. Silmitasertib (**1**) had an *IC*_50_ ranging from 100 nM to 500 nM for CK2α and CK2α’, respectively, which correlated with the cellular activity reported in the literature[37]. We were interested if the decrease in the cellular potency was related to the high intracellular ATP concentration, or other issues related to cellular permeability. As many CK2-inhibitors bear a carboxylic acid, Brear *et al*. recently used a pro-drug strategy to mask the carboxylic acid by a methyl ester to recover the CK2 potency of their compounds in HCT116, Jurkat and A549 cells[18]. Therefore, we repeated the NanoBRET™ assay with the corresponding methyl esters (**22, 26** and **33**). However, after pre-incubation the methyl esters all failed to inhibit CK2α and CK2α’ suggesting that the methyl esters were not rapidly cleaved in the cellular environment during the incubation time of 2 h (see **Figure 5B** and **Table S3/S4**). In order to assess the influence of cell penetration we repeated the NanoBRET™ assay under the same conditions but permeabilized cells using digitonin[36]. We observed potent inhibition of CK2α and CK2α’ in these permeabilised cells. IC20 (**31**) - the most potent inhibitor - showed similar potency as silmitasertib (**1**) with slightly better *IC*_50_-values for CK2α (8 nM) compared to CK2α’ (38 nM) (see **Figure 5C** and **Table S3/S4**), again in very good agreement with our ITC and *T*_m_ data. Thus, we conclude that this new series of pyrazolo[1,5-a]pyrimidines represent highly selective and highly potent inhibitors of CK2α / CK2α’ *in vitro*, however, poor cell penetration reduced cellular activities of IC20 (**31**) and IC19 (**32**) to the low micromolar range.

### IC20 showed no significant cytotoxicity across 60 cancer cell lines

CK2 is a prominent key player in cancer cell biology owing to its central role in cell grow, cell death and cell survival[38, 39]. Antiprolifertive and anti-angiogenic effects have been demonstrated either by using inactive mutants of CK2 catalytic subunit α or by CK2 inhibitors such as silmitasertib [29, 40]. Therefore, we screened our selective and potent CK2 inhibitor IC20 (**31**) against the NCI-60 cell line panel, at the National Cancer Institute (NCI) (see **Figure 6**). The results reported for the one-dose assay at an inhibitor concentration of 10 μM were displayed relative to the no-drug control with a value of 100 representing no growth inhibition whereas negative values indicated cytotoxicity (https://dtp.cancer.gov/discovery_development/nci-60/methodology.htm). For IC20 (**31**) we observed no cytotoxicity in all 60 cell lines, but we observed some growth inhibition in several different cell lines such as the glioblastoma cell lines SNB-19 and U251. The modest anti-proliferative effects may be explained by the selectivity of IC20 (**31**) as many off-targets of silmitasertib (**1**) have been linked to cell proliferation and survival. However, more cell-based studies will be required to clarify the role of CK2 in cancer cell survival and the potential benefits inhibiting this target in cancer patients.

**Figure 6.**
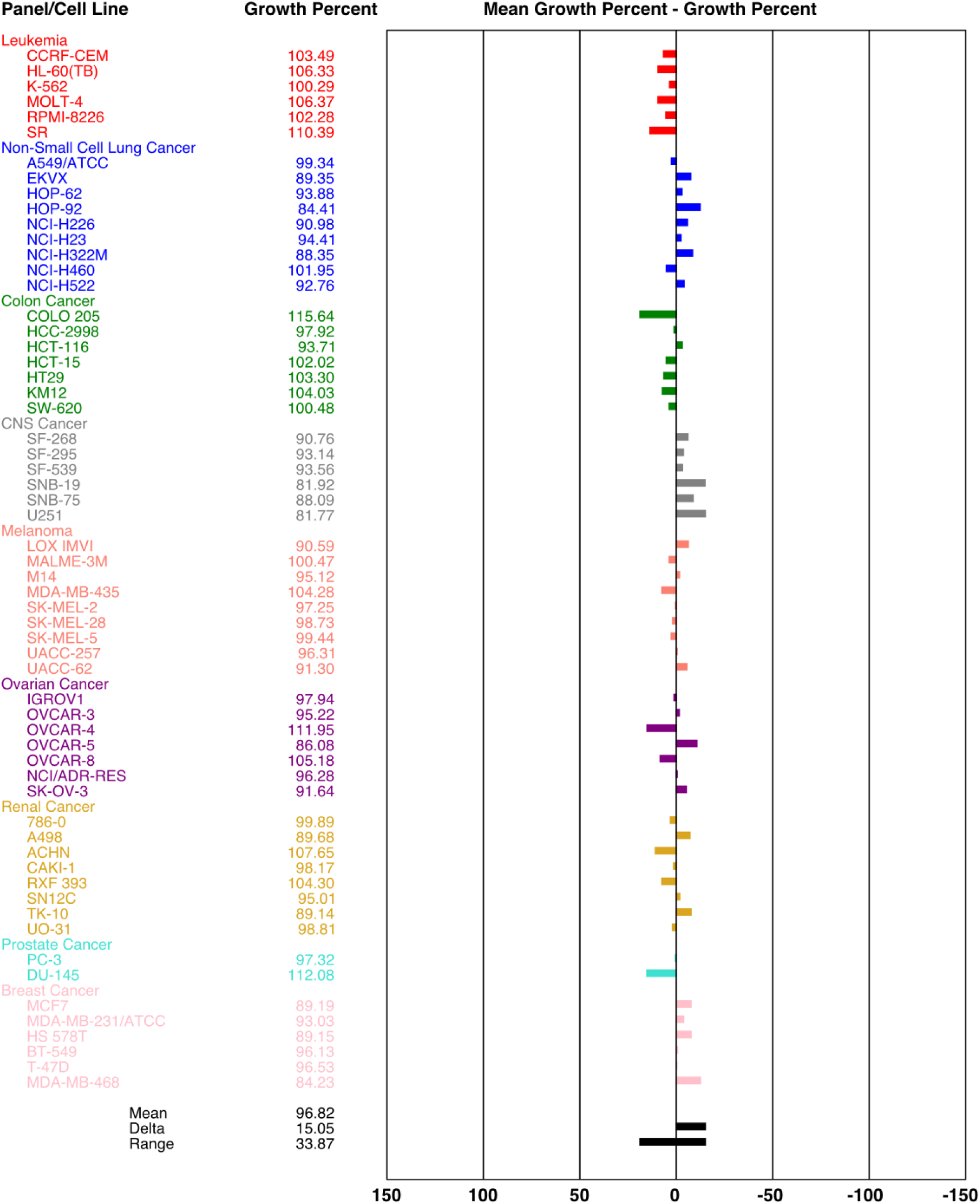
Growth inhibition of IC20 in 60 different cancer cell lines from the National Cancer Institute (NCI60-Panel). IC20 (**31**) was screened in a one-dose format at 10 µM against 60 different cell lines to investigate the potential of growth inhibition. Values of 100 indicate no growth inhibition. Values between 100 and 0 indicate growth inhibition.

### Conclusion/Discussion

To sum up, in this comparative study of acyclic and macrocyclic pyrazolo[1,5-a]pyrimidines, we have seen that small modifications of this scaffold can have a significant effect on the selectivity of this series of compounds. In previous literature, pyrazolo[1,5-a]pyrimidines have been described as PIM1, RET, JAK1 or ALK2/3 kinase inhibitors. In this study we optimized this scaffold to a highly selective Casein kinase 2 inhibitor, with IC20 (**31**) and IC19 (**32**). Both compounds have comparable potency to silmitasertib (**1**) *in vitro* and are low micromolar inhibitors in a cellular system of CK2. The selectivity of this compound has been improved over other well-known CK2 inhibitors including silmitasertib (**1**), TBB[41] or Emodin[42]. CK2 has been suggested to play a significant role in cancer cellular growth and progression and silmitasertib (**1**) is currently tested in several clinical trials for the treatment of various cancer diseases like basal cell carcinoma (NCT03897036), Medulloblastoma (NCT03904862), Multiple Myeloma (NCT01199718), Cholangiocarcinoma (NCT02128282), Breast cancer (NCT00891280) or kidney cancer (NCT03571438). Even though it is not deniable that CK2 plays a central role in cancer diseases it is likely that antiproliferative effects of silmitasertib (**1**) are also a consequence of inhibiting off-targets[29]. Main off-targets of silmitasertib (**1**) reported to have antiproliferative effects are for example the cyclin-dependent kinase (CDK1)[43, 44], PIM-1 proto-oncogene, serine/threonine kinase (PIM1)[45], Homeodomain-interacting protein kinase 3 (HIPK3)[46, 47], TANK-binding kinase 1 (TBK1)[48] or FMS-like tyrosine kinase 3 (FLT3)[49, 50]. More recently some more selective CK2 inhibitors have been reported, including GO289 by Oshima *et al*.[51] or an allosteric inhibitor CAM4066 by Brear *et al*.[18]. However, the allosteric inhibitors are still weak and would require optimization and the selectivity of GO289 has not been comprehensively assessed. Our developed tool compound IC20 (**31**) therefore represents a valuable tool for functional studies. Replacing the carboxylic acid moiety with a back pocket binding motif that will not increase cell penetration of this series would need to be addressed in the future.

## EXPERIMENTAL PROCEDURES

### Cloning

Catalytic domain residues of CK2*α* (residues 1-337) were amplified from cDNA provided by the mammalian gene collection (MGC) and cloned into the vector pNIC28-Bsa4 by ligation-independent cloning[52]. The vector includes a Tobacco Etch Virus (TEV) cleavable site, N-terminal His_6_-tag, a kanamycin resistance. A lambda protein phosphatase was co-expressed to remove possible autophosphorylation.

### Expression and Purification of CK2*α*

Transformed BL21(DE3) cells were grown in Terrific Broth medium containing 50 mg/ml kanamycin. Protein expression was induced at an OD_600_ of 2 by using 0.5 mM isopropyl-thio-galactopyranoside (IPTG) at 18°C for 12 hours. Cells expressing His_6_-tagged CK2*α* were lysed in lysis buffer containing 50 mM HEPES pH 7.5, 500 mM NaCl, 25 mM imidazole, 5% glycerol and 0.5 mM Tris(2-carboxyethyl)phosphine (TCEP) by sonication. After centrifugation, the supernatant was loaded onto a Nickel-Sepharose column equilibrated with 30 ml lysis buffer. The column was washed with 60 ml lysis buffer. Proteins were eluted by an imidazole step gradient (50, 100, 200, 300 mM). Fractions containing protein were pooled together and dialysed overnight with 1 L of final buffer (25 mM HEPES pH 7.5, 500 mM NaCl, 0.5 mM TCEP) at 4°C. Additionally, TEV protease was added (protein:TEV 1:20 molar ratio) to remove the tag. At the next day, the protein solution was loaded onto Nickel-Sepharose column beads again to remove TEV protease and cleaved Tag. The flow through fraction and the wash fraction (25 mM imidazole) containing the protein was concentrated to approx. 4-5 mL and loaded onto a Superdex 75 16/60 HiLoad gel filtration column equilibrated with final buffer. The protein was concentrated to approx. 9 mg/ml.

### Crystallization

CK2*α* was crystallized using the sitting-drop vapour diffusion method by mixing protein (8–10 mg/ml) and well solutions in 2:1, 1:1, and 1:2 ratios using the Mosquito (SPT Labtech). The reservoir solution contained 0.2 M ammonia sulphate, 0.1 M 2-(N-morpholino)ethanesulfonic acid (MES) pH 6.5 and 31-35% (v/v) polyethylene glycol monomethyl ether 5000 (PEG MME 5k). Complex structures were achieved by soaking the crystals at least for 24h with the desired inhibitor. Final concentration of the inhibitor was approx. 1 mM.

### Data collection, Structure Solution and Refinement

Diffraction data were collected at beamline X06DA (Villigen, CH) at a wavelength of 1.0 Å at 100 K. The reservoir condition supplemented with 20% ethylene glycol was used as cryoprotectant. Data were processed using XDS[53] and scaled with aimless. The PDB structure with the accession code 3PE2[54] was used as an initial search MR model using the program MOLREP[55]. The final model was built manually using Coot[56] and refined with REFMAC5[57]. Data collection and refinement statistics are summarized in Table S2.

### Thermal Stability Measurements

Thermal melting experiments were carried out with an Mx3005p realtime PCR machine (Agilent). Proteins were buffered in 25 mM HEPES (pH 7.5), 500 mM NaCl and were assayed in a 96-wellplate at a final concentration of 2 µM in a 20 µl volume. Inhibitors were added at a final concentration of 10 µM. SYPRO-Orange (Molecular Probes) was added as a fluorescence probe at a dilution of 1 in 5000. Excitation and emission filters were set to 465 nm and 590 nm, respectively. The temperature was raised with a step of 3°C per minute, and fluorescence readings were taken at each interval. The temperature dependence of the fluorescence was approximated by the equation

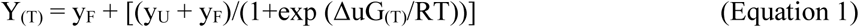

where Δu*G* is the difference in unfolding free energy between the folded and unfolded state, *R* is the gas constant, and *y*_F_ and *y*_U_ are the fluorescence intensity of the probe in the presence of completely folded and unfolded protein, respectively[58]. The baselines of the denatured and native state were approximated by a linear fit. The observed temperature shifts, Δ*T*_m_ obs, for each inhibitor were recorded as the difference between the transition midpoints of sample and reference wells containing protein without inhibitor and were determined by nonliner least-squares fit. Measurements were performed in triplicate.

### Isothermal calorimetry

The *K*_D_ of the compounds was determined by isothermal calorimetry. The sample cell, sample syringe, and injection syringe were all equilibrated with gel filtration buffer (25 mM HEPES pH 7.5, 500 mM NaCl, 0.5 mM TCEP). The sample cell was filled with compound solution (10-15 μM in gel filtration buffer), and the injection syringe was filled with protein solution (85 μM in gel filtration buffer). After equilibration of the device (Nano ITC, TA Instruments), protein was injected over 30 injections, 8 μL each time. For the analysis of the data, the baseline and integration points were defined to determine binding heats. Data was fitted to the Boltzmann equation to determine thermodynamic parameters

### NanoBRET™ target engagement assays

The assay was performed as described previously[36, 59]. In brief: Full-length kinase ORF (Promega) cloned in frame with a NanoLuc-vector (as indicated in Table SX) was transfected into HEK293T cells using FuGENE HD (Promega, E2312) and proteins were allowed to express for 20 h. Serially diluted inhibitor and NanoBRET™ Kinase Tracer (as indicated in Table S8) were pipetted into white 384-well plates (Greiner 781 207) using an ECHO 550 acoustic dispenser (Labcyte). The corresponding transfected cells were added and reseeded at a density of 2 × 10^5^ cells/mL after trypsinization and resuspension in Opti-MEM without phenol red (Life Technologies). The system was allowed to equilibrate for 2 hours at 37 °C and 5 % CO_2_ prior to BRET measurements. To measure BRET, NanoBRET™ NanoGlo Substrate + Extracellular NanoLuc Inhibitor (Promega, N2160) were added as per the manufacturer’s protocol, and filtered luminescence was measured on a PHERAstar plate reader (BMG Labtech) equipped with a luminescence filter pair (450 nm BP filter (donor) and 610 nm LP filter (acceptor)). Competitive displacement data were then plotted using GraphPad Prism 8 software using a normalized 3-parameter curve fit with the following equation: Y=100/(1+10^((X-LogIC50))). For the assay in the lysed format, digitonin (Promega, G9441) was used as per the manufacturer’s instructions at a final concentration of 50 ng/mL.

### Organic synthesis

All commercial chemicals and reagents were obtained from Fluorochem, Merck, TCI, abcr or Apollo Scientific. Unless otherwise indicated, the purity was ≥ 95%. The solvents used in analytical grade were obtained from Fisher Scientific, Merck and VWR Chemicals and all dry solvents, with AcroSeal septum, from Acros Organics. All performed thin layer chromatography (TLC) was done with silica gel on aluminum foils (60 Å pore diameter) obtained from Macherey-Nagel and visualized with ultraviolet light (λ = 254 and 365 nm). The nuclear magnetic resonance spectroscopy (NMR) was performed with spectrometers from Bruker (Karlsruhe, Germany): DPX250, AV400 or AV500 MHz spectrometer. Chemical shifts are reported in parts per million (ppm) in the scale relative to DMSO-*d*_*6*_, 2.50 ppm for ^1^H NMR and 39.52 for ^13^C NMR. Coupling constants (*J*) were reported in hertz (Hz) and multiplicities were designated as followed: s (singlet), bs (broad singlet), d (doublet), dd (double doublet), ddd (doublet of doublet of doublets), t (triplet), dt (doublet of triplets), q (quartet), m (multiplet). Mass spectra were obtained on two ThermoFisher Surveyor MSQ’s, (one MSQ coupled to a Camag TLC-MS interface 2 for direct measurements of TLC spots) measuring in the positive- and/or negative-ion mode. High resolution mass spectra were recorded on a MALDI LTQ Orbitrap XL system from Thermo Scientific (Waltham, MA, USA). Purity of the synthesized compounds was determined with a Shimadzu (Duisburg, Germany) LC-20AD HPLC set at 254 and 280 nm equipped with a Shimadzu LCMS-2020 detector using a Phenomenex LTD (Aschaffenburg, Germany) Luna 10 µm 21.2 × 250 mm reversed phase (C_18_) column. As mobile phase Milli-Q water (A) and acetonitrile (B) + 0.1% formic acid were used with a flowrate of 1 mL/min. The gradient was running over 25 min starting with 95% A and 5% B, going down on 10% A and 90% B and finishing at 95% A and 5% B. The purity of all synthesized final compounds was 95% or higher.

#### Synthesis of 4*H*,5*H*-pyrazolo[1,5-*a*]pyrimidine-5-one (3)

To a solution of 3-aminopyrazole (5.00 g, 60.18 mmol) and ethyl 3-ethoxyacrylat (13.01 g, 90.27 mmol) in dimethylformamide (100 mL) was added cesium carbonate (29.41 g, 90.27 mmol). The reaction mix was heated at 110 °C for 4 h. Afterwards, the solvent was removed and the crude product purified with silica gel column chromatography with a mobile phase of ethyl acetate, petroleum ether and tetrahydrofuran (ratio gradually ranging from 1:0:0 to 0:1:2). The white solid was the title compound (3.90 g, 48%). ^1^H NMR (250 MHz, DMSO-*d*_*6*_): *δ* 12.08 (s, 1H, N*H*), 8.48 (dd, *J* = 7.9, 0.8 Hz, 1H, Het*H*), 7.76 (d, *J* = 2.0 Hz, 1H, Het*H*), 5.94 (d, *J* = 7.9 Hz, 1H, Het*H*), 5.82 (dd, *J* = 2.0, 0.8 Hz, 1H, Het*H*) ppm.

#### Synthesis of 5-Chloropyrazolo[1,5-*a*]pyrimidine (4)

4*H*,5*H*-Pyrazolo[1,5-*a*]pyrimidine-5-one (**3**) (3.90 g, 28.86 mmol) was dissolved in phosphoryl chloride (79.65 g, 519.48 mmol) and stirred under argon atmosphere for 2 h at 120 °C. The reaction was quenched in portions with ice water (200 mL) and it was extracted with ethyl acetate. The organic layers were dried over MgSO_4_ and the solvent was removed in vacuo. Further purification was done with silica gel column chromatography with a mobile phase of *n-*hexane and ethyl acetate (ratio of 1:1). The desired compound was obtained as a white solid (3.53 g, 80%). ^1^H NMR (250 MHz, DMSO-*d*_*6*_): *δ* 9.17 (dd, *J* = 7.3, 0.9 Hz, 1H, Het*H*), 8.28 (d, *J* = 2.3 Hz, 1H, Het*H*), 7.12 (d, *J* = 7.2 Hz, 1H, Het*H*), 6.72 (dd, *J* = 2.4, 0.9 Hz, 1H, Het*H*) ppm.

#### Synthesis of 3-Bromo-5-chloropyrazolo[1,5-*a*]pyrimidine (5)

5-Chloropyrazolo[1,5-*a*]pyrimidine (**4**) (3.43 g, 22.34 mmol) was suspended in dimethylformamide (60 mL) and *N*-bromosuccinimide (4.37 g, 24.57 mmol) was added in small portions while stirring. Alter 1 h at room temperature, the reaction was quenched with water (150 mL). The precipitation was vacuum filtered and washed with water. After drying, the yellowish solid was the desired compound (4.89 g, 94%). ^1^H NMR (250 MHz, DMSO-*d*_*6*_): *δ* 9.22 (d, *J* = 7.3 Hz, 1H, HetH), 8.44 (s, 1H, HetH), 7.23 (d, *J* = 7.2 Hz, 1H, HetH) ppm.

#### Synthesis of 2-[2-({3-Bromopyrazolo[1,5-*a*]pyrimidine-5-yl}amino)ethoxy]ethan-1-ol (6)

A solution of 3-bromo-5-chloropyrazolo[1,5-*a*]pyrimidine (**5**) (4.89 g, 21.03 mmol), 2-(2-aminoethoxy)ethanol (2.43 g, 23.13 mmol) and *N,N*-diisopropylethylamine (3.26 g, 25.24 mmol) in acetonitrile (63 mL) was stirred under reflux for 16 h. Next step was the evaporation of the solvent in vacuo. The residue was taken up in ethyl acetate and washed with water and brine. After drying over MgSO_4_ the solvent was evaporated under reduced pressure and the crude product was purified with silica gel column chromatography with a mobile phase of petroleum ether and tetrahydrofuran (ratio gradually ranging from 3:1 to 0:1). The title compound was obtained as a white solid (3.36 g, 53%). ^1^H NMR (250 MHz, DMSO-*d*_*6*_): *δ* 8.45 (d, *J* = 7.6 Hz, 1H Het*H*), 7.87 (s, 1H, Het*H*), 7.73 (t, *J* = 5.2 Hz, 1H, N*H*), 6.35 (d, *J* = 7.6 Hz, 1H, Het*H*), 4.60 (s, 1H, O*H*), 3.65 – 3.42 (m, 8H, C*H*_2_) ppm.

#### Synthesis of 3-Bromo-5-(8,8,9,9-tetramethyl-4,7-dioxa-1-aza-8-siladecan-1-yl)pyrazolo[1,5-*a*]pyrimidine (7)

2-[2-({3-Bromopyrazolo[1,5-*a*]pyrimidine-5-yl}amino)ethoxy]ethan-1-ol (**6**) (2.46 g, 8.17 mmol) and triethylamine (1.65 g, 16.34 mmol) are dissolved in dry dimethylformamide (40 mL). While stirring, *tert*-butyldimethylsilyl chloride (1.85 g, 12.26 mmol) was added in small portions. After 1 h at room temperature the solvent was removed under reduced pressure and the residue was dissolved in water. The aqueous layer was extracted with ethyl acetate, the organic layer washed with brine and dried over MgSO_4_. The solvent was removed under reduced pressure and further purification was carried out by a silica gel column chromatography with a mobile phase of *n-*hexane an ethyl acetate (ratio of 1:1). The title compound was obtained as a white solid (3.30 g, 97%). ^1^H NMR (250 MHz, DMSO-*d*_*6*_): *δ* 8.44 (d, *J* = 7.6 Hz, 1H, Het*H*), 7.86 (s, 1H, Het*H*), 7.70 (t, *J* = 5.4 Hz, 1H, N*H*), 6.33 (d, *J* = 7.6 Hz, 1H, Het*H*), 3.72 – 3.48 (m, 8H, C*H*_2_), 0.83 (s, 9H, (C*H*_3_) _3_), 0.02 (s, 6H, Si(C*H*_3_)_2_) ppm.

#### Synthesis of tert-Butyl *N*-{3-bromopyrazolo[1,5-*a*]pyrimidin-5-yl}-*N*-(2-{2-[(tert-butyldimethylsilyl)oxy]ethoxy}ethyl)carbamate (8)

Triethylamine (1.99 g, 19.67 mmol) was added to a solution of 3-bromo-5-(8,8,9,9-tetramethyl-4,7-dioxa-1-aza-8-siladecan-1-yl)pyrazolo[1,5-*a*]pyrimidine (**7**) (6.81 g, 16.39 mmol), di-*tert*-butyldicarbonate (8.77 g, 40.16 mmol) and 4-(dimethylamino)pyridine (0.04 g, 0.33 mmol) in tetrahydrofuran (50 mL). The reaction mixture was heated to reflux for 3 h and afterwards, the solvent was evaporated. The residue was taken up in ethyl acetate and washed with water and brine. After drying over MgSO_4_ the organic layer was removed under reduced pressure. Further purification was done with silica gel column chromatography with a mobile phase of *n-*hexane an ethyl acetate (ratio of 5:1). The desired compound was yielded as a brownish oil (8.24 g, 97%). ^1^H NMR (250 MHz, DMSO-*d*_*6*_): *δ* 8.94 (d, *J* = 7.8 Hz, 1H, Het*H*), 8.23 (s, 1H, Het*H*), 7.42 (d, *J* = 7.8 Hz, 1H, Het*H*), 4.17 (t, *J* = 6.1 Hz, 2H, C*H*_2_), 3.69 (t, *J* = 6.1 Hz, 2H, C*H*_2_), 3.58 (t, *J* = 5.1 Hz, 2H, C*H*_2_), 3.47 (t, *J* = 5.3 Hz, 2H, C*H*_2_), 1.51 (s, 9H, (C*H*_3_)_3_), 0.78 (s, 9H, (C*H*_3_)_3_), −0.07 (s, 6H, Si(C*H*_3_)_2_) ppm.

#### Synthesis of Methyl 2,4-dihydroxybenzoate (10)

A solution of 2,4-dihydroxybenzoic acid (**9**) (20.00 g, 129.77 mmol) in dry methanol (100 mL) and concentrated sulfuric acid (2.6 mL) was stirred under reflux for 16 h. After reaction completion, the solvent was removed under reduced pressure and the residue was taken up with ethyl acetate. The organic layer was washed with brine, dried over MgSO_4_ and evaporated in vacuo. Further purification was done with silica gel column chromatography with a mobile phase of *n-*hexane and ethyl acetate (ratio of 4:1). The desired compound was obtained as a white solid (7.76 g, 36%). ^1^H NMR (250 MHz, DMSO-*d*_*6*_): *δ* 10.70 (s, 1H, O*H*), 10.44 (s, 1H, O*H*), 7.64 (d, *J* = 8.7 Hz, Ph*H*), 6.37 (dd, *J* = 8.8, 2.4 Hz, 1H, Ph*H*), 6.29 (d, *J* = 2.3 Hz, 1H, Ph*H*), 3.84 (s, 3H, OC*H*_3_) ppm.

#### Synthesis of Methyl 2-hydroxy-4-(trifluoromethansulfonyloxy)benzoate (11)

Methyl-2,4-dihydroxybenzoate (10) (7.63 g, 45.39 mmol), pyridine (3.53 g, 45.39 mmol) and 4-(dimethylamino)-pyridine (0.21 g, 1.70 mmol) were dissolved in dry methylene chloride (120 mL) under argon atmosphere. The solution was cooled to 0 °C and 1 M trifluoromethanesulfonic anhydride in methylene chloride (25 mL, 22.69 mmol) was added dropwise. The reaction was allowed to stir for 16 h at room temperature and was quenched with 10% hydrochloric acid (20 mL). The aqueous layer was extracted with ethyl acetate and the organic layer was dried over MgSO_4_ and evaporated in vacuo. The crude product was purified with silica gel column chromatography with a mobile phase of *n-*hexane and ethyl acetate (ratio gradually ranging from 1:0 to 20:1). The white solid obtained is the title compound (6.50 g, 95%). ^1^H NMR (250 MHz, DMSO-*d*_*6*_): *δ* 10.88 (s, 1H, O*H*), 7.92 (d, *J* = 8.8 Hz, 1H, Ph*H*), 7.13 (d, *J* = 2.5 Hz, 1H, Ph*H*), 7.05 (dd, *J* = 8.8, 2.5 Hz, 1H, Ph*H*), 3.88 (s, 3H, OC*H*_3_) ppm.

#### Synthesis of Methyl 2-hydroxy-4(4,4,5,5-tetramethyl-1,3,2-dioxaborolan-2-yl)benzoate (12)

To a solution of methyl-2-hydroxy-4-(((trifluoromethyl)sulfonyl)oxy)benzoate (**11**) (2.29 g, 7.61 mmol) in dry 1,4-dioxane (50 mL), bis(pinacolato)diboron (2.90 g, 11.42 mmol), potassium acetate (2.24 g, 22.83 mmol) and Pd(dppf)Cl_2_·DCM (0.31 g, 0.38 mmol) were added. The reaction solution was flushed with argon for 10 min and stirred at 100 °C for 3.5 h. Subsequently the reaction was diluted with ethyl acetate and water and the layers were separated. The organic layer was washed with brine and dried over MgSO_4_. After removal of the solvent under reduced pressure, the crude product was purified with silica gel column chromatography with a mobile phase of *n-*hexane and ethyl acetate (ratio gradually ranging from 1:0 to 30:1). The title compound was obtained as a white solid (1.69 g, 80%). ^1^H NMR (500 MHz, DMSO-*d*_*6*_): *δ* 10.36 (s, 1H, O*H*), 7.76 (d, *J* = 7.8 Hz, 1H, Ph*H*), 7.31 – 7.11 (m, 2H, Ph*H*), 3.88 (s, 3H, OC*H*_3_), 1.29 (s, 12H, C*H*_3_) ppm.

#### Synthesis of 2-Hydroxy-4-(4,4,5,5-tetramethyl-1,3,2-dioxaborolan-2-yl)benzonitrile (14)

A suspension of 4-bromo-2-hydroxybenzonitrile (**13**) (0.50 g, 2.52 mmol), Pd(dppf)Cl_2_ (0.06 g, 0.08 mmol), potassium acetate (0.74 g, 7.56 mmol) and bis(pinacolato)diboron (0.70 g, 2.77 mmol) in dry 1,4-dioxane was stirred at 100 °C for 3.5 h. Thereafter the reaction mixture was filtrated over celite and washed with 1,4-dioxane. The organic layer was washed with water, dried over Na_2_SO_4_ and the solvent was removed in vacuo. Further purification was done by silica gel column chromatography with a mobile phase of *n-*hexane an ethyl acetate (ratio gradually ranging from 10:1 to 3:1). The title compound was yielded as a yellowish solid (0.37 g, 60%). ^1^H NMR (500 MHz, DMSO-*d*_*6*_): *δ* 11.06 (s, 1H, O*H*), 7.59 (d, *J* = 7.7 Hz, 1H, Ph*H*), 7.30 (s, 1H, Ph*H*), 7.16 (dd, *J* = 7.7, 1.0 Hz, 1H, Ph*H*), 1.29 (s, 12H, (C*H*_3_)_4_) ppm.

#### Synthesis of 3-Hydroxy-5-(4,4,5,5-tetramethyl-1,3,2-dioxaborolan-2-yl)benzonitrile (16)

In dimethoxymethane (10 mL) 3-chloro-5-hydroxybenzonitrile (**15**) (1.00 g, 6.50 mmol), Pd_2_(dba)_3_ (0.18 g, 0.20 mmol), tricyclohexylphosphine (0.22 g, 0.78 mmol), potassium acetate (1.44 g, 15.12 mmol) and bis(pinacolato)diboron (1.80 g, 7.16 mmol) were suspended and the reaction mix was stirred for 2 h at 150 °C under microwave radiation. Afterwards the reaction mixture was filtrated over celite and washed with diethyl ether. The organic layer was washed with water, dried over Na_2_SO_4_ and the solvent was evaporated. Purification was performed by silica gel column chromatography with a mobile phase of *n-*hexane an ethyl acetate (ratio gradually ranging from 3:1 to 1:1). The yellowish solid (0.52 g, 32%) was the desired compound. ^1^H NMR (250 MHz, DMSO-*d*_*6*_): *δ* 10.22 (s, 1H, O*H*), 7.38 – 7.35 (m, 2H, Ph*H*), 7.24 (dd, *J* = 2.6, 1.5 Hz, 1H, Ph*H*), 1.29 (s, 12H, (C*H*_3_)_4_) ppm.

#### Synthesis of Methyl 4-[5-(2,2,3,3,13,13-hexamethyl-11-oxo-4,7,12-trioxa-10-aza-3-silatetradecan-10-yl)pyrazolo[1,5-*a*]pyrimidin-3-yl]-2-hydroxybenzoate (18)

*tert*-Butyl *N*-{3-bromopyrazolo[1,5-*a*]pyrimidin-5-yl}-*N*-(2-{2-[(tert-butyldimethylsilyl)oxy]ethoxy}ethyl)carbamate (**8**) (2.15 g, 4.17 mmol) and Methyl 2-hydroxy-4(4,4,5,5-tetramethyl-1,3,2-dioxaborolan-2-yl)benzoate (**12**) (2.32 g, 8.34 mmol) were suspended together with Pd_2_(dba)_3_ (38 mg, 0.04 mmol), XPhos (0.24 g, 0.50 mmol) and potassium phosphate (3.54 g, 16.68 mmol) in 1,4-dioxane (27 mL) and water (11 mL). The reaction was running for 40 min at 110 °C. After diluting with ethyl acetate layers were separated, the organic one was washed with water and brine and dried over MgSO_4_. The solvent was evaporated in vacuo. Purification was performed by silica gel column chromatography with a mobile phase of *n-*hexane an ethyl acetate (ratio gradually ranging from 5:1 to 1:1). The white solid obtained was the desired compound (2.23 g, 91%). ^1^H NMR (250 MHz, DMSO-*d*_*6*_): *δ* 10.62 (s, 1H, O*H*), 8.99 (d, *J* = 7.8 Hz, 1H, Het*H*), 8.74 (s, 1H, Het*H*), 7.79 (d, *J* = 8.1 Hz, 1H, Ph*H*), 7.71 – 7.66 (m, 2H, Ph*H*), 7.49 (d, *J* = 7.8 Hz, 1H, Het*H*), 4.23 (t, *J* = 6.2 Hz, 2H, C*H*_2_), 3.90 (s, 3H, OC*H*_3_), 3.77 (t, *J* = 6.2 Hz, 2H, C*H*_2_), 3.61 (t, *J* = 5.0 Hz, 2H, C*H*_2_), 3.48 (t, *J* = 4.9 Hz, 2H, C*H*_2_), 1.53 (s, 9H, (C*H*_3_)_3_), 0.76 (s, 9H, (C*H*_3_)_3_), −0.08 (s, 6H, Si(C*H*_3_)_2_) ppm.

#### Synthesis of *tert*-Butyl *N*-(2-{2-[(*tert*-butyldimethylsilyl)oxy]ethoxy}ethyl)-*N*-[3-(4-cyano-3-hydroxyphenyl)pyrazolo[1,5-*a*]pyrimidin-5-yl]carbamate (19)

The title compound was synthesized according to the procedure of (**18**) using 2-hydroxy-4-(4,4,5,5-tetramethyl-1,3,2-dioxaborolan-2-yl)benzonitrile (**14**) (0.32 g, 1.31 mmol). Purification by silica gel column chromatography with a mobile phase of *n-*hexane an ethyl acetate (ratio gradually ranging from 5:1 to 1:1), afforded the title compound as a yellowish solid (0.39 g, 81%). ^1^H NMR (500 MHz, DMSO-*d*_*6*_): *δ* 11.04 (s, 1H, O*H*), 9.00 (d, *J* = 7.7 Hz, 1H, Het*H*), 8.65 (s, 1H, Het*H*), 7.72 (s, 1H, Ph*H*), 7.62 – 7.57 (m, 2H, Ph*H*), 7.44 (d, *J* = 7.7 Hz, 1H, Het*H*), 4.30 (t, *J* = 5.9 Hz, 2H, C*H*_2_), 3.71 (t, *J* = 5.9 Hz, 2H, C*H*_2_), 3.53 (t, *J* = 5.1 Hz, 2H, C*H*_2_), 3.42 (t, *J* = 5.0 Hz, 2H, C*H*_2_), 1.52 (s, 9H, (C*H*_3_)_3_), 0.74 (s, 9H, (C*H*_3_)_3_), −0.12 (s, 6H, Si(C*H*_3_)_2_) ppm.

#### Synthesis of *tert*-Butyl *N*-(2-{2-[(*tert*-butyldimethylsilyl)oxy]ethoxy}ethyl)-*N*-[3-(3-cyano-5-hydroxyphenyl)pyrazolo[1,5-*a*]pyrimidin-5-yl]carbamate (20)

The desired compound was synthesized according to the procedure of (**18**) using 3-hydroxy-5-(4,4,5,5-tetramethyl-1,3,2-dioxaborolan-2-yl)benzonitrile (**16**) (0.44 g, 1.78 mmol). Purification was done by silica gel column chromatography with a mobile phase of *n-*hexane an ethyl acetate (ratio gradually ranging from 5:1 to 1:1). The yellow solid (0.43 g, 66%) was the titled product. ^1^H NMR (400 MHz, DMSO-*d*_*6*_): *δ* 10.18 (s 1H, O*H*), 8.98 (d, *J* = 7.2 Hz, 1H, Het*H*), 8.70 (s, 1H, Het*H*), 7.91 (s, 1H, Ph*H*), 7.80 (s, 1H, Ph*H*), 7.44 (d, *J* = 7.8 Hz, 1H, Het*H*), 6.96 (s, 1H, Ph*H*), 4.25 (t, *J* = 6.0 Hz, 2H, C*H*_2_), 3.75 (t, *J* = 5.9 Hz, 2H, C*H*_2_), 3.57 (t, *J* = 5.0 Hz, 2H, C*H*_2_), 3.45 (t, *J* = 5.0 Hz, 2H, C*H*_2_), 1.52 (s, 9H, (C*H*_3_)_3_), 0.74 (s, 9H, (C*H*_3_)_3_), −0.11 (s, 6H, Si(C*H*_3_)_2_) ppm.

#### Synthesis of *tert-*Butyl *N*-(2-{2-[(*tert*-butyldimethylsilyl)oxy]ethoxy}ethyl)-*N*-[3-(3-hydroxyphenyl)pyrazolo[1,5-*a*]pyrimidin-5-yl]carbamate (21)

The desired compound was synthesized according to the procedure of (**18**) using 3-(4,4,5,5-tetramethyl-1,3,2-dioxaborolan-2-yl)phenol (**17**) (0.50 g, 2.27 mmol). Purification by silica gel column chromatography with a mobile phase of *n-*hexane and ethyl acetate (ratio gradually ranging from 5:1 to 1:1), afforded the title compound as a yellowish solid (0.69 g, 86%). ^1^H NMR (500 MHz, DMSO-*d*_*6*_): *δ* 9.33 (s, 1H, O*H*), 8.94 (d, *J* = 7.7 Hz, 1H, Het*H*), 8.54 (s, 1H, Het*H*), 7.48 (dt, *J* = 7.7, 1.2 Hz, 1H, Ph*H*), 7.44 (t, *J* = 2.0 Hz, 1H, Ph*H*), 7.37 (d, *J* = 7.7 Hz, 1H, Het*H*), 7.18 (t, *J* = 7.9 Hz, 1H, Ph*H*), 6.63 (ddd, *J* = 8.0, 2.4, 1.0 Hz, 1H, Ph*H*), 4.22 (t, *J* = 6.2 Hz, 2H, C*H*_2_), 3.74 (t, *J* = 6.2 Hz, 2H, C*H*_2_), 3.58 (dd, *J* = 5.7, 4.4 Hz, 2H, C*H*_2_), 3.46 (dd, *J* = 5.7, 4.4 Hz, 2H, C*H*_2_), 1.52 (s, 9H, (C*H*_3_)_3_), 0.77 (s, 9H, (C*H*_3_)_3_), −0.07 (s, 6H, Si(C*H*_3_)_2_) ppm.

#### Synthesis of Methyl 4-(5-{[(*tert*-butoxy)carbonyl][2-(2-hydroxyethoxy)ethyl]amino}pyrazolo[1,5-*a*]pyrimidin-3-yl)-2-hydroxybenzoate (22)

To a solution of methyl 4-[5-(2,2,3,3,13,13-hexamethyl-11-oxo-4,7,12-trioxa-10-aza-3-silatetradecan-10-yl)pyrazolo[1,5-*a*]pyrimidin-3-yl]-2-hydroxybenzoate (**18**) (2.20 g, 3.75 mmol) in tetrahydrofuran (32 mL) was added a 1 M tetrabutylammonium fluoride solution (5.6 mL, 5.63 mmol) in tetrahydrofuran. The reaction stirred at room temperature. After 3 h the solvent was removed under reduced pressure. Subsequently the residue was taken up with ethyl acetate, washed with water and brine and the solvent was evaporated again. Further purification was done with silica gel column chromatography with a mobile phase of *n-*hexane an ethyl acetate (ratio gradually ranging from 1:1 to 1:2). The title compound was yielded as white solid (1.43 g, 81%). ^1^H NMR (500 MHz, DMSO-*d*_*6*_): *δ* 10.63 (s, 1H, O*H*), 9.00 (d, *J* = 7.8 Hz, 1H, Het*H*), 8.77 (s, 1H, Het*H*), 7.82 (d, *J* = 8.3 Hz, 1H, Ph*H*), 7.74 – 7.69 (m, 2H, Ph*H*), 7.50 (d, *J* = 7.8 Hz, 1H, Het*H*), 4.54 (t, *J* = 5.3 Hz, 1H, O*H*), 4.23 (t, *J* = 6.3 Hz, 2H, C*H*_2_), 3.91 (s, 3H, OC*H*_3_), 3.77 (t, *J* = 6.3 Hz, 2H, C*H*_2_), 3.48 – 3.42 (m, 4H, C*H*_2_), 1.53 (s, 9H, (C*H*_2_)_3_) ppm. ^13^C NMR (126 MHz, DMSO-*d*_*6*_): *δ* 169.44, 160.84, 153.87, 152.84, 143.83, 143.11, 139.81, 136.63, 130.20, 116.38, 112.71, 109.37, 106.21, 104.80, 82.56, 72.25, 67.99, 60.22, 52.38, 45.84, 27.67 ppm. MS (ESI+) *m/z:* 473.04 [M + H]^+^. HRMS *m/z:* [M + H]^+^ calcd for C_23_H_29_N_4_O_7_, 473.20308; found 473.20216. HPLC: t_R_ = 16.064, purity ≥ 95%.

#### Synthesis of *tert-*Butyl *N*-[3-(4-cyano-3-hydroxyphenyl)pyrazolo[1,5-*a*]pyrimidin-5-yl]-*N*-[2-(2-hydroxyethoxy)ethyl]carbamate (23)

The desired product was synthesized according to the procedure of (**22**) using *tert*-butyl *N*-(2-{2-[(*tert*-butyldimethylsilyl)oxy]ethoxy}ethyl)-*N*-[3-(4-cyano-3-hydroxyphenyl)pyrazolo[1,5-*a*]pyrimidin-5-yl]carbamate (**19**) (0.38 g, 0.69 mmol). Purification was carried out by silica gel column chromatography with a mobile phase of *n-*hexane and ethyl acetate (ratio gradually ranging from 1:1 to 1:5). The title compound was yielded as yellowish solid (0.29 g, 96%). ^1^H NMR (500 MHz, DMSO-*d*_*6*_): *δ* 11.04 (s, 1H, O*H*), 9.00 (d, *J* = 7.8 Hz, 1H, Het*H*), 8.65 (s, 1H, Het*H*), 7.71 (s, 1H, Ph*H*), 7.65 – 7.58 (m, 2H, Ph*H*), 7.46 (d, *J* = 7.8 Hz, 1H, Het*H*), 4.53 (s, 1H, O*H*), 4.27 (t, *J* = 6.0 Hz, 2H, C*H*_2_), 3.71 (t, *J* = 6.0 Hz, 2H, C*H*_2_), 3.45 – 3.36 (m, 4H, C*H*_2_), 1.53 (s, 9H, (C*H*_3_)_3_) ppm. ^13^C NMR (126 MHz, DMSO-*d*_*6*_): *δ* 160.37, 153.93, 152.88, 143.55, 142.99, 138.45, 136.55, 133.35, 117.49, 116.64, 111.96, 106.27, 105.33, 95.62, 82.47, 72.08, 68.09, 60.20, 45.75, 27.68 ppm. MS (ESI+) *m/z:* 440.20 [M + H]^+^. HRMS *m/z:* [M + H]^+^ calcd for C_22_H_26_N_5_O_5_, 440.19285; found 440.19244. HPLC: t_R_ = 13.903, purity ≥ 95%.

#### Synthesis of *tert-*Butyl *N*-[3-(3-cyano-5-hydroxyphenyl)pyrazolo[1,5-*a*]pyrimidin-5-yl]-*N*-[2-(2-hydroxyethoxy)ethyl]carbamate (24)

The title compound was synthesized according to the procedure of (**22**) using *tert*-butyl *N*-(2-{2-[(*tert*-butyldimethylsilyl)oxy]ethoxy}ethyl)-*N*-[3-(3-cyano-5-hydroxyphenyl)pyrazolo[1,5-*a*]pyrimidin-5-yl]carbamate (**20**) (0.42 g, 0.76 mmol). Further purification was performed by silica gel column chromatography with a mobile phase of *n-*hexane an ethyl acetate (ratio gradually ranging from 1:1 to 1:5). The yellowish solid (0.30 g, 90%) was obtained as the title product. ^1^H NMR (500 MHz, DMSO-*d*_*6*_): *δ* 10.22 (s, 1H, O*H*), 8.98 (d, *J* = 7.8 Hz, 1H, Het*H*), 8.70 (s, 1H, Het*H*), 7.91 (t, *J* = 1.5 Hz, 1H, Ph*H*), 7.81 (t, *J* = 1.7 Hz, 1H, Ph*H*), 7.44 (d, *J* = 7.7 Hz, 1H, Het*H*), 6.96 (dd, *J* = 2.4, 1.4 Hz, 1H, Ph*H*), 4.50 (t, *J* = 5.2 Hz, 1H, O*H*), 4.23 (t, *J* = 6.1 Hz, 2H, C*H*_2_), 3.75 (t, *J* = 6.1 Hz, 2H, C*H*_2_), 3.47 – 3.38 (m, 4H, C*H*_2_), 1.53 (s, 9H, (C*H*_3_)_3_) ppm. ^13^C NMR (126 MHz, DMSO-*d*_*6*_): *δ* 153.73, 152.87, 143.31, 142.67, 136.46, 135.00, 119.50, 119.02, 116.86, 115.18, 112.33, 105.75, 104.92, 82.45, 72.19, 68.16, 60.20, 45.90, 27.67 ppm. MS (ESI+) *m/z:* 440.20 [M + H]^+^. HRMS *m/z:* [M + H]^+^ calcd for C_22_H_26_N_5_O_5_, 440.19285; found 440.19240. HPLC: t_R_ = 13.906, purity ≥ 95%.

#### Synthesis of *tert-*Butyl *N*-[2-(2-hydroxyethoxy)ethyl]-*N*-[3-(3-hydroxyphenyl)pyrazolo[1,5-*a*]pyrimidin-5-yl]carbamate (25)

The title compound was synthesized according to the procedure of (**22**) using *tert-*butyl *N*-(2-{2-[(*tert*-butyldimethylsilyl)oxy]ethoxy}ethyl)-*N*-[3-(3-hydroxyphenyl)pyrazolo[1,5-*a*]pyrimidin-5-yl]carbamate (**21**) (0.26 g, 0.49 mmol). Purification was done by silica gel column chromatography with a mobile phase of *n-*hexane and ethyl acetate (ratio gradually ranging from 1:1 to 1:2). The yellow solid (0.18 g, 0.43 mmol) was the desired product. ^1^H NMR (500 MHz, DMSO-*d*_*6*_): *δ* 9.35 (s, 1H, O*H*), 8.94 (d, *J* = 7.7 Hz, 1H, Het*H*), 8.55 (s, 1H, Het*H*), 7.52 – 7.46 (m, 1H, Ph*H*), 7.45 (t, *J* = 2.0 Hz, 1H, Ph*H*), 7.39 (d, *J* = 7.8 Hz, 1H, Het*H*), 7.19 (t, *J* = 7.9 Hz, 1H, Ph*H*), 6.64 (dd, *J* = 8.0, 2.5 Hz, 1H, Ph*H*), 4.52 (t, *J* = 5.2 Hz, 1H, O*H*), 4.21 (t, *J* = 6.3 Hz, 2H, C*H*_2_), 3.74 (t, *J* = 6.3 Hz, 2H, C*H*_2_), 3.50 – 3.38 (m, 4H, C*H*_2_), 1.52 (s, 9H, (C*H*_3_)_3_) ppm. ^13^C NMR (126 MHz, DMSO-*d*_*6*_): *δ* 157.57, 153.06, 152.89, 142.93, 142.23, 136.17, 133.21, 129.51, 116.26, 112.84, 112.27, 107.85, 104.63, 82.26, 72.20, 68.04, 60.22, 45.74, 27.68 ppm. MS (ESI+) *m/z:* 415.22 [M + H]^+^. HRMS *m/z:* [M + H]^+^ calcd for C_21_H_27_N_4_O_5_, 415.19760; found 415.19744. HPLC: t_R_ = 13.681, purity ≥ 95%.

#### Synthesis of 13-*tert*-Butyl 5-methyl 7,10-dioxa-13,17,18,21-tetraazatetracyclo[12.5.2.1^2^,^6^.0^17^,^20^]docosa-1(20),2,4,6(22),14(21),15,18-heptaene-5,13-dicarboxylate (26)

A suspension of triphenylphosphine (0.83 g, 3.17 mmol) and sodium sulfate in dry toluene (240 mL) was added dropwise with a solution of diisopropyl azodicarboxylate (0.64 g, 3.17 mmol) in dry toluene (65 mL). Afterwards methyl 4-(5-{[(*tert*-butoxy)carbonyl][2-(2-hydroxyethoxy)ethyl]amino}pyrazolo[1,5-*a*]pyrimidin-3-yl)-2-hydroxybenzoate (**22**) (0.50 g, 1.06 mmol) was dissolved in 2-methyltetrahydrofuran (20 mL) and this solution was added dropwise. Reaction took place over 3 h at 90 °C under argon atmosphere. Sodium sulfate was separated by filtration and the solvent was evaporated in vacuo. Purification was achieved by silica gel column chromatography with a mobile phase of *n-*hexane an ethyl acetate (ratio gradually ranging from 9:1 to 0:1). The yellowish solid obtained was the title compound (0.44 g, 91%). ^1^H NMR (500 MHz, DMSO-*d*_*6*_): *δ* 9.01 (d, *J* = 7.8 Hz, 1H, Het*H*), 8.75 (d, *J* = 1.2 Hz, 1H, Ph*H*), 8.71 (s, 1H, Het*H*), 7.69 (d, *J* = 8.1 Hz, 1H, Ph*H*), 7.55 (d, *J* = 7.8 Hz, 1H, Het*H*), 7.43 (dd, *J* = 8.1, 1.4 Hz, 1H, Ph*H*), 4.39 (t, *J* = 6.0 Hz, 2H, C*H*_2_), 4.09 (t, *J* = 6.6 Hz, 2H, C*H*_2_), 3.92 (t, *J* = 6.6 Hz, 2H, C*H*_2_), 3.88 (t, *J* = 6.0 Hz, 2H, C*H*_2_), 3.78 (s, 3H, OC*H*_3_), 1.54 (s, 9H, (C*H*_3_)_3_) ppm. ^13^C NMR (126 MHz, DMSO-*d*_*6*_): *δ* 165.78, 158.86, 153.40, 152.43, 143.08, 142.97, 137.60, 136.52, 131.23, 116.61, 112.64, 106.34, 104.16, 82.78, 67.51, 66.80, 66.57, 51.69, 46.57, 27.66 ppm. MS (ESI+) *m/z:* 455.03 [M + H]^+^. HRMS *m/z:* [M + H]^+^ calcd for C_23_H_27_N_4_O_6_, 455.19251; found 455.19082. HPLC: t_R_ = 16.695, purity ≥ 95%.

#### Synthesis of *tert-*Butyl 5-cyano-7,10-dioxa-13,17,18,21-tetraazatetracyclo[12.5.2.1^2^,^6^.0^17^,^20^]docosa-1(20),2,4,6(22),14(21),15,18-heptaene-13-carboxylate (27)

The title compound was synthesized according to the procedure of (**26**) using *tert-*butyl *N*-[3-(4-cyano-3-hydroxyphenyl)pyrazolo[1,5-*a*]pyrimidin-5-yl]-*N*-[2-(2-hydroxyethoxy)ethyl]carbamate (**23**) (0.27 g, 1.82 mmol). Purification by silica gel column chromatography with a mobile phase of *n-*hexane and ethyl acetate (ratio gradually ranging from 5:1 to 1:1) afforded the desired compound as a yellowish solid (0.13 g, 52%). ^1^H NMR (500 MHz, DMSO-*d*_*6*_): *δ* 9.03 (d, *J* = 7.8 Hz, 1H, Het*H*), 8.84 (d, *J* = 1.5 Hz, 1H, Ph*H*), 8.75 (s, 1H, Het*H*), 7.69 (d, *J* = 8.1 Hz, 1H, Ph*H*), 7.55 (d, *J* = 7.7 Hz, 1H, Het*H*), 7.49 (dd, *J* = 8.1, 1.4 Hz, 1H, Ph*H*), 4.49 (t, *J* = 5.7 Hz, 2H, C*H*_2_), 4.07 (t, *J* = 6.5 Hz, 2H, C*H*_2_), 3.92 (dt, *J* = 11.0, 6.0 Hz, 4H, C*H*_2_), 1.54 (s, 9H, (C*H*_3_)_3_) ppm.. ^13^C NMR (126 MHz, DMSO-*d*_*6*_): 161.14, 153.67, 152.38, 143.31, 143.22, 138.75, 136.62, 133.38, 117.47, 116.98, 111.59, 106.08, 104.46, 96.75, 82.86, 82.86, 68.13, 66.96, 46.87, 30.42, 27.67 ppm. MS (ESI+) *m/z:* 422.17 [M + H]^+^. HRMS *m/z:* [M + H]^+^ calcd for C_22_H_24_N_5_O_4_, 422.18228; found 422.18203. HPLC: t_R_ = 17.198, purity ≥ 95%.

#### Synthesis of *tert-*Butyl 4-cyano-7,10-dioxa-13,17,18,21-tetraazatetracyclo[12.5.2.1^2^,^6^.0^17^,^20^]docosa-1(20),2,4,6(22),14(21),15,18-heptaene-13-carboxylate (28)

The desired compound was synthesized according to the procedure of (**26**) using *tert-*butyl *N*-[3-(3-cyano-5-hydroxyphenyl)pyrazolo[1,5-*a*]pyrimidin-5-yl]-*N*-[2-(2-hydroxyethoxy)ethyl]carbamate (**24**) (0.27 g, 0.61 mmol). Purification was done by silica gel column chromatography with a mobile phase of *n-*hexane and ethyl acetate (ratio gradually ranging from 1:1 to 1:2). The title compound was yielded as a yellowish solid (0.17 g, 66%). ^1^H NMR (500 MHz, DMSO-*d*_*6*_): *δ* 9.01 (d, *J* = 7.8 Hz, 1H, Het*H*), 8.98 (dd, *J* = 2.6, 1.4 Hz, 1H, Ph*H*), 8.77 (s, 1H, Het*H*), 7.84 (t, *J* = 1.4 Hz, 1H, Ph*H*), 7.53 (d, *J* = 7.8 Hz, 1H, Het*H*), 7.15 (dd, *J* = 2.6, 1.3 Hz, 1H, Ph*H*), 4.38 (t, *J* = 5.7 Hz, 2H, C*H*_2_), 4.07 (t, *J* = 6.4 Hz, 2H, C*H*_2_), 3.91 (t, *J* = 6.4 Hz, 2H, C*H*_2_), 3.85 (t, *J* = 5.7 Hz, 2H, C*H*_2_), 1.54 (s, 9H, (C*H*_3_)_3_) ppm. 13C NMR (126 MHz, DMSO-*d*_*6*_): *δ* 159.57, 153.40, 152.41, 142.98, 142.91, 136.49, 135.07, 120.40, 118.82, 117.60, 117.60, 116.26, 112.19, 105.64, 104.25, 82.79, 67.58, 66.98, 46.83, 27.67 ppm. MS (ESI+) *m/z:* 444.08 [M + Na]^+^. HRMS *m/z:* [M + H]^+^ calcd for C_22_H_24_N_5_O_4_, 422.18228; found 422.18204. HPLC: t_R_ = 17.714, purity ≥ 95%.

#### Synthesis of *tert*-Butyl 7,10-dioxa-13,17,18,21-tetraazatetracyclo[12.5.2.1^2^,^6^.0^17^,^20^]docosa-1(20),2,4,6(22),14(21),15,18-heptaene-13-carboxylate (29)

The desired compound was synthesized according to the procedure of (**26**) using *tert-*butyl *N*-[2-(2-hydroxyethoxy)ethyl]-*N*-[3-(3-hydroxyphenyl)pyrazolo[1,5-*a*]pyrimidin-5-yl]carbamate (**25**) (90 mg, 0.22 mmol). Purification by silica gel column chromatography with a mobile phase of *n-*hexane and ethyl acetate (ratio gradually ranging from 5:1 to 1:1) led to the desired compound as a bright yellow solid (51 mg, 59%). ^1^H NMR (500 MHz, DMSO-*d*_*6*_): *δ* 8.97 (d, *J* = 7.8 Hz, 1H, Het*H*), 8.67 – 8.57 (m, 2H, Het*H*, Ph*H*), 7.51 (d, *J* = 7.8 Hz, 1H, Het*H*), 7.33 (d, *J* = 7.6 Hz, 1H Ph*H*), 7.25 (t, *J* = 7.9 Hz, 1H, Ph*H*), 6.69 (dd, *J* = 8.3, 2.7 Hz, 1H, Ph*H*), 4.31 (t, *J* = 6.3 Hz, 2H, C*H*_2_), 4.08 (t, *J* = 6.9 Hz, 2H, C*H*_2_), 3.92 (t, *J* = 6.9 Hz, 2H, C*H*_2_), 3.87 (t, *J* = 6.3 Hz, 2H, C*H*_2_), 1.54 (s, 9H, (C*H*_3_)_3_) ppm. ^13^C NMR (126 MHz, DMSO-*d*_*6*_): *δ* 158.94, 152.70, 152.46, 142.43, 136.29, 133.32, 129.38, 116.92, 114.41, 110.88, 107.29, 103.62, 82.64, 66.15, 65.84, 46.18, 27.66 ppm. MS (ESI+) *m/z:* 397.20 [M + H]^+^. HRMS *m/z:* [M + H]^+^ calcd for C_21_H_25_N_4_O_4_, 397.18703; found 397.18689. HPLC: t_R_ = 18.047, purity ≥ 95%.

#### Synthesis of Methyl 2-hydroxy-4-(5-{[2-(2-hydroxyethoxy)ethyl]amino}pyrazolo[1,5-*a*]pyrimidin-3-yl)benzoate (30)

Methyl 4-(5-{[(*tert*-butoxy)carbonyl][2-(2-hydroxyethoxy)ethyl]amino}pyrazolo[1,5-*a*]pyrimidin-3-yl)-2-hydroxybenzoate (**22**) was suspended in methylene chloride (2.5 mL) and trifluoroacetic acid (0.75 mg, 6.50 mmol) was added slowly at 0 °C. After stirring for 16 h at room temperature the solvent was evaporated under vacuum. The crude product was purified by silica gel column chromatography with a mobile phase of *n-*hexane and ethyl acetate (ratio gradually ranging from 1:0 to 0:1). The white solid obtained was the desired compound (50 mg g, 85%). ^1^H NMR (500 MHz, DMSO-*d*_*6*_): *δ* 10.62 (s, 1H, O*H*), 8.52 (d, *J* = 7.6 Hz, 1H, Het*H*), 8.44 (s, 1H, Het*H*), 7.88 (t, *J* = 5.3 Hz, 1H, N*H*), 7.78 – 7.75 (m, 2H, Ph*H*), 7.66 (dd, *J* = 8.4, 1.6 Hz, 1H, Ph*H*), 6.39 (d, *J* = 7.6 Hz, 1H, Het*H*), 3.89 (s, 3H, OC*H*_3_), 3.72 – 3.49 (m, 9H, (C*H*_3_)_3_) ppm. ^13^C NMR (126 MHz, DMSO-*d*_*6*_): *δ* 169.63, 160.93, 157.89, 156.33, 145.58, 142.43, 141.34, 135.58, 129.88, 115.81, 111.74, 108.03, 103.05, 100.52, 72.25, 68.41, 60.23, 52.25, 40.51 ppm. MS (ESI+) *m/z:* 372.96 [M + H]^+^. HRMS *m/z:* [M + H]^+^ calcd for C_18_H_21_N_4_O_5_, 373.15065; found 373.15150. HPLC: t_R_ = 12.942, purity ≥ 95%.

#### Synthesis of 4-(5-{[(*tert*-Butoxy)carbonyl][2-(2-hydroxyethoxy)ethyl]amino}pyrazolo[1,5-*a*]pyrimidin-3-yl)-2-hydroxybenzoic acid (31)

A solution of lithium hydroxide monohydrate (31 mg, 0.74 mmol) in water (1 mL) was added to a solution of methyl 4-(5-{[(*tert*-butoxy)carbonyl][2-(2-hydroxyethoxy)ethyl]amino}pyrazolo[1,5-*a*]pyrimidin-3-yl)-2-hydroxybenzoate (**22**) (70 mg, 0.15 mmol) in tetrahydrofuran (5 mL) and it was stirred for 16 h at 50 °C. The solvent was removed in vacuo and the residue was taken up with water. Afterwards the solution was acidified with 1N HCl and it was extracted with ethyl acetate. The organic layers were dried over MgSO_4_ and the solvent was evaporated. The crude product was purified by silica gel column chromatography with a mobile phase of petroleum ether and tetrahydrofuran with the addition of 1% acetic acid (ratio of 1:2). The white solid (29 mg, 42%) was the title compound. ^1^H NMR (500 MHz, DMSO-*d*_*6*_): *δ* 11.40 (s, 1H, O*H*), 9.00 (d, *J* = 7.7 Hz, 1H, Het*H*), 8.76 (s, 1H, Het*H*), 7.80 (d, *J* = 8.2 Hz, 1H, Ph*H*), 7.69 – 7.66 (m, 2H, Ph*H*), 7.49 (d, *J* = 7.7 Hz, 1H, Het*H*), 4.54 (s, 1H, O*H*), 4.22 (t, *J* = 6.4 Hz, 2H, C*H*_2_), 3.78 (t, *J* = 6.4 Hz, 2H, C*H*_2_), 3.49 – 3.43 (m, 4H, C*H*_2_), 1.53 (s, 9H, (C*H*_3_)_3_) ppm. ^13^C NMR (126 MHz, DMSO-*d*_*6*_): *δ* 171.90, 161.71, 153.79, 152.83, 143.79, 143.04, 139.53, 136.58, 130.51, 116.07, 112.49, 106.40, 104.75, 82.54, 72.27, 67.98, 60.23, 45.83, 31.16, 29.83, 27.67 ppm. MS (ESI+) *m/z:* 458.98 [M + H]^+^. HRMS *m/z:* [M + H]^+^ calcd for C_22_H_27_N_4_O_7_, 459.18743; found 459.18597. HPLC: t_R_ = 14.985, purity ≥ 95%.

#### Synthesis of 13-[(*tert*-Butoxy)carbonyl]-7,10-dioxa-13,17,18,21-tetraazatetracyclo[12.5.2.1^2^,^6^.0^17^,^20^]docosa-1(20),2,4,6(22),14(21),15,18-heptaene-5-carboxylic acid (32)

The desired compound was synthesized according to the procedure of (**31**) using 13-*tert*-butyl 5-methyl 7,10-dioxa-13,17,18,21-tetraazatetracyclo[12.5.2.1^2^,6.0^1^7,^20^]docosa-1(20),2,4,6(22),14(21), 15,18-heptaene-5,13-dicarboxylate (**26**) (68 mg, 0.15 mmol). Purification was done by silica gel column chromatography with a mobile phase of ethyl acetate with 1% AcOH. The title compound was obtained as a white solid (23 mg, 35%). ^1^H NMR (500 MHz, DMSO-*d*_*6*_): *δ* 12.35 (s, 1H, COO*H*), 9.01 (d, *J* = 7.8 Hz, 1H, Het*H*), 8.75 (d, *J* = 1.5 Hz, 1H, Het*H*), 8.72 (s, 1H, Ph*H*), 7.70 (d, *J* = 8.1 Hz, 1H, Ph*H*), 7.55 (d, *J* = 7.8 Hz, 1H, Het*H*), 7.42 (dd, *J* = 8.1, 1.5 Hz, 1H, Ph*H*), 4.40 (t, *J* = 6.0 Hz, 2H, C*H*_2_), 4.10 (t, *J* = 6.6 Hz, 2H, C*H*_2_), 3.93 (t, *J* = 6.6 Hz, 2H, C*H*_2_), 3.89 (t, *J* = 6.0 Hz, 2H, C*H*_2_), 1.54 (s, 9H, (C*H*_3_)_3_) ppm. ^13^C NMR (126 MHz, DMSO-*d*_*6*_): *δ* 166.83, 158.82, 153.36, 152.45, 143.03, 142.93, 137.23, 136.50, 131.42, 117.64, 116.61, 112.61, 106.46, 104.15, 82.77, 67.51, 66.86, 66.58, 46.60, 27.67 ppm. MS (ESI+) *m/z:* 441.00 [M + H]^+^. HRMS *m/z:* [M + H]^+^ calcd for C_22_H_25_N_4_O_6_, 441.17686; found 441.17510. HPLC: t_R_ = 15.137, purity ≥ 95%.

#### Synthesis of Methyl 7,10-dioxa-13,17,18,21-tetraazatetracyclo[12.5.2.1^2^,^6^.0^17^,^20^]docosa-1(20),2,4,6(22),14(21),15,18-heptaene-5-carboxylate (33)

To a 0 °C cooled solution of 13-*tert*-butyl 5-methyl 7,10-dioxa-13,17,18,21-tetraazatetracyclo[12.5.2.1^2^,6.0^1^7,^20^]docosa-1(20),2,4,6(22),14(21),15,18-heptaene-5,13-dicarboxylate (**26**) (0.30 g, 0.66 mmol) in methylene chloride (12 mL) trifluoroacetic acid (3.58 g, 31.36 mmol) was added dropwise and it was stirred for 16 h at room temperature. Afterwards the solvent was removed under reduced pressure and the residue was taken up in methanol. Potassium carbonate was added to the stirring solution. Subsequently, the solution was filtrated and the solvent was removed again. The residue was taken up with ethyl acetate and it was washed with water and brine. After drying over MgSO_4_ and removal of the organic solvent under vacuum, the title compound was yielded as a white solid (0.23 g, 99%). ^1^H NMR (500 MHz, DMSO-*d*_*6*_): *δ* 8.83 (d, *J* = 1.5 Hz, 1H, Het*H*), 8.57 (d, *J* = 7.6 Hz, 1H, Het*H*), 8.39 (s, 1H, Ph*H*), 7.94 (t, *J* = 5.4 Hz, 1H, N*H*), 7.70 (d, *J* = 8.1 Hz, 1H, Ph*H*), 7.29 (dd, *J* = 8.1, 1.4 Hz, 1H, Ph*H*), 6.34 (d, *J* = 7.6 Hz, 1H, Het*H*), 4.37 (t, *J* = 7.9 Hz, 2H, C*H*_2_), 4.00 – 3.97 (m, 2H, C*H*_2_), 3.89 – 3.86 (m, 2H, C*H*_2_), 3.76 (s, 3H, OC*H*_3_), 3.56 – 3.50 (m, 2H, C*H*_2_) ppm. ^13^C NMR (126 MHz, DMSO-*d*_*6*_): *δ* 165.66, 158.08, 156.18, 145.33, 141.81, 139.04, 135.82, 131.61, 115.64, 114.82, 110.45, 103.40, 100.21, 65.35, 65.27, 51.57 ppm. MS (ESI+) *m/z:* 354.96 [M + H]^+^. HRMS *m/z:* [M + H]^+^ calcd for C_18_H_19_N_4_O_4_, 355.14008; found 355.14078. HPLC: t_R_ = 12.869, purity ≥ 95%.

#### Synthesis of 7,10-Dioxa-13,17,18,21-tetraazatetracyclo[12.5.2.1^2^,^6^.0^17^,^20^]docosa-1(20),2,4,6(22),14(21),15,18-heptaene-5-carboxylic acid (34)

In tetrahydrofuran (3 mL) methyl 7,10-dioxa-13,17,18,21-tetraazatetracyclo[12.5.2.1^2^,6.0^1^7,^20^]docosa-1(20),2,4,6(22),14(21),15,18-heptaene-5-carboxylate (**33**) (50 mg, 0.14 mmol) and lithium hydroxide monohydrate (30 mg, 0.71 mmol) were dissolved and water (0.6 mL) was added. The reaction mix was stirred at 50 °C for 16 h. After that, the solvent was removed under reduced pressure and the residue was taken up in water. By adding 10% hydrochloric acid pH 1 was set and the precipitated solid was filtrated. Washing and drying of the filtrate resulted in the white solid tile compound (47 mg, 99%). ^1^H NMR (500 MHz, DMSO-*d*_*6*_): *δ* 12.24 (bs, 1H, COO*H*), 8.81 (s, 1H, Het*H*), 8.57 (d, *J* = 7.6 Hz, 1H, Het*H*), 8.38 (s, 1H, Ph*H*), 7.93 (t, *J* = 5.4 Hz, 1H, N*H*), 7.70 (d, *J* = 8.1 Hz, Ph*H*), 7.27 (dd, *J* = 8.1, 1.4 Hz, 1H, Ph*H*), 6.34 (d, *J* = 7.6 Hz, 1H, Het*H*), 4.38 (t, *J* = 7.1 Hz, 2H, C*H*_2_), 3.99 (t, *J* = 7.5 Hz, 2H, C*H*_2_), 3.89 – 3.86 (m, 2H, C*H*_2_), 3.56 – 3.50 (m, 2H, C*H*_2_) ppm. ^13^C NMR (126 MHz, DMSO-*d*_*6*_): *δ* 166.70, 158.07, 156.14, 145.27, 141.76, 138.69, 135.80, 131.82, 115.79, 115.61, 110.39, 103.50, 100.18, 65.31, 65.22, 63.89, 59.77 ppm. MS (ESI+) *m/z:* 340.93 [M + H]^+^. HRMS *m/z:* [M + H]^+^ calcd for C_17_H_17_N_4_O_4_, 341.12443; found 341.12573. HPLC: t_R_ = 11.408, purity ≥ 95%.

#### Synthesis of Methyl 13-acetyl-7,10-dioxa-13,17,18,21-tetraazatetracyclo[12.5.2.1^2^,^6^.0^17^,^20^]docosa-1(20),2,4,6(22),14(21),15,18-heptaene-5-carboxylate (35)

Acetyl chloride (138 mg, 1.76 mmol) and *N,N*-diisopropylethylamine (227 mg, 1.76 mmol) were added to a solution of methyl 7,10-dioxa-13,17,18,21-tetraazatetracyclo[12.5.2.1^2^,6.0^1^7,^20^]docosa-1(20),2,4,6(22),14(21),15,18-heptaene-5-carboxylate (**33**) (50 mg, 0.14 mmol) in acetonitrile (2 mL) under argon atmosphere. Reaction took place at room temperature for 2 h. Afterwards the solvent was removed under reduced pressure and purification was done by silica gel column chromatography with a mobile phase of *n-*hexane and ethyl acetate (ratio gradually ranging from 1:1 to 0:1). The desired product was obtained as a brownish solid (42 mg, 76%). ^1^H NMR (500 MHz, DMSO-*d*_*6*_): *δ* 9.06 (d, *J* = 7.7 Hz, 1H, Het*H*), 8.82 (d, *J* = 1.0 Hz, 1H, Ph*H*), 8.77 (s, 1H, Het*H*), 7.70 (d, *J* = 8.1 Hz, 1H, Ph*H*), 7.56 (d, *J* = 7.7 Hz, 1H, Het*H*), 7.46 (dd, *J* = 8.2, 1.4 Hz, 1H, Ph*H*), 4.40 (t, *J* = 5.6 Hz, 2H, C*H*_2_), 4.16 (t, *J* = 6.7 Hz, 2H, C*H*_2_), 4.00 (t, *J* = 6.7 Hz, 2H, C*H*_2_), 3.88 (t, *J* = 5.6 Hz, 2H, C*H*_2_), 3.78 (s, 3H, C*H*_3_), 2.42 (s, 3H, C*H*_3_) ppm. ^13^C NMR (126 MHz, DMSO-*d*_*6*_): *δ* 171.76, 165.81, 159.10, 153.75, 143.17, 143.04, 137.53, 136.59, 131.15, 116.88, 116.76, 113.14, 106.64, 105.25, 68.14, 66.87, 59.77, 51.72, 47.62, 24.33 ppm. MS (ESI+) *m/z:* 396.95 [M + H]^+^. HRMS *m/z:* [M + H]^+^ calcd for C_20_H_21_N_4_O_5_, 397.15065; found 397.14991. HPLC: t_R_ = 16.955, purity ≥ 95%.

#### Synthesis of Methyl 13-(2-methylpropanoyl)-7,10-dioxa-13,17,18,21-tetraazatetracyclo[12.5.2.1^2^,^6^.0^17^,^20^]docosa-1(20),2,4,6(22),14(21),15,18-heptaene-5-carboxylate (36)

The title compound was synthesized according to the procedure of (**35**) using isobutyryl chloride (60 mg, 0.56 mmol). Purification by silica gel column chromatography with a mobile phase of *n-*hexane and ethyl acetate (ratio of 1:1) led to the desired compound as a yellow oil (50 mg, 83%). ^1^H NMR (500 MHz, DMSO-*d*_*6*_): *δ* 9.05 (d, *J* = 7.6 Hz, 1H, Het*H*), 8.85 (d, *J* = 1.5 Hz, 1H, Ph*H*), 8.78 (s, 1H, Het*H*), 7.70 (d, *J* = 8.1 Hz, 1H, Ph*H*), 7.47 (dd, *J* = 8.2, 1.5 Hz, 1H, Ph*H*), 7.39 (d, *J* = 7.7 Hz, 1H, Het*H*), 4.40 (t, *J* = 5.5 Hz, 2H, C*H*_2_), 4.18 (t, *J* = 6.5 Hz, 2H, C*H*_2_), 3.97 (t, *J* = 6.5 Hz, 2H, C*H*_2_), 3.88 (t, *J* = 5.5 Hz, 2H, C*H*_2_), 3.78 (s, 3H, C*H*_3_), 3.16 (hept, *J* = 6.7 Hz, 1H, C*H*), 1.16 (d, *J* = 6.6 Hz, 6H, (C*H*_3_)_2_) ppm. ^13^C NMR (126 MHz, DMSO-*d*_*6*_): *δ* 178.57, 165.82, 159.14, 154.24, 143.16, 137.50, 136.53, 131.14, 116.91, 116.78, 113.26, 106.70, 105.62, 68.34, 67.88, 67.65, 51.71, 47.19, 32.29, 19.60 ppm. MS (ESI+) *m/z:* 424.99 [M + H]^+^. HRMS *m/z:* [M + H]^+^ calcd for C_22_H_25_N_4_O_5_, 425.18195; found 425.18032. HPLC: t_R_ = 14.710, purity ≥ 95%.

#### Synthesis of *N*-Benzyl-7,10-dioxa-13,17,18,21-tetraazatetracyclo[12.5.2.1^2^,^6^.0^17^,^20^]docosa-1(20),2,4,6(22),14(21),15,18-heptaene-5-carboxamide (37)

A solution of 7,10-dioxa-13,17,18,21-tetraazatetracyclo[12.5.2.1^2^,6.0^1^7,^20^]docosa-1(20),2,4,6(22),14(21),15,18-heptaene-5-carboxylic acid (**34**) (127 mg, 0.37 mmol) and *N*-(3-dimethylaminopropyl)-*N*′-ethylcarbodiimide hydrochloride (104 mg, 0.67 mmol) in methylene chloride (25 mL) was stirred at room temperature. After 1 h benzylamine (72 mg, 0.67 mmol) was added and the mixture was kept on stirring for 16 h at room temperature. The reaction was quenched by the addition of water and it was acidified with 1N HCl. The product was extracted with ethyl acetate. The combined organic layers were dried over MgSO_4_, filtrated and the solvent was evaporated under reduced pressure. The residue was recrystallized with water to obtain the pure product as a white solid (56 mg, 35%). ^1^H NMR (500 MHz, DMSO-*d*_*6*_): *δ* 8.82 (d, *J* = 0.9 Hz, 1H, Ph*H*), 8.64 (t, *J* = 6.1 Hz, 1H, N*H*), 8.57 (d, *J* = 7.6 Hz, 1H, Het*H*), 8.38 (s, 1H, Het*H*), 7.91 (t, *J* = 5.4 Hz, 1H, N*H*), 7.83 (d, *J* = 8.1 Hz, 1H, Ph*H*), 7.35 – 7.31 (m, 5H, Ph*H*), 7.27 – 7.21 (m, 1H, Ph*H*), 6.34 (d, *J* = 7.6 Hz, 1H, Het*H*), 4.52 (d, *J* = 6.1 Hz, 2H, C*H*_2_), 4.48 (t, *J* = 7.9 Hz, 2H, C*H*_2_), 4.05 – 4.01 (m, 2H, C*H*_2_), 3.90 – 3.87 (m, 2H, C*H*_2_), 3.54 (q, *J* = 6.7 Hz, 2H, C*H*_2_) ppm. ^13^C NMR (126 MHz, DMSO-*d*_*6*_): *δ* 164.68, 156.50, 156.09, 145.19, 141.66, 139.99, 137.63, 135.79, 131.10, 128.26, 127.01, 126.57, 117.92, 116.19, 110.02, 103.57, 100.14, 65.46, 65.37, 64.03, 42.57 ppm. MS (ESI+) *m/z:* 430.00 [M + H]^+^. HRMS *m/z:* [M + H]^+^ calcd for C_24_H_24_N_5_O_3_, 430.18737; found 430.18709. HPLC: t_R_ = 13.794, purity ≥ 95%.

## ACKNOWLEDGMENT

The authors are grateful for support by the SGC, a registered charity (number 1097737) that receives funds from AbbVie, Bayer Pharma AG, Boehringer Ingelheim, Canada Foundation for Innovation, Eshelman Institute for Innovation, Genome Canada, Innovative Medicines Initiative EUbOPEN (No 875510), Janssen, Merck KGaA Darmstadt Germany, MSD, Novartis Pharma AG, Ontario Ministry of Economic Development and Innovation, Pfizer, São Paulo Research Foundation-FAPESP, Takeda, and Wellcome. S.K. and C.K. are grateful for support by the German cancer network (DKTK) and the Frankfurt cancer center (FCI). We also thank the beamline scientists at SLS (CH) for their great support during data collection.

## SCHEMES

**Scheme 1.**
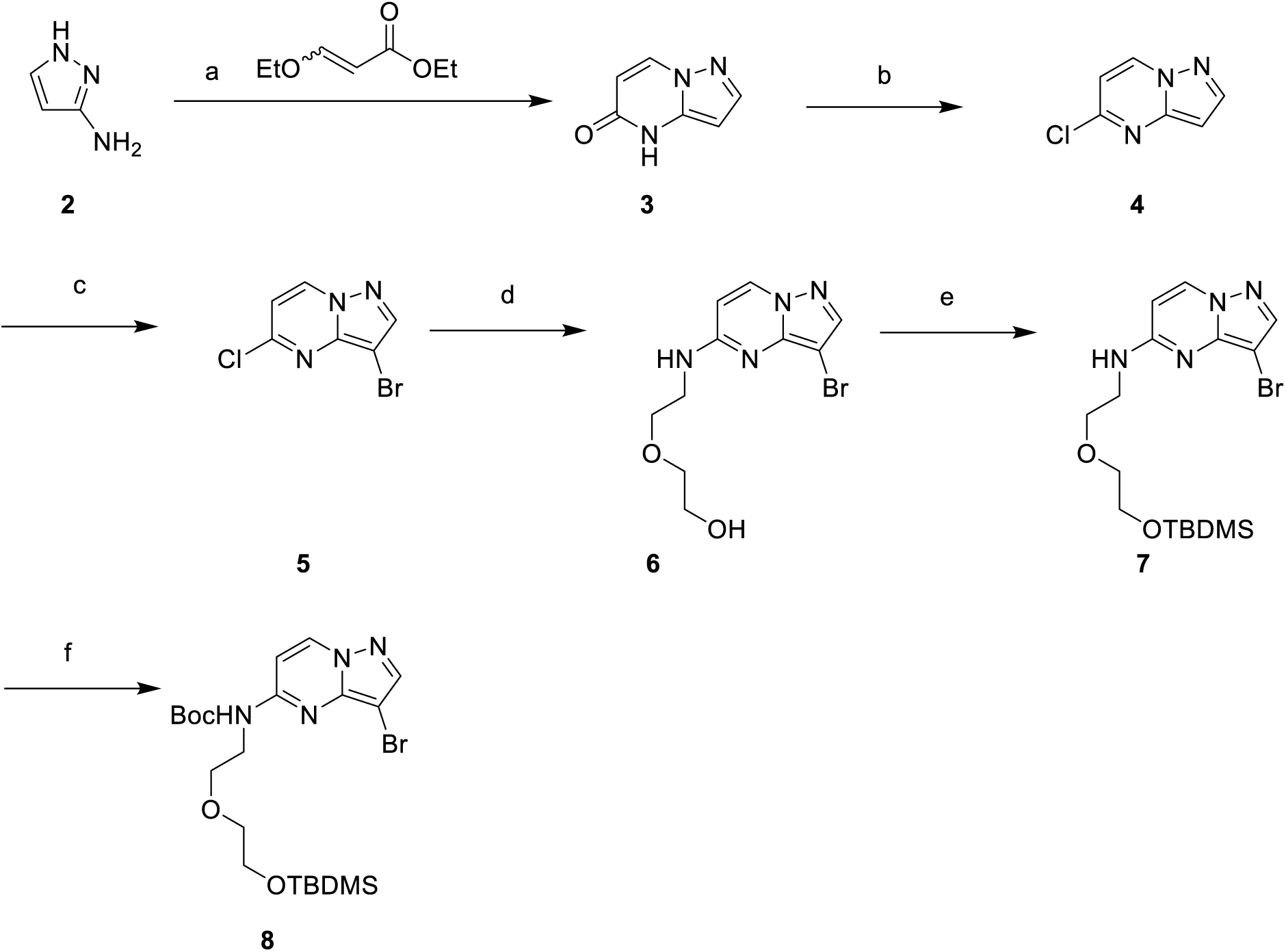
Synthesis of pyrazolo[1,5-a]pyrimidine (hinge-binder) with double protected ether-linker^a^. ^a^Reagents and conditions: (a) Cs_2_CO_3_, DMF, 4 h, 110 °C; (b) POCl_3_, 2 h, 120 °C; (c) NBS, DMF, rt; (d) 2-(2-aminoethoxy)ethanol, DIPEA, ACN, 1 h, reflux; (e) TBDMSCl, TEA, DMF, 1 h, rt; (f) Boc_2_O, TEA, 4-DMAP, THF, 3 h, reflux.

**Scheme 2.**
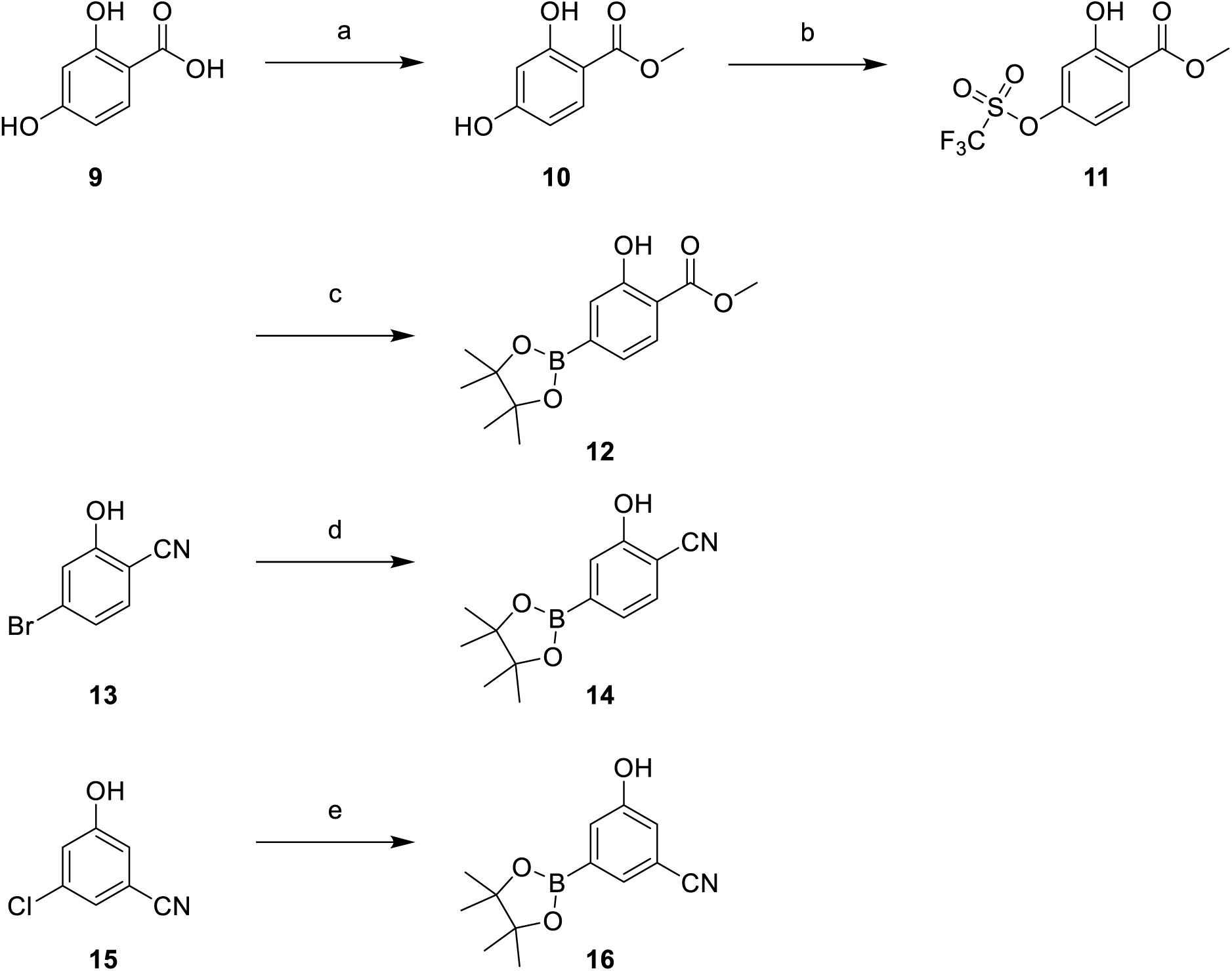
Synthesis of pinacol boronic esters (Compound 12, 14 and 16)^a^. ^a^Reagents and conditions: (a) MeOH, H_2_SO_4_, 16 h, reflux; (b) Tf_2_O, pyridine, 4-DMAP, DCM, 16 h, 0 °C to rt; (c) B_2_pin_2_, Pd(dppf)Cl_2_·DCM, KOAc, 1,4-dioxane, 3.5 h, 100 °C; (d) B_2_pin_2_, Pd(dppf)Cl_2_, KOAc, 1,4-dioxane, 3.5 h, 100 °C; (e) B_2_pin_2_, Pd_2_(dba)_3_, P(Cy)_3_, KOAc, dimethoxymethane, 2 h, 150 °C, MW.

**Scheme 3.**
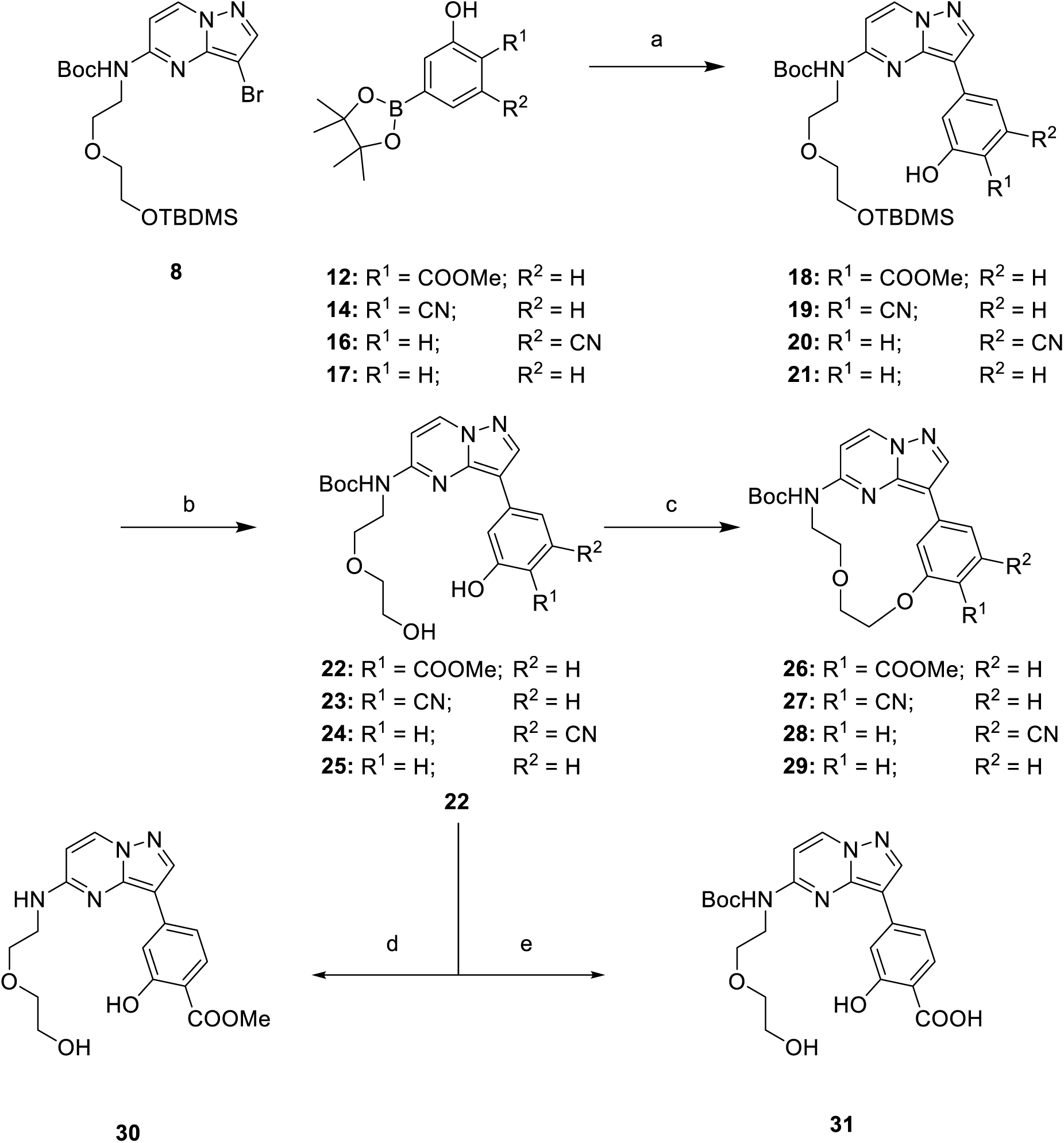
Synthesis of macrocyclic compounds (26–29)^a^. ^a^Reagents and conditions: (a) Pd_2_(dba)_3_, XPhos, K_3_PO_4_, 1,4-dioxane, H_2_O, 40 min, 110 °C; (b) 1 M TBAF, THF, 3 h, rt; (c) DIAD, TPP, Na_2_SO_4_, toluene, 2-MTHF, 3 h, 90 °C; (d) TFA, DCM, 16 h, 0 °C to rt; (e) LiOH·H_2_O, H_2_O, THF, 16 h, 50 °C.

**Scheme 4.**
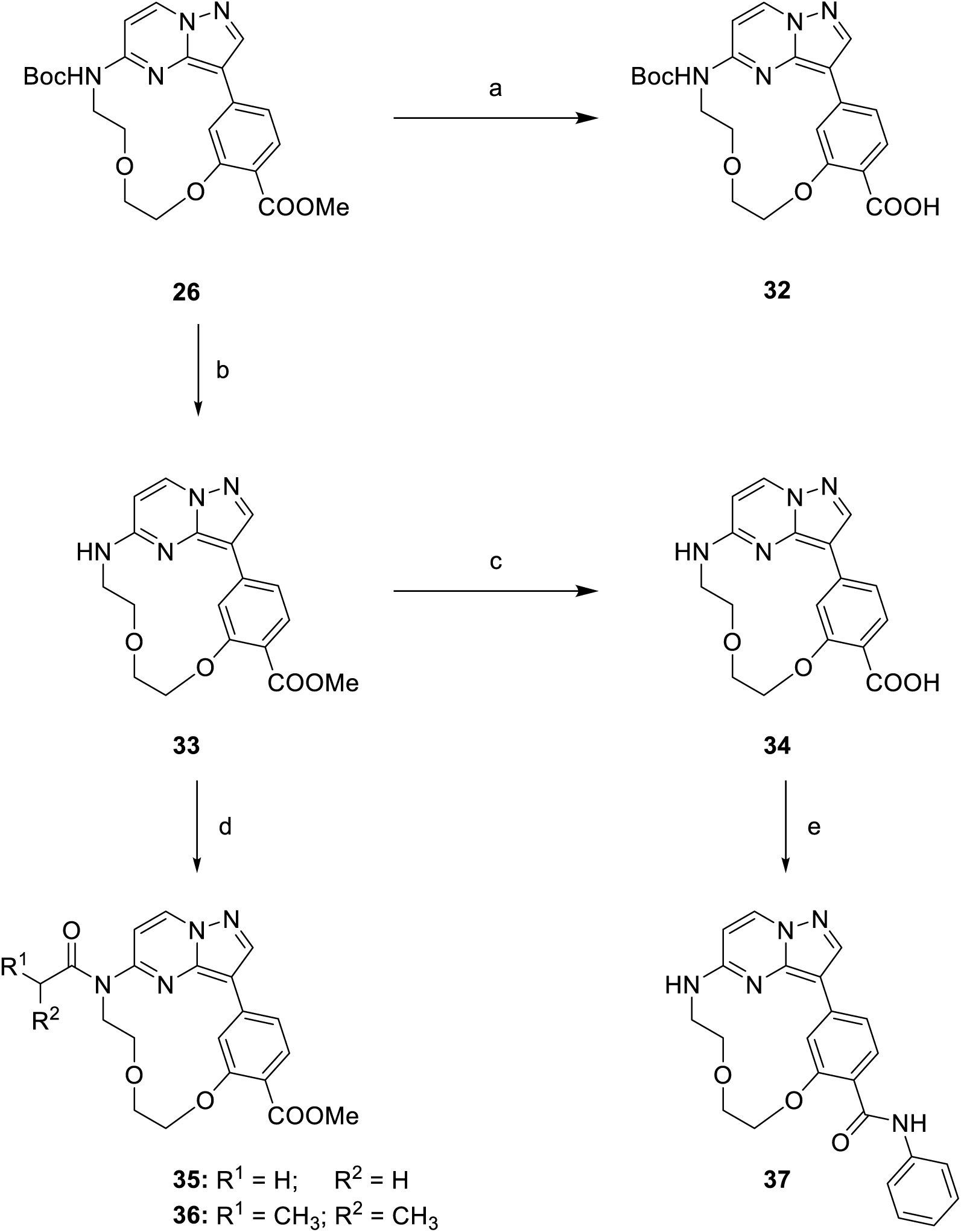
Synthesis of macrocyclic compounds (32–37)^a^. ^a^Reagents and conditions: (a) LiOH·H_2_O, H_2_O, THF, 16 h, 50 °C; (b) TFA, DCM, 16 h, 0 °C to rt; (c) LiOH·H_2_O, H_2_O, THF, 16 h, 50 °C; (d) acetyl chloride or isobutyryl chloride, DIPEA, ACN, 2 h, rt; (e) benzylamine EDC·HCl, DCM, 16 h, rt.

## Abbreviations

2-MTHF: 2-methyltetrahydrofuran
ATP: Adenosine triphosphate
B_2_pin_2_: Bis(pinacolato)diboron
CK2: Casein kinase 2
DIAD: diisopropyl azodicarboxylate
DIPEA: *N,N*-diisopropylethylamine
HEPES: (4-(2-hydroxyethyl)-1-piperazineethanesulfonic acid
IPTG: Isopropyl-thio-galactopyranoside
MES: 2-(*N*-morpholino)ethanesulfonic acid
MGC: mammalian gene collection
PEG MME: polyethylene glycol monomethyl ether
P(Cy)_3_: Tricyclohexylphosphine
TCEP: Tris(2-carboxyethyl)phosphine
TEV: Tobacco Etch Virus
Tf_2_O: trifluoromethanesulfonic anhydride
TPP triphenylphosphine;
XPhos: 2-dicyclohexylphosphino-2′,4′,6′-triisopropylbiphenyl

