## Supporting Info for "Optimization of pyrazolo[1,5-a]pyrimidines lead to the identification of a highly selective casein kinase 2 inhibitor"

[&]German Translational Cancer Network (DKTK) site Frankfurt/Mainz.

[#]Frankfurt Cancer Institute (FCI), Paul-Ehrlich-Straße 42-44, 60596 Frankfurt am Main.

### I Supplementary Figures and Tables

Table S1: Differential scanning fluorimetry assay of compound 22–37.

| | 22 | 23 | 24 | 25 | 30 | 31 | 26 | 27 | 28 | 29 | 32 | 33 | 34 | 35 | 36 | 37 |
|---|---|---|---|---|---|---|---|---|---|---|---|---|---|---|---|---|
| **AAK1** | 2,3 | 3,6 | 2,1 | n.d. | 3,8 | 3,2 | 1,7 | n.d. | n.d. | n.d. | 3,9 | 7,9 | 9,3 | 4,1 | 3,2 | n.d. |
| **AurB** | 0,6 | 1,9 | 1,1 | n.d. | 1,6 | -2,8 | 0,8 | n.d. | n.d. | n.d. | -1,6 | 1 | 2,1 | 1,7 | 1 | n.d. |
| **BMX** | 1 | -0,3 | -0,3 | n.d. | 0,5 | 0,3 | 0,5 | n.d. | n.d. | n.d. | 0,5 | 0,9 | 0,1 | 0,1 | 0,1 | n.d. |
| **BRAF** | 0,3 | 7,2 | 5,3 | n.d. | -0,5 | 2,5 | 0 | n.d. | n.d. | n.d. | 2,8 | 1,8 | 1 | 0,6 | 0,5 | n.d. |
| **CaMK1g** | -0,7 | -0,1 | -0,4 | n.d. | -0,7 | -0,2 | -0,3 | n.d. | n.d. | n.d. | -0,3 | -0,5 | 0 | -0,5 | 0 | n.d. |
| **CaMKK2** | 0,8 | 4,7 | 0,5 | 1,6 | 3 | 5,5 | 1,6 | 6,7 | 0,9 | 2,5 | 6 | 5 | 8,7 | 2,5 | 1,9 | 7,2 |
| **CASK** | -0,3 | 0,3 | -0,3 | n.d. | -0,4 | -0,1 | 0 | n.d. | n.d. | n.d. | 0,3 | 0,2 | 0,1 | -0,1 | -0,1 | n.d. |
| **CDK2** | 0 | 1,7 | 0 | 0,9 | 0,7 | 2,6 | 0 | 2,1 | -0,2 | 0,5 | 2,4 | 5,3 | 7 | 1,7 | 0,6 | 1,5 |
| **CHK2** | 0,9 | 0,5 | 0,5 | 0,6 | 0,8 | 1,1 | 0,7 | 0,4 | 0,4 | 0,4 | 0,8 | 2,4 | 4,3 | 0,6 | 0,4 | 5,9 |
| **CK1d** | 0,3 | 0,6 | 0,6 | n.d. | 0,2 | 1,3 | 0,3 | n.d. | n.d. | n.d. | 1,6 | 0,7 | 2,5 | 0,5 | 0,4 | n.d. |
| **CK2α** | 0,1 | 7,4 | 2,2 | 2,2 | 0,4 | 15,1 | 0,4 | 3,8 | 0,4 | 0,3 | 11,7 | 2,3 | 10,6 | 0,8 | 1,5 | 0,8 |
| **CK2α`** | 0,8 | 7,6 | 3,4 | 2,1 | 1,2 | 15,7 | 1,6 | 4,4 | 1 | 0,7 | 11,8 | 3,5 | 11,3 | 1,7 | 2,4 | 2,7 |
| **CLK1** | 1,4 | 2,6 | 1 | 1,2 | 3,9 | 2,9 | 0,9 | 2,8 | 0,1 | 0,6 | 1,5 | 6,3 | 7,1 | 2,9 | 1,4 | 6,5 |
| **DAPK1** | 0,2 | 0,5 | 1 | 0,7 | 0,8 | 4,9 | 0,2 | 0,9 | 1 | 0,1 | 2 | 1,1 | 3,1 | 0 | 0,2 | 1,3 |
| **DAPK3** | 0,5 | 1,6 | 3 | 2 | 2,6 | 11,6 | -0,2 | 1,4 | 1 | 0,1 | 6,2 | 3,2 | 8,7 | 0,5 | 0,3 | 5,9 |
| **DRAK2** | 1,1 | 1,4 | 0,6 | 1 | 3,1 | 1,9 | 1,4 | 0,8 | -0,1 | 0,2 | 1,2 | 3,9 | 0,7 | 2,6 | 1,5 | 2,1 |
| **DYRK1A** | -1,1 | 0 | -1 | 0,6 | 0,5 | 1,7 | -1 | 0,6 | -0,7 | -0,3 | -0,1 | 0,2 | 0,6 | -1,1 | -1,2 | -1,1 |
| **EphA2** | 0,3 | 0,2 | 0,5 | n.d. | 0,2 | 0 | 0,3 | n.d. | n.d. | n.d. | 0,2 | 0,7 | 0,7 | 0,3 | 0,4 | n.d. |
| **Erk1** | -0,3 | n.d. | n.d. | n.d. | -0,3 | 0 | 0,6 | n.d. | n.d. | n.d. | 1 | -0,7 | 0,8 | -0,1 | -0,6 | n.d. |
| **Erk2** | -0,4 | 0,8 | -0,6 | n.d. | 0,5 | 0,5 | -0,5 | n.d. | n.d. | n.d. | 1 | 1,5 | -0,2 | -0,4 | 0 | n.d. |
| **Erk3** | 0,5 | n.d. | n.d. | n.d. | 0,9 | -0,1 | -0,2 | n.d. | n.d. | n.d. | -0,3 | 1,2 | 0,5 | 0,1 | 0,4 | n.d. |
| **FER** | 0 | n.d. | n.d. | n.d. | 0,4 | 0,8 | 0,1 | n.d. | n.d. | n.d. | 0,5 | 0,3 | -0,7 | -1,3 | 0,2 | n.d. |
| **FES** | -0,6 | -0,1 | -0,7 | -0,6 | -0,7 | -0,4 | -0,5 | -0,5 | -0,6 | -0,8 | -0,4 | 0,1 | 0 | -0,4 | -0,5 | -0,4 |
| **FGFR2** | -0,1 | 0,4 | 1,3 | 1 | 1,1 | 0,6 | -0,1 | 0 | 0,2 | 0,2 | 0,1 | 1,8 | 1,6 | 0,5 | 0 | 5,2 |
| **FLT1** | 0,3 | 1,2 | 4,6 | 2,9 | 1,8 | 2,3 | 0,4 | 1,5 | 1,6 | 1,1 | 1,2 | 3,4 | 3,5 | 1,6 | 0,8 | 3,9 |
| **GAK** | 1,3 | 3,1 | 3,3 | 2,5 | 3,9 | 4,6 | 1,2 | 3,5 | 2,1 | 1 | 4,5 | 6 | 9,1 | 3,6 | 3,7 | 7 |
| **GPRK5** | -0,2 | 0 | -0,1 | n.d. | 0 | 0,7 | -0,2 | n.d. | n.d. | n.d. | 0 | 0,8 | 0,8 | 0,1 | 0,2 | n.d. |
| **Haspin** | 0,4 | 2,2 | 1 | n.d. | 1,4 | 1 | 0,6 | n.d. | n.d. | n.d. | 0,9 | 3,4 | 2,9 | 1,9 | 1 | n.d. |
| **JNK1** | 0,4 | 0 | -0,3 | n.d. | 0,1 | 0,9 | 0,1 | n.d. | n.d. | n.d. | -0,2 | 0,4 | 0,3 | 0,4 | 0,5 | n.d. |
| **JNK2** | -0,3 | 0,5 | 0,2 | 0 | 0,7 | 0,6 | 0 | 0,3 | -0,1 | 0 | 0,3 | 1,1 | 0,3 | 0,2 | 0,2 | -0,1 |
| **LOK** | -0,3 | 0,3 | 2 | n.d. | 0 | 2,3 | -0,2 | n.d. | n.d. | n.d. | 0,8 | 3,4 | 6 | 1,1 | 1,2 | n.d. |
| **MAP2K2** | 1,6 | n.d. | n.d. | n.d. | 7,6 | 4,6 | 4,2 | n.d. | n.d. | n.d. | 2,5 | 6,4 | 7,6 | 6,1 | 7,3 | n.d. |
| **MAP2K4** | -0,1 | 1,7 | 0,6 | 1 | 2,3 | 1,6 | 0,5 | 1,1 | 0,3 | 1,1 | 2 | 3,7 | 6,5 | 1,6 | 1,6 | 5,4 |
| **MAP2K6** | -0,1 | 0,4 | -0,1 | 0,2 | 0,6 | 0,6 | 0,4 | 0,2 | 0,2 | 0 | 0,4 | 1,5 | 1,2 | 0,9 | 0,4 | 1,8 |
| **MAP2K7** | -0,4 | n.d. | n.d. | n.d. | 0,1 | 0,2 | 0 | n.d. | n.d. | n.d. | 0,4 | 0,4 | 1,8 | -0,6 | 0,2 | n.d. |
| **MAP3K5** | 0,1 | 0,7 | 0,4 | 0,1 | 0,7 | 2,4 | 0,5 | 3,4 | 0,3 | 0,2 | 1,6 | 3,6 | 5 | 0,5 | 0,4 | 5,1 |
| **MELK** | 2,8 | 4,1 | 1,8 | 2,5 | 4,6 | 1,1 | 1,6 | 3,7 | 1,1 | 0,8 | 0,1 | 6,6 | 5 | 3,4 | 3,2 | 6,9 |
| **MER** | -0,5 | -0,2 | 0,1 | n.d. | -0,3 | 0,8 | -0,6 | n.d. | n.d. | n.d. | -0,2 | -0,5 | 0 | -0,3 | 0,1 | n.d. |
| **MSK1_b** | 0,3 | 2,5 | 3,3 | 2,5 | 1,1 | 6 | -0,3 | 1,9 | 0 | 0,7 | 2,5 | 2,8 | 6,2 | 0,4 | 0,2 | 2,6 |
| **MST2** | 0,3 | 0,6 | 1 | n.d. | 1,1 | 0,2 | 0,5 | n.d. | n.d. | n.d. | 0,3 | 5,5 | 3,2 | 2,3 | 0,7 | n.d. |

| | | | | | | | | | | | | | | | | |
|---|---|---|---|---|---|---|---|---|---|---|---|---|---|---|---|---|
| **MYT1** | 0,9 | n.d. | n.d. | n.d. | 0,5 | -0,9 | -0,7 | n.d. | n.d. | n.d. | 1,1 | -0,8 | -1 | 0,8 | 0,9 | n.d. |
| **p38a** | -0,5 | 0 | -0,7 | -0,4 | 0,1 | 0 | -0,5 | -0,5 | -0,3 | -0,5 | 0 | 0,1 | 0,3 | -0,1 | -0,2 | -0,7 |
| **PAK1** | -0,4 | -0,2 | 0,2 | 1,6 | 0,7 | -1,4 | -0,8 | 0,2 | -0,5 | -0,4 | -0,9 | -1,7 | -1,5 | -0,6 | -0,5 | 0,9 |
| **PAK4** | 1,2 | 2,5 | 1,6 | n.d. | 1,1 | 1,1 | 1,3 | n.d. | n.d. | n.d. | 0,8 | 0,9 | 1,3 | 1,3 | 1,7 | n.d. |
| **PBK** | -0,4 | 0,1 | -0,4 | -0,4 | 0,1 | 0,3 | -0,5 | -0,1 | -0,2 | -0,4 | 0,2 | 0,4 | 0,7 | 0,2 | -0,2 | 0,9 |
| **PDHK4** | -0,1 | n.d. | n.d. | n.d. | 0,4 | 0 | -0,7 | n.d. | n.d. | n.d. | -0,9 | 0,1 | -0,4 | 0,3 | -0,2 | n.d. |
| **PHKg2** | 0,2 | 1,1 | -0,1 | n.d. | 0,6 | 0,7 | 0,2 | n.d. | n.d. | n.d. | 0,4 | 5 | 5,8 | 1,6 | 0,4 | n.d. |
| **PIM1** | -1,1 | -0,3 | -0,2 | n.d. | -0,9 | 4,3 | -0,3 | n.d. | n.d. | n.d. | 0,3 | 0,5 | 2,4 | 0,2 | -0,3 | n.d. |
| **PIM3** | 1,4 | n.d. | n.d. | n.d. | 2,8 | 5,7 | 1 | n.d. | n.d. | n.d. | 4 | 3,8 | 4,8 | 1,6 | 2,5 | n.d. |
| **PKG2** | -0,8 | n.d. | n.d. | n.d. | 0,8 | -0,9 | 0,2 | n.d. | n.d. | n.d. | -0,5 | 0,6 | -0,1 | 2,3 | 1,9 | n.d. |
| **PLK4** | 0,2 | 1 | 2 | 1,5 | 1,9 | 4,4 | 0,2 | 1,3 | 0,5 | 0,4 | 3,6 | 4,5 | 7 | 1,8 | 1 | 7,1 |
| **RSK1_b** | 0,1 | 0,1 | 0,9 | n.d. | -0,1 | 0,1 | 0,1 | n.d. | n.d. | n.d. | 1,8 | 2 | 2,8 | 0,4 | 0,1 | n.d. |
| **STLK3** | -0,6 | 0,5 | 1 | -0,1 | 0,7 | 0 | -0,1 | 0,5 | 0,1 | -0,1 | -0,1 | 1,9 | 1,7 | 0,1 | -0,1 | 2,2 |
| **TTK** | -0,5 | -0,1 | -0,5 | n.d. | 0,2 | 1,6 | 0,5 | n.d. | n.d. | n.d. | 1 | 2,2 | -1,1 | 2,3 | 0,7 | n.d. |
| **ULK3** | 1 | 1,7 | 1,8 | 2,5 | 3,4 | 1,5 | 2,6 | 2,4 | 0,6 | 2,1 | 1,8 | 8,8 | 5,3 | 6,5 | 4,4 | 6,9 |
| **Wnk1** | 0,5 | 0,3 | 0,3 | 0,1 | 0,2 | 0,4 | 0,5 | 0,5 | 0,5 | 0,3 | 0,2 | 0,3 | 0,3 | 0,2 | 0,5 | 0,4 |

Table S2: Data Collection and Refinement Statistics

| | **CK2A1-34** | **CK2A1-IC20** |
|---|---|---|
| Beamline | X06DA/PXIII SLS | X06DA/PXIII SLS |
| Wavelength (Å) | 0.999998 | 1.000031 |
| Space group | $P4_32_12$ | $P4_32_12$ |
| Cell dimensions | | |
| a, b, c (Å) | 128.65, 128.65, 125.45 | 125.49, 125.49, 124.59 |
| α, β, γ (°) | 90, 90, 90 | 90, 90, 90 |
| Resolution (Å)* | 45.48-2.40 (2.49-2.40) | 44.37-2.65 (2.78-2.65) |
| unique observations* | 41786 (4317) | 26481 (3806) |
| Rpim* | 0.027 (0.426) | 0.045 (0.316) |
| Completeness (%)* | 100.0 (100.0) | 99.9 (99.9) |
| Multiplicity* | 14.9 (14.5) | 9.6 (10.1) |
| mean I/σI* | 16.9 (2.0) | 10.5 (1.9) |
| CC1/2* | 0.99 (0.72) | 0.99 (0.92) |
| Refinement | | |
| $R_{work}$ / $R_{free}$ | 0.217 / 0.232 | 0.263 / 0.284 |
| No. of atoms | 5594 | 5421 |
| overall B-factors ($Å^2$) | 56.53 | 68.33 |
| Rms deviations | | |
| Bond lengths (Å) | 0.0098 | 0.0094 |
| Bond angles (°) | 1.459 | 1.376 |
| Ramachandran (%) | | |
| favored | 96.5 | 96.8 |
| allowed | 2.9 | 2.9 |
| outlier | 0.6 | 0.3 |
| PDB entry | 6YUL | 6YUM |

*Values for the highest resolution shell are shown in parentheses.

**Table S3**: NanoBRET target engagement assay conditions.

Tracer concentration was determined as the $K_{D,app}$ by a tracer titration experiment.

| **Target** | **N-Luc location** | **Promega Catalog No** | **Tracer** | **Promega Catalog No** | **[Tracer], [M]** |
|---|---|---|---|---|---|
| CSNK2A2 | C | NV1191 | K10 | N2640 | 8.29E-07 |
| CSNK2A1 | C | NV2981 | K10 | N2640 | 2.56E-07 |

**Table S4**: NanoBRET screening of CK2 inhibitors.

S/N = signal/noise ratio, z' calculated as z'=((1-(3*(Err Tracer Control/Err Background))/(Tracer Control Signal/Background signal)). Lysed NanoBRET assay quality was low for CK2 and $IC_{50}$s reported should serve as a range.

| | **CK2a1** | **CK2a2** | **CK2a1 lysed** | **CK2a2 lysed** |
|---|---|---|---|---|
| S/N | 1.72 | 4.36 | 1.62 | 2.29 |
| z' | 0.52 | 0.79 | -0.35 | -0.44 |
| IC19 (**32**) | 1.54E-06 | 7.39E-06 | 4.00E-08 | 7.71E-07 |
| **22** | > 50 µM | > 50 µM | 7.52E-06 | > 50 µM |
| **23** | > 50 µM | > 50 µM | 7.09E-05 | > 50 µM |
| IC20 (**31**) | 1.51E-06 | 7.64E-06 | 8.30E-09 | 3.86E-08 |
| **33** | > 50 µM | > 50 µM | 1.32E-05 | > 50 µM |
| **34** | 5.73E-06 | 1.62E-05 | 7.47E-08 | 3.03E-07 |
| **silmitasertib** | 9.66E-08 | 5.52E-07 | 6.04E-09 | 1.63E-07 |

**Table S5**: Kinome scan of IC19 (**32**) @ 1 µM against 469 kinases from DiscoverX/Eurofins.

| Compound Name | DiscoveRx Gene Symbol | Entrez Gene Symbol | Percent Control |
|---|---|---|---|
| IC19 | AAK1 | AAK1 | 63 |
| IC19 | ABL1-nonphosphorylated | ABL1 | 67 |
| IC19 | ABL1-phosphorylated | ABL1 | 100 |
| IC19 | ABL1(E255K)-phosphorylated | ABL1 | 85 |
| IC19 | ABL1(F317I)-nonphosphorylated | ABL1 | 70 |
| IC19 | ABL1(F317I)-phosphorylated | ABL1 | 97 |
| IC19 | ABL1(F317L)-nonphosphorylated | ABL1 | 94 |
| IC19 | ABL1(F317L)-phosphorylated | ABL1 | 85 |
| IC19 | ABL1(H396P)-nonphosphorylated | ABL1 | 59 |
| IC19 | ABL1(H396P)-phosphorylated | ABL1 | 92 |
| IC19 | ABL1(M351T)-phosphorylated | ABL1 | 90 |
| IC19 | ABL1(Q252H)-nonphosphorylated | ABL1 | 73 |
| IC19 | ABL1(Q252H)-phosphorylated | ABL1 | 100 |
| IC19 | ABL1(T315I)-nonphosphorylated | ABL1 | 86 |
| IC19 | ABL1(T315I)-phosphorylated | ABL1 | 92 |
| IC19 | ABL1(Y253F)-phosphorylated | ABL1 | 100 |
| IC19 | ABL2 | ABL2 | 100 |
| IC19 | ACVR1 | ACVR1 | 78 |
| IC19 | ACVR1B | ACVR1B | 78 |
| IC19 | ACVR2A | ACVR2A | 100 |
| IC19 | ACVR2B | ACVR2B | 38 |
| IC19 | ACVRL1 | ACVRL1 | 63 |
| IC19 | ADCK3 | CABC1 | 100 |
| IC19 | ADCK4 | ADCK4 | 63 |
| IC19 | AKT1 | AKT1 | 100 |
| IC19 | AKT2 | AKT2 | 100 |
| IC19 | AKT3 | AKT3 | 68 |
| IC19 | ALK | ALK | 95 |
| IC19 | ALK(C1156Y) | ALK | 90 |
| IC19 | ALK(L1196M) | ALK | 100 |
| IC19 | AMPK-alpha1 | PRKAA1 | 96 |
| IC19 | AMPK-alpha2 | PRKAA2 | 100 |
| IC19 | ANKK1 | ANKK1 | 55 |
| IC19 | ARK5 | NUAK1 | 73 |
| IC19 | ASK1 | MAP3K5 | 100 |
| IC19 | ASK2 | MAP3K6 | 100 |
| IC19 | AURKA | AURKA | 77 |
| IC19 | AURKB | AURKB | 86 |
| IC19 | AURKC | AURKC | 85 |
| IC19 | AXL | AXL | 99 |
| IC19 | BIKE | BMP2K | 59 |

| | | | |
|---|---|---|---|
| IC19 | BLK | BLK | 64 |
| IC19 | BMPR1A | BMPR1A | 100 |
| IC19 | BMPR1B | BMPR1B | 31 |
| IC19 | BMPR2 | BMPR2 | 10 |
| IC19 | BMX | BMX | 100 |
| IC19 | BRAF | BRAF | 96 |
| IC19 | BRAF(V600E) | BRAF | 71 |
| IC19 | BRK | PTK6 | 96 |
| IC19 | BRSK1 | BRSK1 | 76 |
| IC19 | BRSK2 | BRSK2 | 100 |
| IC19 | BTK | BTK | 94 |
| IC19 | BUB1 | BUB1 | 3,9 |
| IC19 | CAMK1 | CAMK1 | 100 |
| IC19 | CAMK1B | PNCK | 59 |
| IC19 | CAMK1D | CAMK1D | 94 |
| IC19 | CAMK1G | CAMK1G | 100 |
| IC19 | CAMK2A | CAMK2A | 17 |
| IC19 | CAMK2B | CAMK2B | 36 |
| IC19 | CAMK2D | CAMK2D | 46 |
| IC19 | CAMK2G | CAMK2G | 67 |
| IC19 | CAMK4 | CAMK4 | 100 |
| IC19 | CAMKK1 | CAMKK1 | 46 |
| IC19 | CAMKK2 | CAMKK2 | 34 |
| IC19 | CASK | CASK | 73 |
| IC19 | CDC2L1 | CDK11B | 73 |
| IC19 | CDC2L2 | CDC2L2 | 100 |
| IC19 | CDC2L5 | CDK13 | 100 |
| IC19 | CDK11 | CDK19 | 42 |
| IC19 | CDK2 | CDK2 | 100 |
| IC19 | CDK3 | CDK3 | 89 |
| IC19 | CDK4 | CDK4 | 74 |
| IC19 | CDK4-cyclinD1 | CDK4 | 79 |
| IC19 | CDK4-cyclinD3 | CDK4 | 99 |
| IC19 | CDK5 | CDK5 | 87 |
| IC19 | CDK7 | CDK7 | 51 |
| IC19 | CDK8 | CDK8 | 23 |
| IC19 | CDK9 | CDK9 | 88 |
| IC19 | CDKL1 | CDKL1 | 77 |
| IC19 | CDKL2 | CDKL2 | 40 |
| IC19 | CDKL3 | CDKL3 | 100 |
| IC19 | CDKL5 | CDKL5 | 62 |
| IC19 | CHEK1 | CHEK1 | 100 |
| IC19 | CHEK2 | CHEK2 | 100 |
| IC19 | CIT | CIT | 70 |

| | | | |
|---|---|---|---|
| IC19 | CLK1 | CLK1 | 68 |
| IC19 | CLK2 | CLK2 | 25 |
| IC19 | CLK3 | CLK3 | 74 |
| IC19 | CLK4 | CLK4 | 100 |
| IC19 | CSF1R | CSF1R | 94 |
| IC19 | CSF1R-autoinhibited | CSF1R | 97 |
| IC19 | CSK | CSK | 100 |
| IC19 | CSNK1A1 | CSNK1A1 | 89 |
| IC19 | CSNK1A1L | CSNK1A1L | 96 |
| IC19 | CSNK1D | CSNK1D | 100 |
| IC19 | CSNK1E | CSNK1E | 90 |
| IC19 | CSNK1G1 | CSNK1G1 | 43 |
| IC19 | CSNK1G2 | CSNK1G2 | 67 |
| IC19 | CSNK1G3 | CSNK1G3 | 81 |
| IC19 | CSNK2A1 | CSNK2A1 | 12 |
| IC19 | CSNK2A2 | CSNK2A2 | 0 |
| IC19 | CTK | MATK | 69 |
| IC19 | DAPK1 | DAPK1 | 52 |
| IC19 | DAPK2 | DAPK2 | 23 |
| IC19 | DAPK3 | DAPK3 | 20 |
| IC19 | DCAMKL1 | DCLK1 | 85 |
| IC19 | DCAMKL2 | DCLK2 | 97 |
| IC19 | DCAMKL3 | DCLK3 | 73 |
| IC19 | DDR1 | DDR1 | 97 |
| IC19 | DDR2 | DDR2 | 99 |
| IC19 | DLK | MAP3K12 | 8 |
| IC19 | DMPK | DMPK | 88 |
| IC19 | DMPK2 | CDC42BPG | 95 |
| IC19 | DRAK1 | STK17A | 72 |
| IC19 | DRAK2 | STK17B | 8,7 |
| IC19 | DYRK1A | DYRK1A | 69 |
| IC19 | DYRK1B | DYRK1B | 76 |
| IC19 | DYRK2 | DYRK2 | 54 |
| IC19 | EGFR | EGFR | 100 |
| IC19 | EGFR(E746-A750del) | EGFR | 89 |
| IC19 | EGFR(G719C) | EGFR | 89 |
| IC19 | EGFR(G719S) | EGFR | 100 |
| IC19 | EGFR(L747-E749del, A750P) | EGFR | 95 |
| IC19 | EGFR(L747-S752del, P753S) | EGFR | 74 |
| IC19 | EGFR(L747-T751del,Sins) | EGFR | 100 |
| IC19 | EGFR(L858R,T790M) | EGFR | 100 |
| IC19 | EGFR(L858R) | EGFR | 100 |
| IC19 | EGFR(L861Q) | EGFR | 99 |
| IC19 | EGFR(S752-I759del) | EGFR | 100 |

| | | | |
|---|---|---|---|
| IC19 | EGFR(T790M) | EGFR | 0 |
| IC19 | EIF2AK1 | EIF2AK1 | 84 |
| IC19 | EPHA1 | EPHA1 | 100 |
| IC19 | EPHA2 | EPHA2 | 91 |
| IC19 | EPHA3 | EPHA3 | 73 |
| IC19 | EPHA4 | EPHA4 | 100 |
| IC19 | EPHA5 | EPHA5 | 99 |
| IC19 | EPHA6 | EPHA6 | 100 |
| IC19 | EPHA7 | EPHA7 | 100 |
| IC19 | EPHA8 | EPHA8 | 100 |
| IC19 | EPHB1 | EPHB1 | 100 |
| IC19 | EPHB2 | EPHB2 | 100 |
| IC19 | EPHB3 | EPHB3 | 100 |
| IC19 | EPHB4 | EPHB4 | 100 |
| IC19 | EPHB6 | EPHB6 | 68 |
| IC19 | ERBB2 | ERBB2 | 100 |
| IC19 | ERBB3 | ERBB3 | 100 |
| IC19 | ERBB4 | ERBB4 | 92 |
| IC19 | ERK1 | MAPK3 | 66 |
| IC19 | ERK2 | MAPK1 | 86 |
| IC19 | ERK3 | MAPK6 | 94 |
| IC19 | ERK4 | MAPK4 | 90 |
| IC19 | ERK5 | MAPK7 | 99 |
| IC19 | ERK8 | MAPK15 | 6,6 |
| IC19 | ERN1 | ERN1 | 71 |
| IC19 | FAK | PTK2 | 100 |
| IC19 | FER | FER | 100 |
| IC19 | FES | FES | 100 |
| IC19 | FGFR1 | FGFR1 | 100 |
| IC19 | FGFR2 | FGFR2 | 94 |
| IC19 | FGFR3 | FGFR3 | 100 |
| IC19 | FGFR3(G697C) | FGFR3 | 100 |
| IC19 | FGFR4 | FGFR4 | 91 |
| IC19 | FGR | FGR | 100 |
| IC19 | FLT1 | FLT1 | 100 |
| IC19 | FLT3 | FLT3 | 91 |
| IC19 | FLT3-autoinhibited | FLT3 | 76 |
| IC19 | FLT3(D835H) | FLT3 | 51 |
| IC19 | FLT3(D835V) | FLT3 | 9,5 |
| IC19 | FLT3(D835Y) | FLT3 | 22 |
| IC19 | FLT3(ITD,D835V) | FLT3 | 24 |
| IC19 | FLT3(ITD,F691L) | FLT3 | 55 |
| IC19 | FLT3(ITD) | FLT3 | 79 |
| IC19 | FLT3(K663Q) | FLT3 | 61 |

| | | | |
|---|---|---|---|
| IC19 | FLT3(N841I) | FLT3 | 57 |
| IC19 | FLT3(R834Q) | FLT3 | 88 |
| IC19 | FLT4 | FLT4 | 100 |
| IC19 | FRK | FRK | 100 |
| IC19 | FYN | FYN | 65 |
| IC19 | GAK | GAK | 8,1 |
| IC19 | GCN2(Kin.Dom.2,S808G) | EIF2AK4 | 98 |
| IC19 | GRK1 | GRK1 | 88 |
| IC19 | GRK2 | ADRBK1 | 49 |
| IC19 | GRK3 | ADRBK2 | 100 |
| IC19 | GRK4 | GRK4 | 52 |
| IC19 | GRK7 | GRK7 | 100 |
| IC19 | GSK3A | GSK3A | 51 |
| IC19 | GSK3B | GSK3B | 95 |
| IC19 | HASPIN | GSG2 | 100 |
| IC19 | HCK | HCK | 100 |
| IC19 | HIPK1 | HIPK1 | 18 |
| IC19 | HIPK2 | HIPK2 | 25 |
| IC19 | HIPK3 | HIPK3 | 56 |
| IC19 | HIPK4 | HIPK4 | 0,1 |
| IC19 | HPK1 | MAP4K1 | 100 |
| IC19 | HUNK | HUNK | 100 |
| IC19 | ICK | ICK | 2,2 |
| IC19 | IGF1R | IGF1R | 78 |
| IC19 | IKK-alpha | CHUK | 92 |
| IC19 | IKK-beta | IKBKB | 100 |
| IC19 | IKK-epsilon | IKBKE | 100 |
| IC19 | INSR | INSR | 0 |
| IC19 | INSRR | INSRR | 78 |
| IC19 | IRAK1 | IRAK1 | 69 |
| IC19 | IRAK3 | IRAK3 | 74 |
| IC19 | IRAK4 | IRAK4 | 86 |
| IC19 | ITK | ITK | 100 |
| IC19 | JAK1(JH1domain-catalytic) | JAK1 | 69 |
| IC19 | JAK1(JH2domain-pseudokinase) | JAK1 | 100 |
| IC19 | JAK2(JH1domain-catalytic) | JAK2 | 86 |
| IC19 | JAK3(JH1domain-catalytic) | JAK3 | 73 |
| IC19 | JNK1 | MAPK8 | 68 |
| IC19 | JNK2 | MAPK9 | 66 |
| IC19 | JNK3 | MAPK10 | 61 |
| IC19 | KIT | KIT | 96 |
| IC19 | KIT-autoinhibited | KIT | 100 |
| IC19 | KIT(A829P) | KIT | 86 |
| IC19 | KIT(D816H) | KIT | 68 |

| | | | |
|---|---|---|---|
| IC19 | KIT(D816V) | KIT | 70 |
| IC19 | KIT(L576P) | KIT | 91 |
| IC19 | KIT(V559D,T670I) | KIT | 94 |
| IC19 | KIT(V559D,V654A) | KIT | 100 |
| IC19 | KIT(V559D) | KIT | 90 |
| IC19 | LATS1 | LATS1 | 100 |
| IC19 | LATS2 | LATS2 | 100 |
| IC19 | LCK | LCK | 100 |
| IC19 | LIMK1 | LIMK1 | 92 |
| IC19 | LIMK2 | LIMK2 | 77 |
| IC19 | LKB1 | STK11 | 100 |
| IC19 | LOK | STK10 | 88 |
| IC19 | LRRK2 | LRRK2 | 74 |
| IC19 | LRRK2(G2019S) | LRRK2 | 100 |
| IC19 | LTK | LTK | 100 |
| IC19 | LYN | LYN | 94 |
| IC19 | LZK | MAP3K13 | 92 |
| IC19 | MAK | MAK | 52 |
| IC19 | MAP3K1 | MAP3K1 | 92 |
| IC19 | MAP3K15 | MAP3K15 | 71 |
| IC19 | MAP3K2 | MAP3K2 | 96 |
| IC19 | MAP3K3 | MAP3K3 | 82 |
| IC19 | MAP3K4 | MAP3K4 | 92 |
| IC19 | MAP4K2 | MAP4K2 | 100 |
| IC19 | MAP4K3 | MAP4K3 | 97 |
| IC19 | MAP4K4 | MAP4K4 | 78 |
| IC19 | MAP4K5 | MAP4K5 | 72 |
| IC19 | MAPKAPK2 | MAPKAPK2 | 100 |
| IC19 | MAPKAPK5 | MAPKAPK5 | 94 |
| IC19 | MARK1 | MARK1 | 100 |
| IC19 | MARK2 | MARK2 | 60 |
| IC19 | MARK3 | MARK3 | 97 |
| IC19 | MARK4 | MARK4 | 100 |
| IC19 | MAST1 | MAST1 | 98 |
| IC19 | MEK1 | MAP2K1 | 86 |
| IC19 | MEK2 | MAP2K2 | 97 |
| IC19 | MEK3 | MAP2K3 | 78 |
| IC19 | MEK4 | MAP2K4 | 43 |
| IC19 | MEK5 | MAP2K5 | 25 |
| IC19 | MEK6 | MAP2K6 | 100 |
| IC19 | MELK | MELK | 100 |
| IC19 | MERTK | MERTK | 90 |
| IC19 | MET | MET | 74 |
| IC19 | MET(M1250T) | MET | 100 |

| | | | |
|---|---|---|---|
| IC19 | MET(Y1235D) | MET | 100 |
| IC19 | MINK | MINK1 | 78 |
| IC19 | MKK7 | MAP2K7 | 93 |
| IC19 | MKNK1 | MKNK1 | 96 |
| IC19 | MKNK2 | MKNK2 | 58 |
| IC19 | MLCK | MYLK3 | 95 |
| IC19 | MLK1 | MAP3K9 | 65 |
| IC19 | MLK2 | MAP3K10 | 100 |
| IC19 | MLK3 | MAP3K11 | 84 |
| IC19 | MRCKA | CDC42BPA | 100 |
| IC19 | MRCKB | CDC42BPB | 100 |
| IC19 | MST1 | STK4 | 100 |
| IC19 | MST1R | MST1R | 100 |
| IC19 | MST2 | STK3 | 96 |
| IC19 | MST3 | STK24 | 87 |
| IC19 | MST4 | MST4 | 99 |
| IC19 | MTOR | MTOR | 100 |
| IC19 | MUSK | MUSK | 85 |
| IC19 | MYLK | MYLK | 81 |
| IC19 | MYLK2 | MYLK2 | 88 |
| IC19 | MYLK4 | MYLK4 | 59 |
| IC19 | MYO3A | MYO3A | 95 |
| IC19 | MYO3B | MYO3B | 100 |
| IC19 | NDR1 | STK38 | 87 |
| IC19 | NDR2 | STK38L | 100 |
| IC19 | NEK1 | NEK1 | 100 |
| IC19 | NEK10 | NEK10 | 47 |
| IC19 | NEK11 | NEK11 | 100 |
| IC19 | NEK2 | NEK2 | 61 |
| IC19 | NEK3 | NEK3 | 84 |
| IC19 | NEK4 | NEK4 | 100 |
| IC19 | NEK5 | NEK5 | 95 |
| IC19 | NEK6 | NEK6 | 78 |
| IC19 | NEK7 | NEK7 | 100 |
| IC19 | NEK9 | NEK9 | 96 |
| IC19 | NIK | MAP3K14 | 97 |
| IC19 | NIM1 | MGC42105 | 99 |
| IC19 | NLK | NLK | 66 |
| IC19 | OSR1 | OXSR1 | 96 |
| IC19 | p38-alpha | MAPK14 | 100 |
| IC19 | p38-beta | MAPK11 | 100 |
| IC19 | p38-delta | MAPK13 | 100 |
| IC19 | p38-gamma | MAPK12 | 100 |
| IC19 | PAK1 | PAK1 | 0 |

| | | | |
|---|---|---|---|
| IC19 | PAK2 | PAK2 | 0 |
| IC19 | PAK3 | PAK3 | 100 |
| IC19 | PAK4 | PAK4 | 100 |
| IC19 | PAK6 | PAK6 | 100 |
| IC19 | PAK7 | PAK7 | 78 |
| IC19 | PCTK1 | CDK16 | 67 |
| IC19 | PCTK2 | CDK17 | 81 |
| IC19 | PCTK3 | CDK18 | 86 |
| IC19 | PDGFRA | PDGFRA | 72 |
| IC19 | PDGFRB | PDGFRB | 88 |
| IC19 | PDPK1 | PDPK1 | 100 |
| IC19 | PFCDPK1(P.falciparum) | CDPK1 | 100 |
| IC19 | PFPK5(P.falciparum) | MAL13P1.279 | 99 |
| IC19 | PFTAIRE2 | CDK15 | 87 |
| IC19 | PFTK1 | CDK14 | 93 |
| IC19 | PHKG1 | PHKG1 | 91 |
| IC19 | PHKG2 | PHKG2 | 92 |
| IC19 | PIK3C2B | PIK3C2B | 64 |
| IC19 | PIK3C2G | PIK3C2G | 32 |
| IC19 | PIK3CA | PIK3CA | 32 |
| IC19 | PIK3CA(C420R) | PIK3CA | 54 |
| IC19 | PIK3CA(E542K) | PIK3CA | 26 |
| IC19 | PIK3CA(E545A) | PIK3CA | 54 |
| IC19 | PIK3CA(E545K) | PIK3CA | 64 |
| IC19 | PIK3CA(H1047L) | PIK3CA | 36 |
| IC19 | PIK3CA(H1047Y) | PIK3CA | 42 |
| IC19 | PIK3CA(I800L) | PIK3CA | 0 |
| IC19 | PIK3CA(M1043I) | PIK3CA | 39 |
| IC19 | PIK3CA(Q546K) | PIK3CA | 18 |
| IC19 | PIK3CB | PIK3CB | 76 |
| IC19 | PIK3CD | PIK3CD | 58 |
| IC19 | PIK3CG | PIK3CG | 27 |
| IC19 | PIK4CB | PI4KB | 90 |
| IC19 | PIKFYVE | PIKFYVE | 99 |
| IC19 | PIM1 | PIM1 | 93 |
| IC19 | PIM2 | PIM2 | 73 |
| IC19 | PIM3 | PIM3 | 0 |
| IC19 | PIP5K1A | PIP5K1A | 86 |
| IC19 | PIP5K1C | PIP5K1C | 77 |
| IC19 | PIP5K2B | PIP4K2B | 100 |
| IC19 | PIP5K2C | PIP4K2C | 4,6 |
| IC19 | PKAC-alpha | PRKACA | 100 |
| IC19 | PKAC-beta | PRKACB | 99 |
| IC19 | PKMYT1 | PKMYT1 | 100 |

| | | | |
|---|---|---|---|
| IC19 | PKN1 | PKN1 | 97 |
| IC19 | PKN2 | PKN2 | 92 |
| IC19 | PKNB(M.tuberculosis) | pknB | 0 |
| IC19 | PLK1 | PLK1 | 100 |
| IC19 | PLK2 | PLK2 | 89 |
| IC19 | PLK3 | PLK3 | 87 |
| IC19 | PLK4 | PLK4 | 80 |
| IC19 | PRKCD | PRKCD | 100 |
| IC19 | PRKCE | PRKCE | 76 |
| IC19 | PRKCH | PRKCH | 100 |
| IC19 | PRKCI | PRKCI | 73 |
| IC19 | PRKCQ | PRKCQ | 100 |
| IC19 | PRKD1 | PRKD1 | 100 |
| IC19 | PRKD2 | PRKD2 | 88 |
| IC19 | PRKD3 | PRKD3 | 84 |
| IC19 | PRKG1 | PRKG1 | 86 |
| IC19 | PRKG2 | PRKG2 | 100 |
| IC19 | PRKR | EIF2AK2 | 100 |
| IC19 | PRKX | PRKX | 99 |
| IC19 | PRP4 | PRPF4B | 100 |
| IC19 | PYK2 | PTK2B | 100 |
| IC19 | QSK | KIAA0999 | 100 |
| IC19 | RAF1 | RAF1 | 100 |
| IC19 | RET | RET | 81 |
| IC19 | RET(M918T) | RET | 86 |
| IC19 | RET(V804L) | RET | 93 |
| IC19 | RET(V804M) | RET | 99 |
| IC19 | RIOK1 | RIOK1 | 82 |
| IC19 | RIOK2 | RIOK2 | 94 |
| IC19 | RIOK3 | RIOK3 | 89 |
| IC19 | RIPK1 | RIPK1 | 92 |
| IC19 | RIPK2 | RIPK2 | 38 |
| IC19 | RIPK4 | RIPK4 | 93 |
| IC19 | RIPK5 | DSTYK | 80 |
| IC19 | ROCK1 | ROCK1 | 100 |
| IC19 | ROCK2 | ROCK2 | 99 |
| IC19 | ROS1 | ROS1 | 100 |
| IC19 | RPS6KA4(Kin.Dom.1-N-terminal) | RPS6KA4 | 94 |
| IC19 | RPS6KA4(Kin.Dom.2-C-terminal) | RPS6KA4 | 72 |
| IC19 | RPS6KA5(Kin.Dom.1-N-terminal) | RPS6KA5 | 100 |
| IC19 | RPS6KA5(Kin.Dom.2-C-terminal) | RPS6KA5 | 98 |
| IC19 | RSK1(Kin.Dom.1-N-terminal) | RPS6KA1 | 77 |
| IC19 | RSK1(Kin.Dom.2-C-terminal) | RPS6KA1 | 38 |
| IC19 | RSK2(Kin.Dom.1-N-terminal) | RPS6KA3 | 47 |

| | | | |
|---|---|---|---|
| IC19 | RSK2(Kin.Dom.2-C-terminal) | RPS6KA3 | 26 |
| IC19 | RSK3(Kin.Dom.1-N-terminal) | RPS6KA2 | 100 |
| IC19 | RSK3(Kin.Dom.2-C-terminal) | RPS6KA2 | 50 |
| IC19 | RSK4(Kin.Dom.1-N-terminal) | RPS6KA6 | 29 |
| IC19 | RSK4(Kin.Dom.2-C-terminal) | RPS6KA6 | 54 |
| IC19 | S6K1 | RPS6KB1 | 94 |
| IC19 | SBK1 | SBK1 | 73 |
| IC19 | SGK | SGK1 | 23 |
| IC19 | SgK110 | SgK110 | 100 |
| IC19 | SGK2 | SGK2 | 53 |
| IC19 | SGK3 | SGK3 | 51 |
| IC19 | SIK | SIK1 | 87 |
| IC19 | SIK2 | SIK2 | 73 |
| IC19 | SLK | SLK | 100 |
| IC19 | SNARK | NUAK2 | 70 |
| IC19 | SNRK | SNRK | 100 |
| IC19 | SRC | SRC | 100 |
| IC19 | SRMS | SRMS | 100 |
| IC19 | SRPK1 | SRPK1 | 91 |
| IC19 | SRPK2 | SRPK2 | 100 |
| IC19 | SRPK3 | SRPK3 | 61 |
| IC19 | STK16 | STK16 | 50 |
| IC19 | STK33 | STK33 | 96 |
| IC19 | STK35 | STK35 | 87 |
| IC19 | STK36 | STK36 | 100 |
| IC19 | STK39 | STK39 | 83 |
| IC19 | SYK | SYK | 100 |
| IC19 | TAK1 | MAP3K7 | 85 |
| IC19 | TAOK1 | TAOK1 | 87 |
| IC19 | TAOK2 | TAOK2 | 100 |
| IC19 | TAOK3 | TAOK3 | 42 |
| IC19 | TBK1 | TBK1 | 70 |
| IC19 | TEC | TEC | 100 |
| IC19 | TESK1 | TESK1 | 100 |
| IC19 | TGFBR1 | TGFBR1 | 100 |
| IC19 | TGFBR2 | TGFBR2 | 53 |
| IC19 | TIE1 | TIE1 | 74 |
| IC19 | TIE2 | TEK | 100 |
| IC19 | TLK1 | TLK1 | 86 |
| IC19 | TLK2 | TLK2 | 100 |
| IC19 | TNIK | TNIK | 80 |
| IC19 | TNK1 | TNK1 | 100 |
| IC19 | TNK2 | TNK2 | 96 |
| IC19 | TNNI3K | TNNI3K | 75 |

| | | | |
|---|---|---|---|
| IC19 | TRKA | NTRK1 | 79 |
| IC19 | TRKB | NTRK2 | 73 |
| IC19 | TRKC | NTRK3 | 91 |
| IC19 | TRPM6 | TRPM6 | 100 |
| IC19 | TSSK1B | TSSK1B | 100 |
| IC19 | TSSK3 | TSSK3 | 88 |
| IC19 | TTK | TTK | 73 |
| IC19 | TXK | TXK | 100 |
| IC19 | TYK2(JH1domain-catalytic) | TYK2 | 95 |
| IC19 | TYK2(JH2domain-pseudokinase) | TYK2 | 81 |
| IC19 | TYRO3 | TYRO3 | 100 |
| IC19 | ULK1 | ULK1 | 78 |
| IC19 | ULK2 | ULK2 | 64 |
| IC19 | ULK3 | ULK3 | 86 |
| IC19 | VEGFR2 | KDR | 77 |
| IC19 | VPS34 | PIK3C3 | 28 |
| IC19 | VRK2 | VRK2 | 99 |
| IC19 | WEE1 | WEE1 | 87 |
| IC19 | WEE2 | WEE2 | 92 |
| IC19 | WNK1 | WNK1 | 93 |
| IC19 | WNK2 | WNK2 | 86 |
| IC19 | WNK3 | WNK3 | 89 |
| IC19 | WNK4 | WNK4 | 92 |
| IC19 | YANK1 | STK32A | 84 |
| IC19 | YANK2 | STK32B | 100 |
| IC19 | YANK3 | STK32C | 92 |
| IC19 | YES | YES1 | 92 |
| IC19 | YSK1 | STK25 | 100 |
| IC19 | YSK4 | MAP3K19 | 52 |
| IC19 | ZAK | ZAK | 100 |
| IC19 | ZAP70 | ZAP70 | 100 |

Table S6: Kinome scan of IC20 (31) @ 1 µM against 469 kinases from DiscoverX/Eurofins.

| Compound Name | DiscoveRx Gene Symbol | Entrez Gene Symbol | Percent Control |
|---|---|---|---|
| IC20 | AAK1 | AAK1 | 90 |
| IC20 | ABL1(E255K)-phosphorylated | ABL1 | 82 |
| IC20 | ABL1(F317I)-nonphosphorylated | ABL1 | 91 |
| IC20 | ABL1(F317I)-phosphorylated | ABL1 | 90 |
| IC20 | ABL1(F317L)-nonphosphorylated | ABL1 | 97 |
| IC20 | ABL1(F317L)-phosphorylated | ABL1 | 100 |
| IC20 | ABL1(H396P)-nonphosphorylated | ABL1 | 81 |
| IC20 | ABL1(H396P)-phosphorylated | ABL1 | 94 |
| IC20 | ABL1(M351T)-phosphorylated | ABL1 | 97 |
| IC20 | ABL1(Q252H)-nonphosphorylated | ABL1 | 72 |
| IC20 | ABL1(Q252H)-phosphorylated | ABL1 | 100 |
| IC20 | ABL1(T315I)-nonphosphorylated | ABL1 | 100 |
| IC20 | ABL1(T315I)-phosphorylated | ABL1 | 91 |
| IC20 | ABL1(Y253F)-phosphorylated | ABL1 | 86 |
| IC20 | ABL1-nonphosphorylated | ABL1 | 85 |
| IC20 | ABL1-phosphorylated | ABL1 | 92 |
| IC20 | ABL2 | ABL2 | 100 |
| IC20 | ACVR1 | ACVR1 | 100 |
| IC20 | ACVR1B | ACVR1B | 78 |
| IC20 | ACVR2A | ACVR2A | 65 |
| IC20 | ACVR2B | ACVR2B | 74 |
| IC20 | ACVRL1 | ACVRL1 | 71 |
| IC20 | ADCK3 | CABC1 | 61 |
| IC20 | ADCK4 | ADCK4 | 100 |
| IC20 | AKT1 | AKT1 | 84 |
| IC20 | AKT2 | AKT2 | 100 |
| IC20 | AKT3 | AKT3 | 90 |
| IC20 | ALK | ALK | 100 |
| IC20 | ALK(C1156Y) | ALK | 100 |
| IC20 | ALK(L1196M) | ALK | 100 |
| IC20 | AMPK-alpha1 | PRKAA1 | 100 |
| IC20 | AMPK-alpha2 | PRKAA2 | 89 |
| IC20 | ANKK1 | ANKK1 | 100 |
| IC20 | ARK5 | NUAK1 | 97 |
| IC20 | ASK1 | MAP3K5 | 100 |
| IC20 | ASK2 | MAP3K6 | 97 |
| IC20 | AURKA | AURKA | 98 |
| IC20 | AURKB | AURKB | 89 |
| IC20 | AURKC | AURKC | 90 |
| IC20 | AXL | AXL | 100 |
| IC20 | BIKE | BMP2K | 61 |
| IC20 | BLK | BLK | 100 |

| | | | |
|---|---|---|---|
| IC20 | BMPR1A | BMPR1A | 87 |
| IC20 | BMPR1B | BMPR1B | 88 |
| IC20 | BMPR2 | BMPR2 | 52 |
| IC20 | BMX | BMX | 41 |
| IC20 | BRAF | BRAF | 100 |
| IC20 | BRAF(V600E) | BRAF | 100 |
| IC20 | BRK | PTK6 | 80 |
| IC20 | BRSK1 | BRSK1 | 93 |
| IC20 | BRSK2 | BRSK2 | 75 |
| IC20 | BTK | BTK | 100 |
| IC20 | BUB1 | BUB1 | 100 |
| IC20 | CAMK1 | CAMK1 | 89 |
| IC20 | CAMK1B | PNCK | 100 |
| IC20 | CAMK1D | CAMK1D | 91 |
| IC20 | CAMK1G | CAMK1G | 100 |
| IC20 | CAMK2A | CAMK2A | 89 |
| IC20 | CAMK2B | CAMK2B | 100 |
| IC20 | CAMK2D | CAMK2D | 98 |
| IC20 | CAMK2G | CAMK2G | 96 |
| IC20 | CAMK4 | CAMK4 | 75 |
| IC20 | CAMKK1 | CAMKK1 | 84 |
| IC20 | CAMKK2 | CAMKK2 | 88 |
| IC20 | CASK | CASK | 96 |
| IC20 | CDC2L1 | CDK11B | 100 |
| IC20 | CDC2L2 | CDC2L2 | 85 |
| IC20 | CDC2L5 | CDK13 | 100 |
| IC20 | CDK11 | CDK19 | 68 |
| IC20 | CDK2 | CDK2 | 94 |
| IC20 | CDK3 | CDK3 | 94 |
| IC20 | CDK4 | CDK4 | 94 |
| IC20 | CDK4-cyclinD1 | CDK4 | 100 |
| IC20 | CDK4-cyclinD3 | CDK4 | 96 |
| IC20 | CDK5 | CDK5 | 88 |
| IC20 | CDK7 | CDK7 | 84 |
| IC20 | CDK8 | CDK8 | 86 |
| IC20 | CDK9 | CDK9 | 100 |
| IC20 | CDKL1 | CDKL1 | 67 |
| IC20 | CDKL2 | CDKL2 | 95 |
| IC20 | CDKL3 | CDKL3 | 72 |
| IC20 | CDKL5 | CDKL5 | 100 |
| IC20 | CHEK1 | CHEK1 | 100 |
| IC20 | CHEK2 | CHEK2 | 100 |
| IC20 | CIT | CIT | 100 |
| IC20 | CLK1 | CLK1 | 71 |
| IC20 | CLK2 | CLK2 | 100 |
| IC20 | CLK3 | CLK3 | 85 |
| IC20 | CLK4 | CLK4 | 84 |

| | | | |
|---|---|---|---|
| IC20 | CSF1R | CSF1R | 91 |
| IC20 | CSF1R-autoinhibited | CSF1R | 100 |
| IC20 | CSK | CSK | 89 |
| IC20 | CSNK1A1 | CSNK1A1 | 91 |
| IC20 | CSNK1A1L | CSNK1A1L | 87 |
| IC20 | CSNK1D | CSNK1D | 89 |
| IC20 | CSNK1E | CSNK1E | 100 |
| IC20 | CSNK1G1 | CSNK1G1 | 98 |
| IC20 | CSNK1G2 | CSNK1G2 | 99 |
| IC20 | CSNK1G3 | CSNK1G3 | 91 |
| IC20 | CSNK2A1 | CSNK2A1 | 5,3 |
| IC20 | CSNK2A2 | CSNK2A2 | 0 |
| IC20 | CTK | MATK | 100 |
| IC20 | DAPK1 | DAPK1 | 51 |
| IC20 | DAPK2 | DAPK2 | 9,9 |
| IC20 | DAPK3 | DAPK3 | 15 |
| IC20 | DCAMKL1 | DCLK1 | 99 |
| IC20 | DCAMKL2 | DCLK2 | 99 |
| IC20 | DCAMKL3 | DCLK3 | 65 |
| IC20 | DDR1 | DDR1 | 100 |
| IC20 | DDR2 | DDR2 | 100 |
| IC20 | DLK | MAP3K12 | 100 |
| IC20 | DMPK | DMPK | 89 |
| IC20 | DMPK2 | CDC42BPG | 100 |
| IC20 | DRAK1 | STK17A | 81 |
| IC20 | DRAK2 | STK17B | 37 |
| IC20 | DYRK1A | DYRK1A | 30 |
| IC20 | DYRK1B | DYRK1B | 100 |
| IC20 | DYRK2 | DYRK2 | 59 |
| IC20 | EGFR | EGFR | 100 |
| IC20 | EGFR(E746-A750del) | EGFR | 100 |
| IC20 | EGFR(G719C) | EGFR | 79 |
| IC20 | EGFR(G719S) | EGFR | 100 |
| IC20 | EGFR(L747-E749del, A750P) | EGFR | 96 |
| IC20 | EGFR(L747-S752del, P753S) | EGFR | 83 |
| IC20 | EGFR(L747-T751del,Sins) | EGFR | 100 |
| IC20 | EGFR(L858R) | EGFR | 100 |
| IC20 | EGFR(L858R,T790M) | EGFR | 71 |
| IC20 | EGFR(L861Q) | EGFR | 89 |
| IC20 | EGFR(S752-I759del) | EGFR | 96 |
| IC20 | EGFR(T790M) | EGFR | 75 |
| IC20 | EIF2AK1 | EIF2AK1 | 100 |
| IC20 | EPHA1 | EPHA1 | 100 |
| IC20 | EPHA2 | EPHA2 | 44 |
| IC20 | EPHA3 | EPHA3 | 95 |
| IC20 | EPHA4 | EPHA4 | 90 |
| IC20 | EPHA5 | EPHA5 | 99 |

| | | | |
|---|---|---|---|
| IC20 | EPHA6 | EPHA6 | 80 |
| IC20 | EPHA7 | EPHA7 | 90 |
| IC20 | EPHA8 | EPHA8 | 100 |
| IC20 | EPHB1 | EPHB1 | 100 |
| IC20 | EPHB2 | EPHB2 | 90 |
| IC20 | EPHB3 | EPHB3 | 78 |
| IC20 | EPHB4 | EPHB4 | 96 |
| IC20 | EPHB6 | EPHB6 | 76 |
| IC20 | ERBB2 | ERBB2 | 100 |
| IC20 | ERBB3 | ERBB3 | 100 |
| IC20 | ERBB4 | ERBB4 | 74 |
| IC20 | ERK1 | MAPK3 | 100 |
| IC20 | ERK2 | MAPK1 | 96 |
| IC20 | ERK3 | MAPK6 | 100 |
| IC20 | ERK4 | MAPK4 | 100 |
| IC20 | ERK5 | MAPK7 | 100 |
| IC20 | ERK8 | MAPK15 | 84 |
| IC20 | ERN1 | ERN1 | 100 |
| IC20 | FAK | PTK2 | 83 |
| IC20 | FER | FER | 88 |
| IC20 | FES | FES | 86 |
| IC20 | FGFR1 | FGFR1 | 89 |
| IC20 | FGFR2 | FGFR2 | 83 |
| IC20 | FGFR3 | FGFR3 | 82 |
| IC20 | FGFR3(G697C) | FGFR3 | 100 |
| IC20 | FGFR4 | FGFR4 | 95 |
| IC20 | FGR | FGR | 80 |
| IC20 | FLT1 | FLT1 | 80 |
| IC20 | FLT3 | FLT3 | 96 |
| IC20 | FLT3(D835H) | FLT3 | 100 |
| IC20 | FLT3(D835V) | FLT3 | 100 |
| IC20 | FLT3(D835Y) | FLT3 | 80 |
| IC20 | FLT3(ITD) | FLT3 | 82 |
| IC20 | FLT3(ITD,D835V) | FLT3 | 77 |
| IC20 | FLT3(ITD,F691L) | FLT3 | 89 |
| IC20 | FLT3(K663Q) | FLT3 | 76 |
| IC20 | FLT3(N841I) | FLT3 | 87 |
| IC20 | FLT3(R834Q) | FLT3 | 95 |
| IC20 | FLT3-autoinhibited | FLT3 | 85 |
| IC20 | FLT4 | FLT4 | 100 |
| IC20 | FRK | FRK | 72 |
| IC20 | FYN | FYN | 55 |
| IC20 | GAK | GAK | 83 |
| IC20 | GCN2(Kin.Dom.2,S808G) | EIF2AK4 | 100 |
| IC20 | GRK1 | GRK1 | 100 |
| IC20 | GRK2 | ADRBK1 | 100 |
| IC20 | GRK3 | ADRBK2 | 100 |

| | | | |
|---|---|---|---|
| IC20 | GRK4 | GRK4 | 100 |
| IC20 | GRK7 | GRK7 | 100 |
| IC20 | GSK3A | GSK3A | 81 |
| IC20 | GSK3B | GSK3B | 100 |
| IC20 | HASPIN | GSG2 | 100 |
| IC20 | HCK | HCK | 100 |
| IC20 | HIPK1 | HIPK1 | 19 |
| IC20 | HIPK2 | HIPK2 | 43 |
| IC20 | HIPK3 | HIPK3 | 50 |
| IC20 | HIPK4 | HIPK4 | 54 |
| IC20 | HPK1 | MAP4K1 | 98 |
| IC20 | HUNK | HUNK | 99 |
| IC20 | ICK | ICK | 100 |
| IC20 | IGF1R | IGF1R | 93 |
| IC20 | IKK-alpha | CHUK | 100 |
| IC20 | IKK-beta | IKBKB | 80 |
| IC20 | IKK-epsilon | IKBKE | 75 |
| IC20 | INSR | INSR | 100 |
| IC20 | INSRR | INSRR | 100 |
| IC20 | IRAK1 | IRAK1 | 100 |
| IC20 | IRAK3 | IRAK3 | 99 |
| IC20 | IRAK4 | IRAK4 | 100 |
| IC20 | ITK | ITK | 100 |
| IC20 | JAK1(JH1domain-catalytic) | JAK1 | 88 |
| IC20 | JAK1(JH2domain-pseudokinase) | JAK1 | 87 |
| IC20 | JAK2(JH1domain-catalytic) | JAK2 | 100 |
| IC20 | JAK3(JH1domain-catalytic) | JAK3 | 100 |
| IC20 | JNK1 | MAPK8 | 80 |
| IC20 | JNK2 | MAPK9 | 74 |
| IC20 | JNK3 | MAPK10 | 100 |
| IC20 | KIT | KIT | 84 |
| IC20 | KIT(A829P) | KIT | 100 |
| IC20 | KIT(D816H) | KIT | 100 |
| IC20 | KIT(D816V) | KIT | 70 |
| IC20 | KIT(L576P) | KIT | 58 |
| IC20 | KIT(V559D) | KIT | 79 |
| IC20 | KIT(V559D,T670I) | KIT | 80 |
| IC20 | KIT(V559D,V654A) | KIT | 90 |
| IC20 | KIT-autoinhibited | KIT | 100 |
| IC20 | LATS1 | LATS1 | 100 |
| IC20 | LATS2 | LATS2 | 100 |
| IC20 | LCK | LCK | 100 |
| IC20 | LIMK1 | LIMK1 | 100 |
| IC20 | LIMK2 | LIMK2 | 100 |
| IC20 | LKB1 | STK11 | 100 |
| IC20 | LOK | STK10 | 100 |
| IC20 | LRRK2 | LRRK2 | 100 |

| | | | |
|---|---|---|---|
| IC20 | LRRK2(G2019S) | LRRK2 | 72 |
| IC20 | LTK | LTK | 96 |
| IC20 | LYN | LYN | 99 |
| IC20 | LZK | MAP3K13 | 100 |
| IC20 | MAK | MAK | 86 |
| IC20 | MAP3K1 | MAP3K1 | 98 |
| IC20 | MAP3K15 | MAP3K15 | 100 |
| IC20 | MAP3K2 | MAP3K2 | 100 |
| IC20 | MAP3K3 | MAP3K3 | 83 |
| IC20 | MAP3K4 | MAP3K4 | 94 |
| IC20 | MAP4K2 | MAP4K2 | 100 |
| IC20 | MAP4K3 | MAP4K3 | 73 |
| IC20 | MAP4K4 | MAP4K4 | 52 |
| IC20 | MAP4K5 | MAP4K5 | 69 |
| IC20 | MAPKAPK2 | MAPKAPK2 | 84 |
| IC20 | MAPKAPK5 | MAPKAPK5 | 100 |
| IC20 | MARK1 | MARK1 | 73 |
| IC20 | MARK2 | MARK2 | 72 |
| IC20 | MARK3 | MARK3 | 100 |
| IC20 | MARK4 | MARK4 | 76 |
| IC20 | MAST1 | MAST1 | 93 |
| IC20 | MEK1 | MAP2K1 | 100 |
| IC20 | MEK2 | MAP2K2 | 100 |
| IC20 | MEK3 | MAP2K3 | 100 |
| IC20 | MEK4 | MAP2K4 | 100 |
| IC20 | MEK5 | MAP2K5 | 100 |
| IC20 | MEK6 | MAP2K6 | 72 |
| IC20 | MELK | MELK | 100 |
| IC20 | MERTK | MERTK | 100 |
| IC20 | MET | MET | 76 |
| IC20 | MET(M1250T) | MET | 68 |
| IC20 | MET(Y1235D) | MET | 86 |
| IC20 | MINK | MINK1 | 89 |
| IC20 | MKK7 | MAP2K7 | 100 |
| IC20 | MKNK1 | MKNK1 | 97 |
| IC20 | MKNK2 | MKNK2 | 100 |
| IC20 | MLCK | MYLK3 | 82 |
| IC20 | MLK1 | MAP3K9 | 100 |
| IC20 | MLK2 | MAP3K10 | 69 |
| IC20 | MLK3 | MAP3K11 | 83 |
| IC20 | MRCKA | CDC42BPA | 100 |
| IC20 | MRCKB | CDC42BPB | 98 |
| IC20 | MST1 | STK4 | 95 |
| IC20 | MST1R | MST1R | 100 |
| IC20 | MST2 | STK3 | 86 |
| IC20 | MST3 | STK24 | 100 |
| IC20 | MST4 | MST4 | 81 |

| | | | |
|---|---|---|---|
| IC20 | MTOR | MTOR | 80 |
| IC20 | MUSK | MUSK | 100 |
| IC20 | MYLK | MYLK | 100 |
| IC20 | MYLK2 | MYLK2 | 86 |
| IC20 | MYLK4 | MYLK4 | 85 |
| IC20 | MYO3A | MYO3A | 84 |
| IC20 | MYO3B | MYO3B | 100 |
| IC20 | NDR1 | STK38 | 74 |
| IC20 | NDR2 | STK38L | 52 |
| IC20 | NEK1 | NEK1 | 51 |
| IC20 | NEK10 | NEK10 | 100 |
| IC20 | NEK11 | NEK11 | 100 |
| IC20 | NEK2 | NEK2 | 100 |
| IC20 | NEK3 | NEK3 | 100 |
| IC20 | NEK4 | NEK4 | 100 |
| IC20 | NEK5 | NEK5 | 79 |
| IC20 | NEK6 | NEK6 | 88 |
| IC20 | NEK7 | NEK7 | 81 |
| IC20 | NEK9 | NEK9 | 83 |
| IC20 | NIK | MAP3K14 | 100 |
| IC20 | NIM1 | MGC42105 | 100 |
| IC20 | NLK | NLK | 100 |
| IC20 | OSR1 | OXSR1 | 100 |
| IC20 | p38-alpha | MAPK14 | 100 |
| IC20 | p38-beta | MAPK11 | 100 |
| IC20 | p38-delta | MAPK13 | 100 |
| IC20 | p38-gamma | MAPK12 | 100 |
| IC20 | PAK1 | PAK1 | 89 |
| IC20 | PAK2 | PAK2 | 63 |
| IC20 | PAK3 | PAK3 | 94 |
| IC20 | PAK4 | PAK4 | 100 |
| IC20 | PAK6 | PAK6 | 94 |
| IC20 | PAK7 | PAK7 | 100 |
| IC20 | PCTK1 | CDK16 | 100 |
| IC20 | PCTK2 | CDK17 | 79 |
| IC20 | PCTK3 | CDK18 | 99 |
| IC20 | PDGFRA | PDGFRA | 91 |
| IC20 | PDGFRB | PDGFRB | 88 |
| IC20 | PDPK1 | PDPK1 | 89 |
| IC20 | PFCDPK1(P.falciparum) | CDPK1 | 100 |
| IC20 | PFPK5(P.falciparum) | MAL13P1.279 | 100 |
| IC20 | PFTAIRE2 | CDK15 | 74 |
| IC20 | PFTK1 | CDK14 | 100 |
| IC20 | PHKG1 | PHKG1 | 77 |
| IC20 | PHKG2 | PHKG2 | 100 |
| IC20 | PIK3C2B | PIK3C2B | 76 |
| IC20 | PIK3C2G | PIK3C2G | 100 |

| | | | |
|---|---|---|---|
| IC20 | PIK3CA | PIK3CA | 64 |
| IC20 | PIK3CA(C420R) | PIK3CA | 90 |
| IC20 | PIK3CA(E542K) | PIK3CA | 66 |
| IC20 | PIK3CA(E545A) | PIK3CA | 100 |
| IC20 | PIK3CA(E545K) | PIK3CA | 69 |
| IC20 | PIK3CA(H1047L) | PIK3CA | 51 |
| IC20 | PIK3CA(H1047Y) | PIK3CA | 100 |
| IC20 | PIK3CA(I800L) | PIK3CA | 35 |
| IC20 | PIK3CA(M1043I) | PIK3CA | 75 |
| IC20 | PIK3CA(Q546K) | PIK3CA | 64 |
| IC20 | PIK3CB | PIK3CB | 89 |
| IC20 | PIK3CD | PIK3CD | 70 |
| IC20 | PIK3CG | PIK3CG | 40 |
| IC20 | PIK4CB | PI4KB | 93 |
| IC20 | PIKFYVE | PIKFYVE | 66 |
| IC20 | PIM1 | PIM1 | 60 |
| IC20 | PIM2 | PIM2 | 100 |
| IC20 | PIM3 | PIM3 | 56 |
| IC20 | PIP5K1A | PIP5K1A | 100 |
| IC20 | PIP5K1C | PIP5K1C | 70 |
| IC20 | PIP5K2B | PIP4K2B | 94 |
| IC20 | PIP5K2C | PIP4K2C | 15 |
| IC20 | PKAC-alpha | PRKACA | 91 |
| IC20 | PKAC-beta | PRKACB | 100 |
| IC20 | PKMYT1 | PKMYT1 | 70 |
| IC20 | PKN1 | PKN1 | 100 |
| IC20 | PKN2 | PKN2 | 95 |
| IC20 | PKNB(M.tuberculosis) | pknB | 92 |
| IC20 | PLK1 | PLK1 | 100 |
| IC20 | PLK2 | PLK2 | 100 |
| IC20 | PLK3 | PLK3 | 100 |
| IC20 | PLK4 | PLK4 | 80 |
| IC20 | PRKCD | PRKCD | 72 |
| IC20 | PRKCE | PRKCE | 75 |
| IC20 | PRKCH | PRKCH | 82 |
| IC20 | PRKCI | PRKCI | 100 |
| IC20 | PRKCQ | PRKCQ | 100 |
| IC20 | PRKD1 | PRKD1 | 63 |
| IC20 | PRKD2 | PRKD2 | 72 |
| IC20 | PRKD3 | PRKD3 | 100 |
| IC20 | PRKG1 | PRKG1 | 100 |
| IC20 | PRKG2 | PRKG2 | 84 |
| IC20 | PRKR | EIF2AK2 | 50 |
| IC20 | PRKX | PRKX | 95 |
| IC20 | PRP4 | PRPF4B | 67 |
| IC20 | PYK2 | PTK2B | 88 |
| IC20 | QSK | KIAA0999 | 72 |

| | | | |
|---|---|---|---|
| IC20 | RAF1 | RAF1 | 85 |
| IC20 | RET | RET | 100 |
| IC20 | RET(M918T) | RET | 100 |
| IC20 | RET(V804L) | RET | 98 |
| IC20 | RET(V804M) | RET | 85 |
| IC20 | RIOK1 | RIOK1 | 73 |
| IC20 | RIOK2 | RIOK2 | 90 |
| IC20 | RIOK3 | RIOK3 | 95 |
| IC20 | RIPK1 | RIPK1 | 100 |
| IC20 | RIPK2 | RIPK2 | 56 |
| IC20 | RIPK4 | RIPK4 | 100 |
| IC20 | RIPK5 | DSTYK | 100 |
| IC20 | ROCK1 | ROCK1 | 100 |
| IC20 | ROCK2 | ROCK2 | 100 |
| IC20 | ROS1 | ROS1 | 100 |
| IC20 | RPS6KA4(Kin.Dom.1-N-terminal) | RPS6KA4 | 87 |
| IC20 | RPS6KA4(Kin.Dom.2-C-terminal) | RPS6KA4 | 39 |
| IC20 | RPS6KA5(Kin.Dom.1-N-terminal) | RPS6KA5 | 81 |
| IC20 | RPS6KA5(Kin.Dom.2-C-terminal) | RPS6KA5 | 90 |
| IC20 | RSK1(Kin.Dom.1-N-terminal) | RPS6KA1 | 89 |
| IC20 | RSK1(Kin.Dom.2-C-terminal) | RPS6KA1 | 100 |
| IC20 | RSK2(Kin.Dom.1-N-terminal) | RPS6KA3 | 99 |
| IC20 | RSK2(Kin.Dom.2-C-terminal) | RPS6KA3 | 88 |
| IC20 | RSK3(Kin.Dom.1-N-terminal) | RPS6KA2 | 100 |
| IC20 | RSK3(Kin.Dom.2-C-terminal) | RPS6KA2 | 93 |
| IC20 | RSK4(Kin.Dom.1-N-terminal) | RPS6KA6 | 82 |
| IC20 | RSK4(Kin.Dom.2-C-terminal) | RPS6KA6 | 89 |
| IC20 | S6K1 | RPS6KB1 | 80 |
| IC20 | SBK1 | SBK1 | 95 |
| IC20 | SGK | SGK1 | 100 |
| IC20 | SgK110 | SgK110 | 92 |
| IC20 | SGK2 | SGK2 | 78 |
| IC20 | SGK3 | SGK3 | 72 |
| IC20 | SIK | SIK1 | 98 |
| IC20 | SIK2 | SIK2 | 100 |
| IC20 | SLK | SLK | 84 |
| IC20 | SNARK | NUAK2 | 88 |
| IC20 | SNRK | SNRK | 79 |
| IC20 | SRC | SRC | 93 |
| IC20 | SRMS | SRMS | 93 |
| IC20 | SRPK1 | SRPK1 | 93 |
| IC20 | SRPK2 | SRPK2 | 83 |
| IC20 | SRPK3 | SRPK3 | 100 |
| IC20 | STK16 | STK16 | 84 |
| IC20 | STK33 | STK33 | 97 |
| IC20 | STK35 | STK35 | 100 |
| IC20 | STK36 | STK36 | 99 |

| | | | |
|---|---|---|---|
| IC20 | STK39 | STK39 | 100 |
| IC20 | SYK | SYK | 87 |
| IC20 | TAK1 | MAP3K7 | 81 |
| IC20 | TAOK1 | TAOK1 | 84 |
| IC20 | TAOK2 | TAOK2 | 81 |
| IC20 | TAOK3 | TAOK3 | 82 |
| IC20 | TBK1 | TBK1 | 68 |
| IC20 | TEC | TEC | 100 |
| IC20 | TESK1 | TESK1 | 100 |
| IC20 | TGFBR1 | TGFBR1 | 88 |
| IC20 | TGFBR2 | TGFBR2 | 64 |
| IC20 | TIE1 | TIE1 | 100 |
| IC20 | TIE2 | TEK | 77 |
| IC20 | TLK1 | TLK1 | 88 |
| IC20 | TLK2 | TLK2 | 90 |
| IC20 | TNIK | TNIK | 100 |
| IC20 | TNK1 | TNK1 | 99 |
| IC20 | TNK2 | TNK2 | 100 |
| IC20 | TNNI3K | TNNI3K | 57 |
| IC20 | TRKA | NTRK1 | 92 |
| IC20 | TRKB | NTRK2 | 93 |
| IC20 | TRKC | NTRK3 | 83 |
| IC20 | TRPM6 | TRPM6 | 84 |
| IC20 | TSSK1B | TSSK1B | 100 |
| IC20 | TSSK3 | TSSK3 | 91 |
| IC20 | TTK | TTK | 85 |
| IC20 | TXK | TXK | 81 |
| IC20 | TYK2(JH1domain-catalytic) | TYK2 | 100 |
| IC20 | TYK2(JH2domain-pseudokinase) | TYK2 | 100 |
| IC20 | TYRO3 | TYRO3 | 97 |
| IC20 | ULK1 | ULK1 | 100 |
| IC20 | ULK2 | ULK2 | 100 |
| IC20 | ULK3 | ULK3 | 100 |
| IC20 | VEGFR2 | KDR | 82 |
| IC20 | VPS34 | PIK3C3 | 100 |
| IC20 | VRK2 | VRK2 | 100 |
| IC20 | WEE1 | WEE1 | 88 |
| IC20 | WEE2 | WEE2 | 81 |
| IC20 | WNK1 | WNK1 | 100 |
| IC20 | WNK2 | WNK2 | 100 |
| IC20 | WNK3 | WNK3 | 100 |
| IC20 | WNK4 | WNK4 | 100 |
| IC20 | YANK1 | STK32A | 76 |
| IC20 | YANK2 | STK32B | 58 |
| IC20 | YANK3 | STK32C | 100 |
| IC20 | YES | YES1 | 100 |
| IC20 | YSK1 | STK25 | 100 |

| | | | |
|---|---|---|---|
| IC20 | YSK4 | MAP3K19 | 85 |
| IC20 | ZAK | ZAK | 98 |
| IC20 | ZAP70 | ZAP70 | 90 |

**Figure S1**: $^{1}$H and $^{13}$C NMR of compounds **3–8**; **10–12**; **14**; **16** and **18–37**.

#### Compound 3

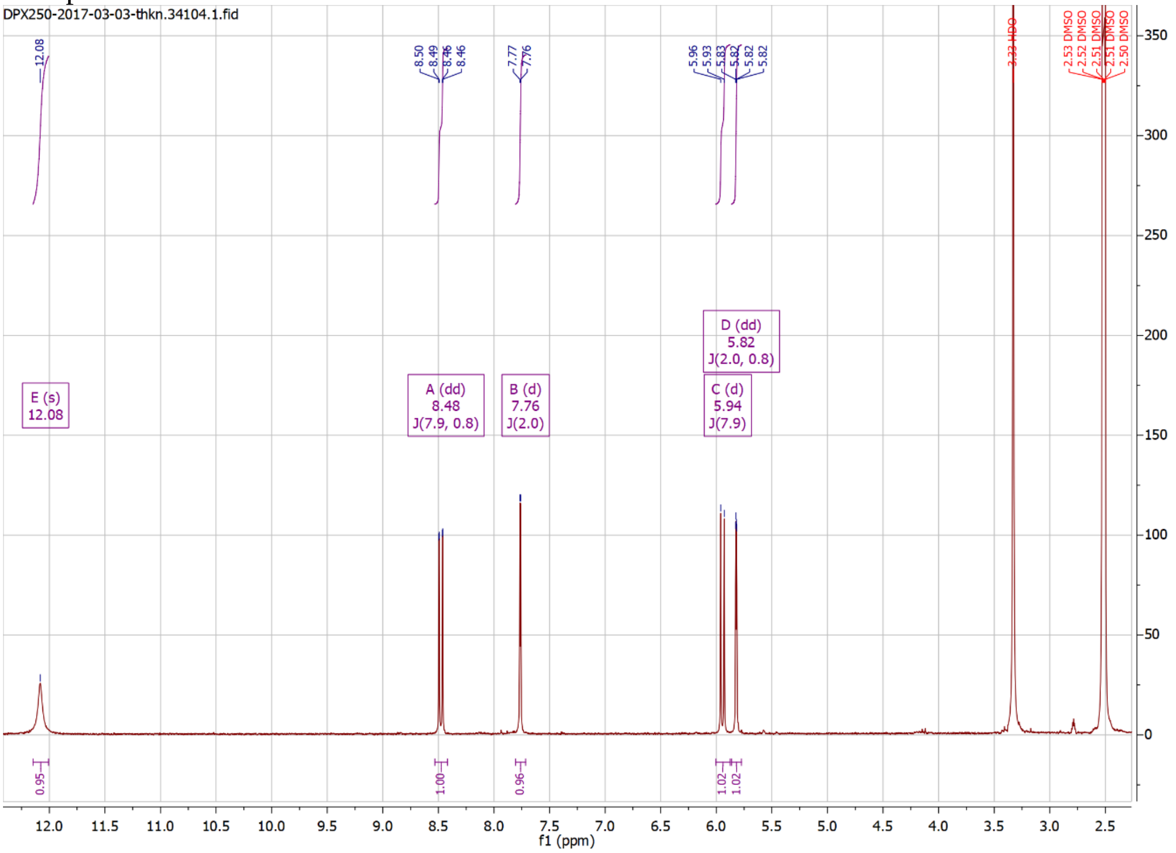

#### Compound 4

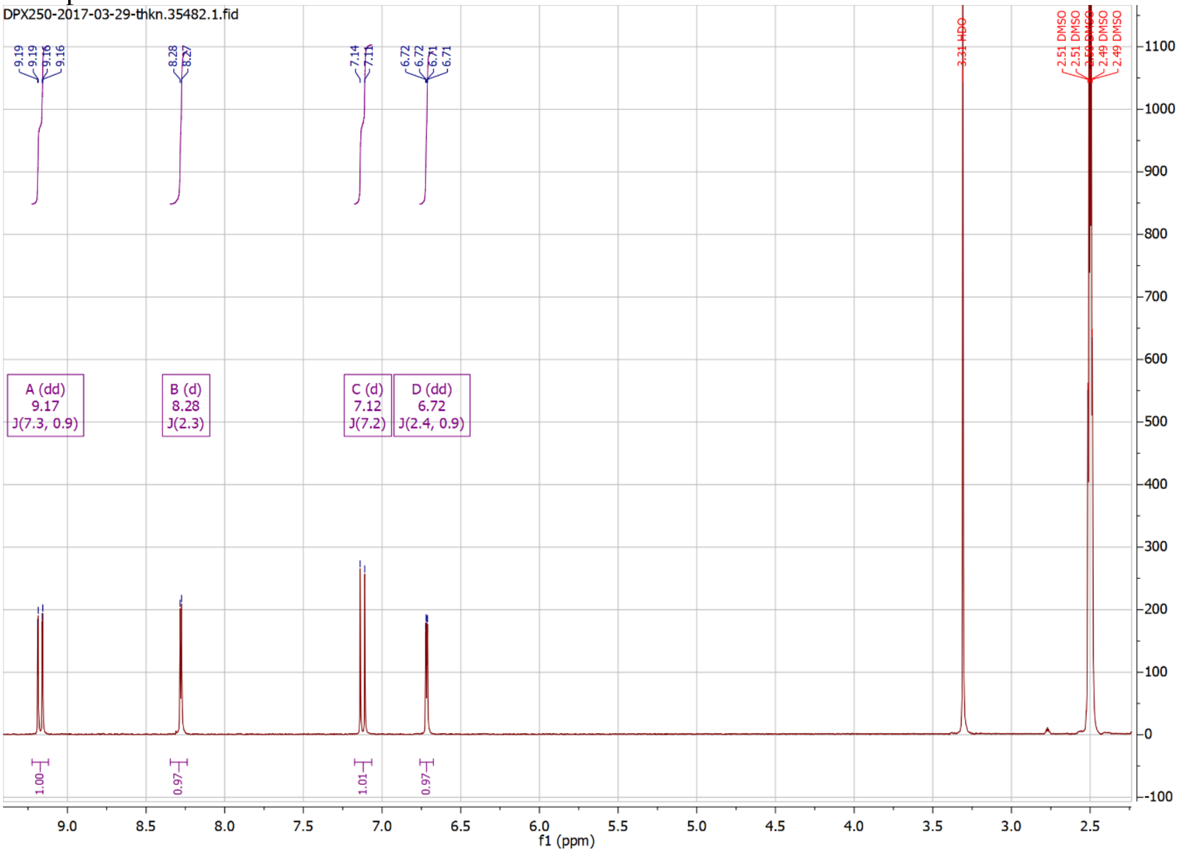

Compound 5

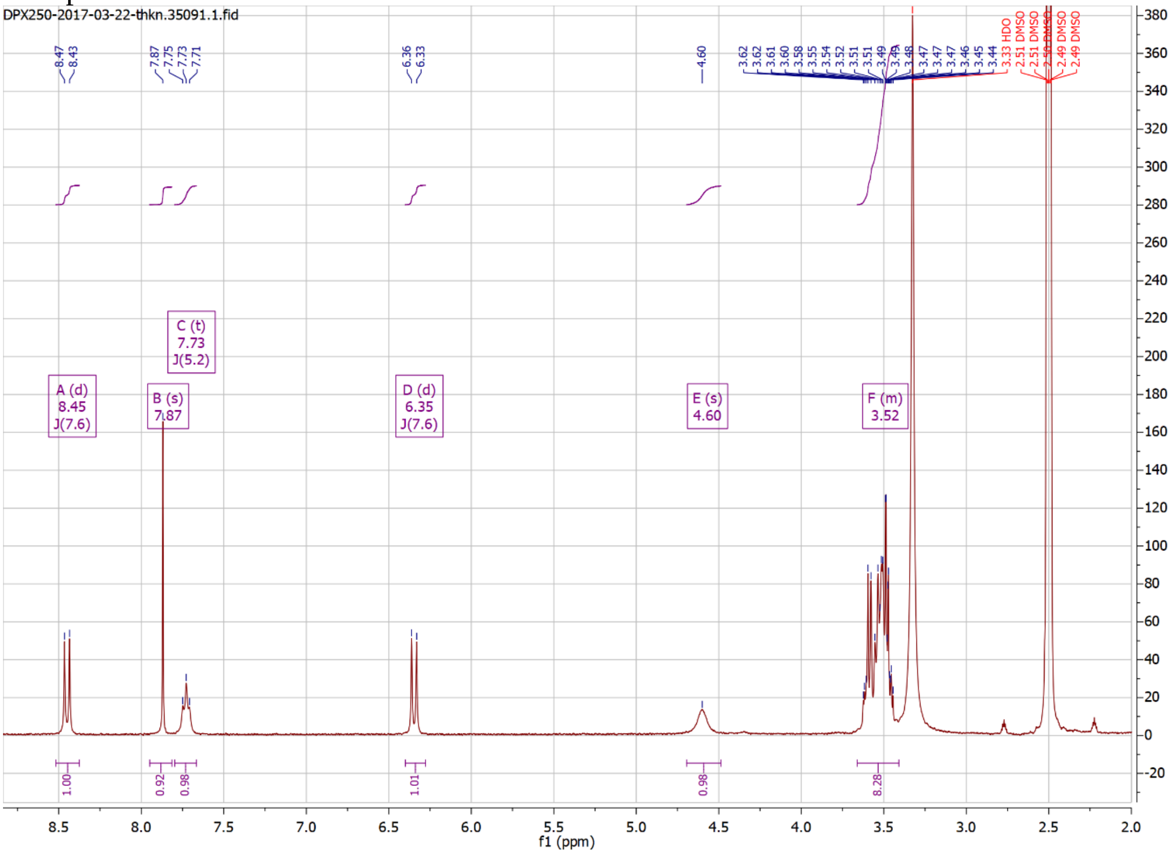

Compound 6

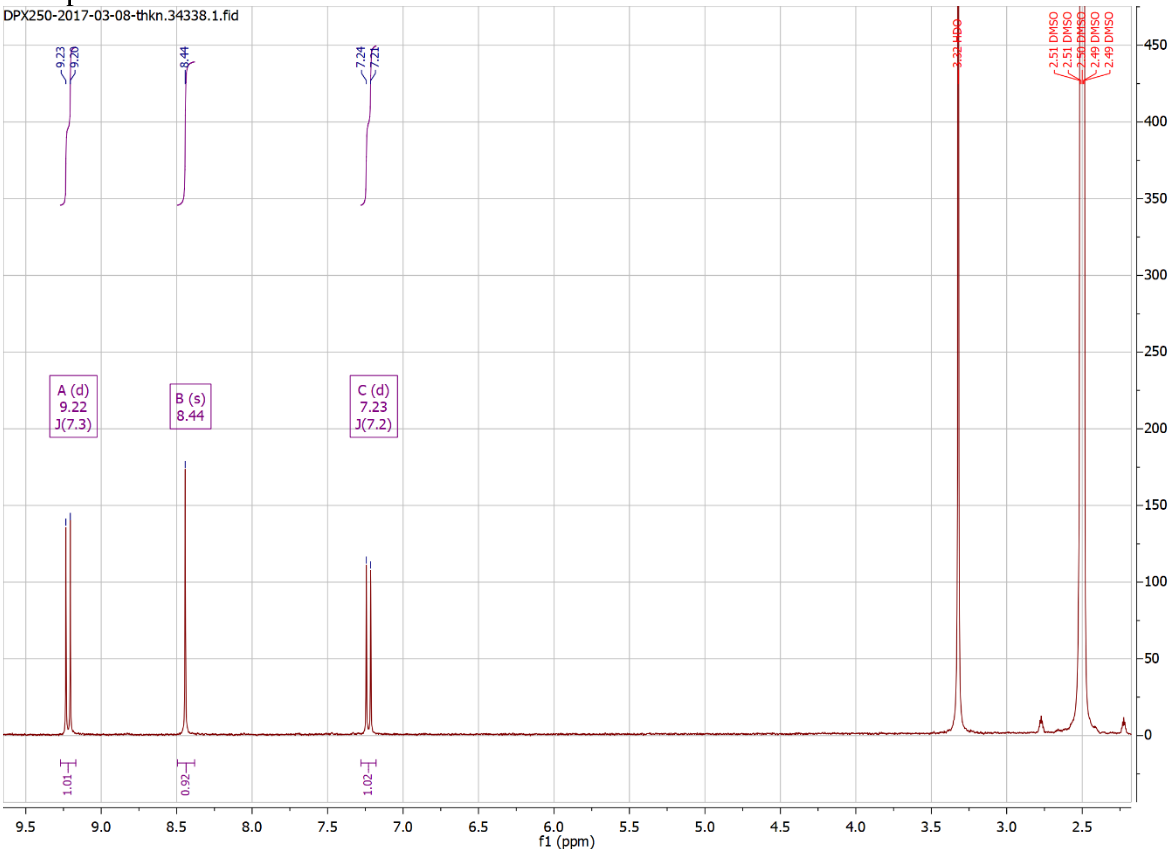

Compound 7

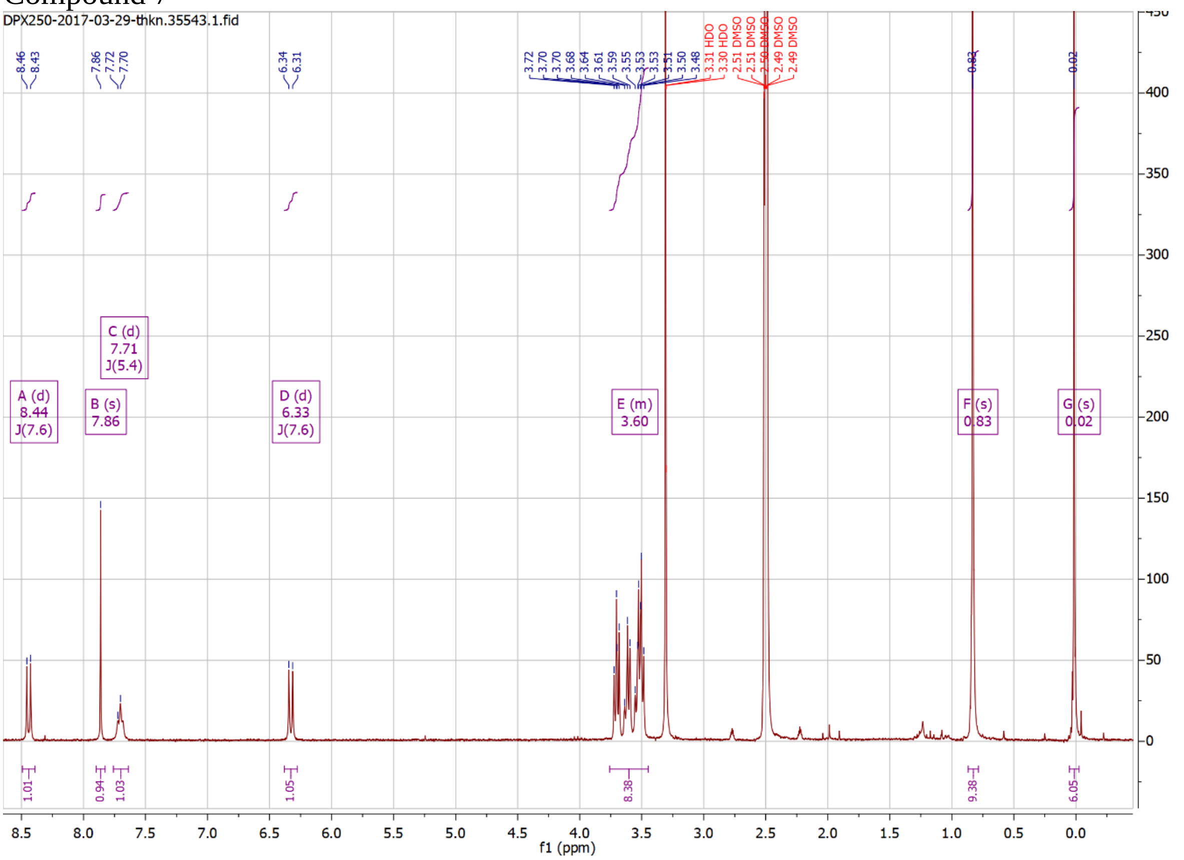

Compound 8

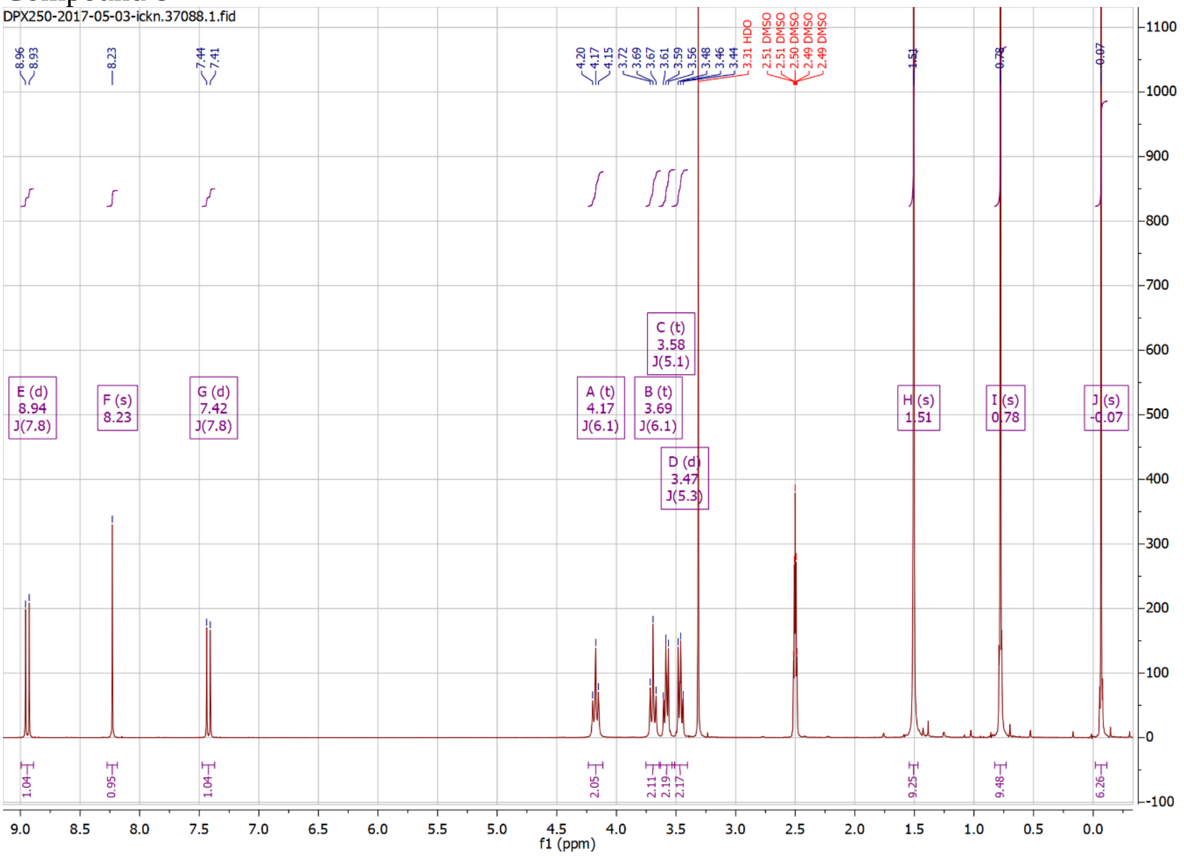

Compound 10

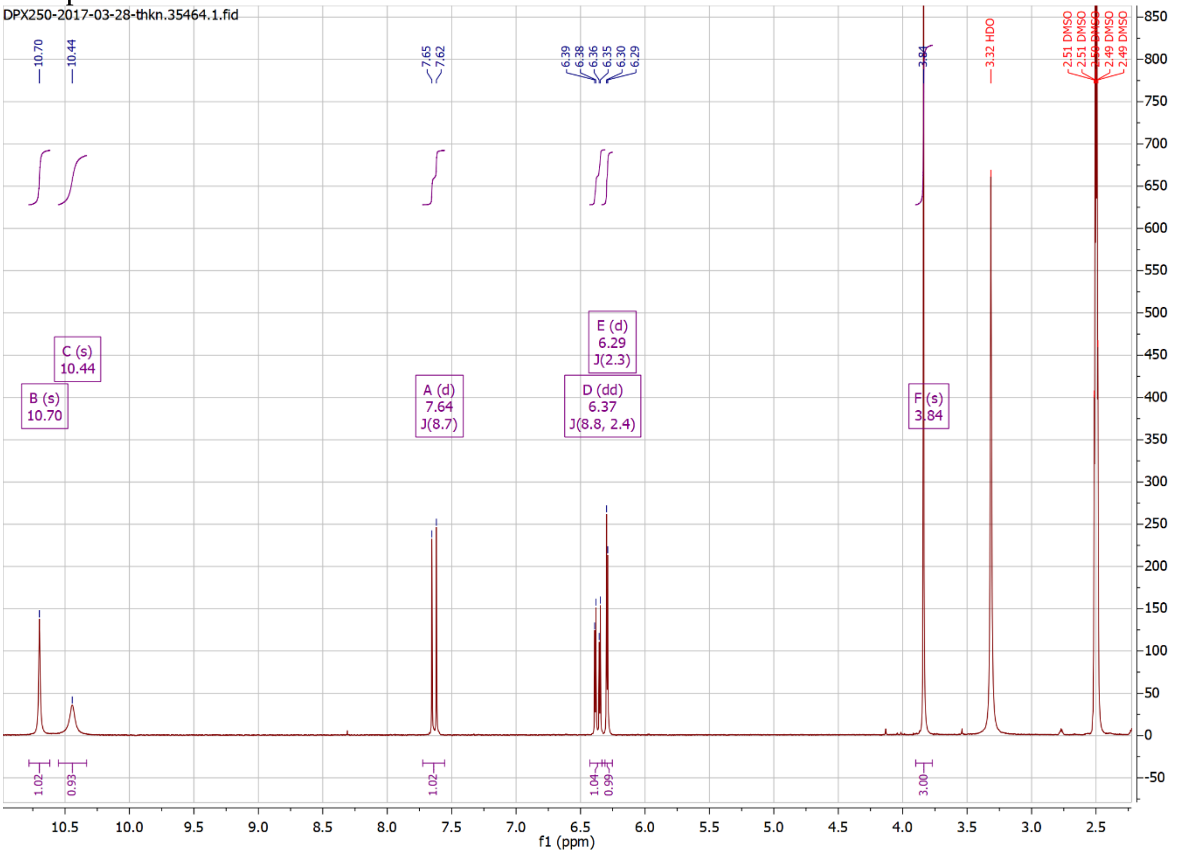

Compound 11

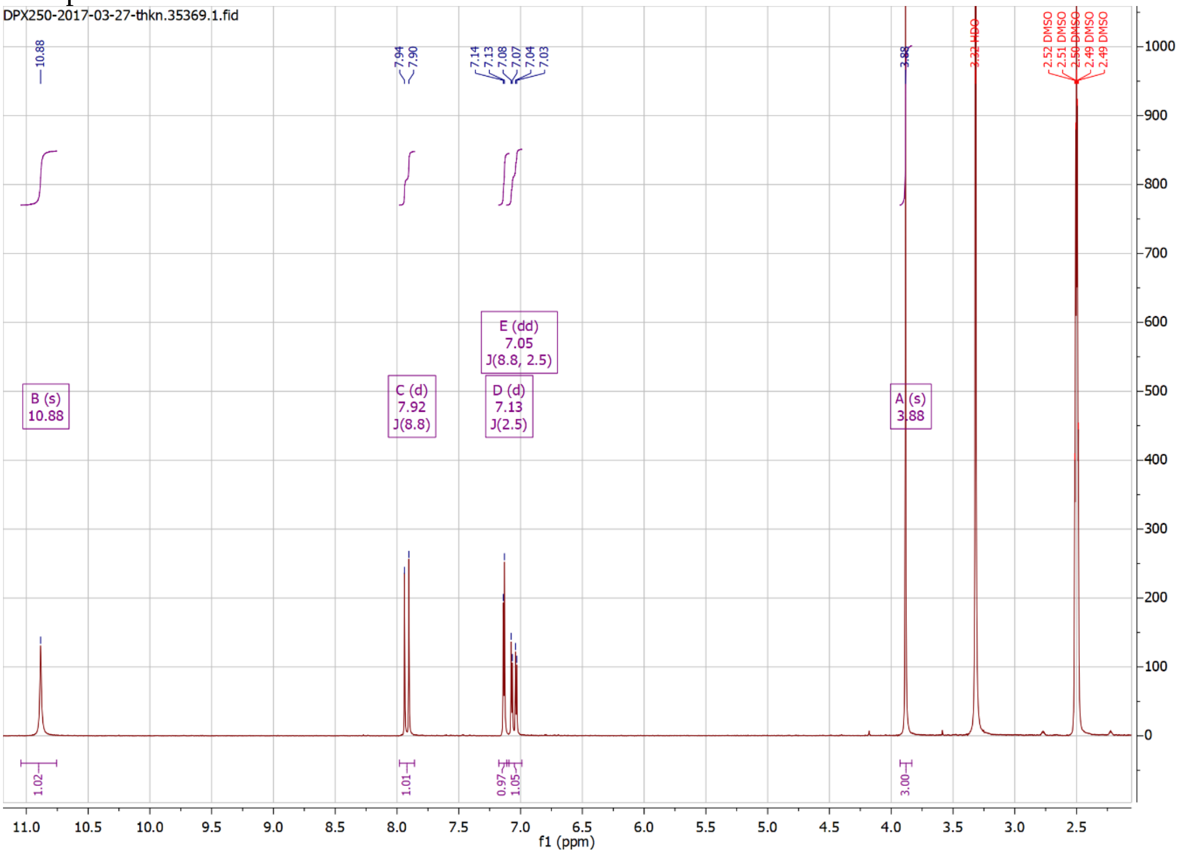

### Compound 12

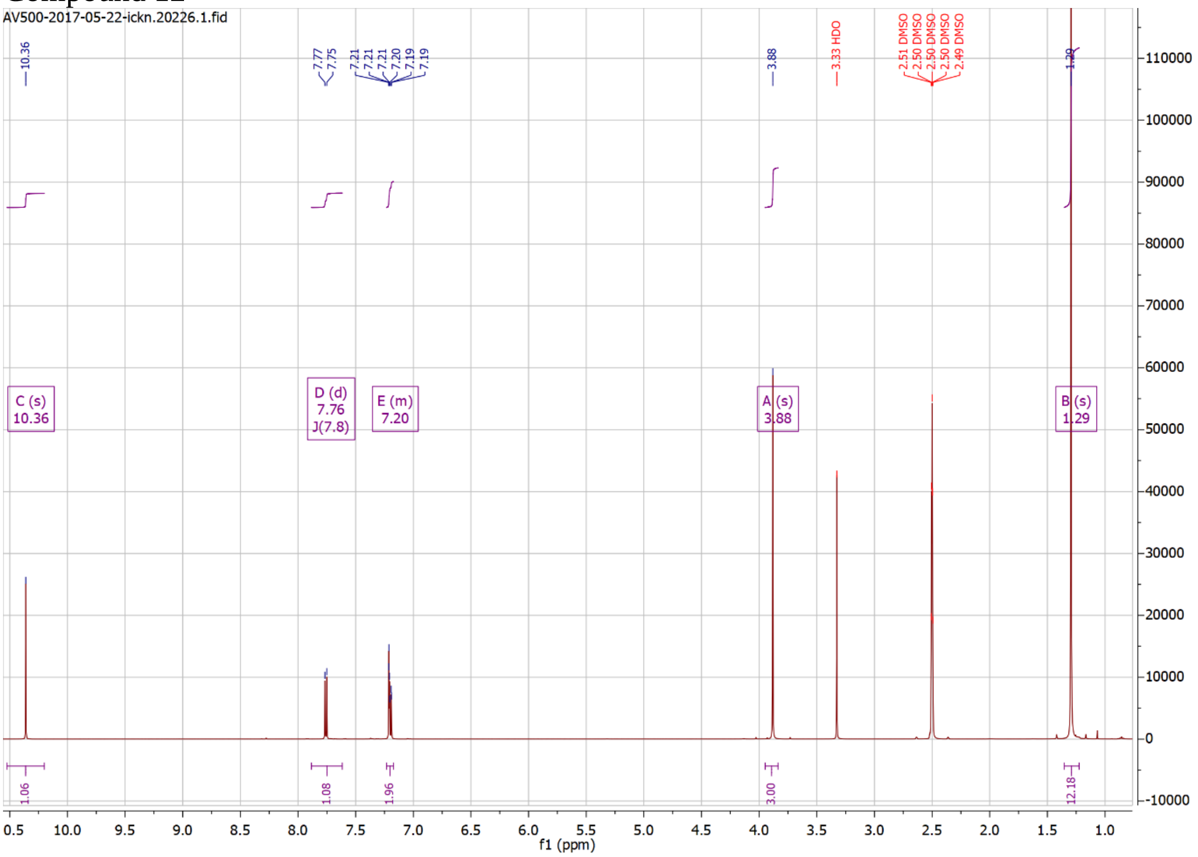

AV500-2017-05-22-ickn.20226.1.fid
10.36
7.77
7.75
7.21
7.21
7.21
7.20
7.19
7.19
3.88
3.33 HDO
2.51 DMSO
2.50 DMSO
2.50 DMSO
2.50 DMSO
2.49 DMSO
1.29
C (s) 10.36
D (d) 7.76 J(7.8)
E (m) 7.20
A (s) 3.88
B (s) 1.29
1.06
1.08
1.96
3.00
12.18
f1 (ppm)

#### Compound 14

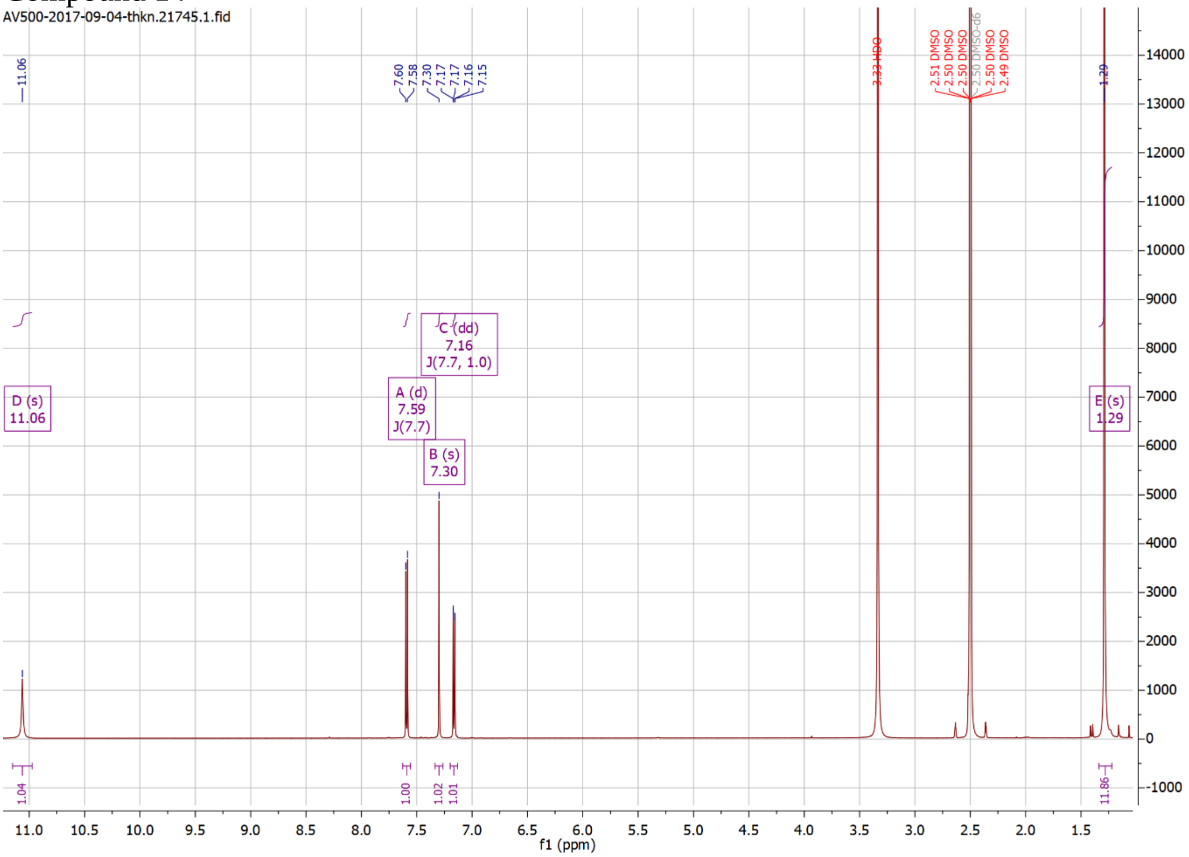

#### Compound 16

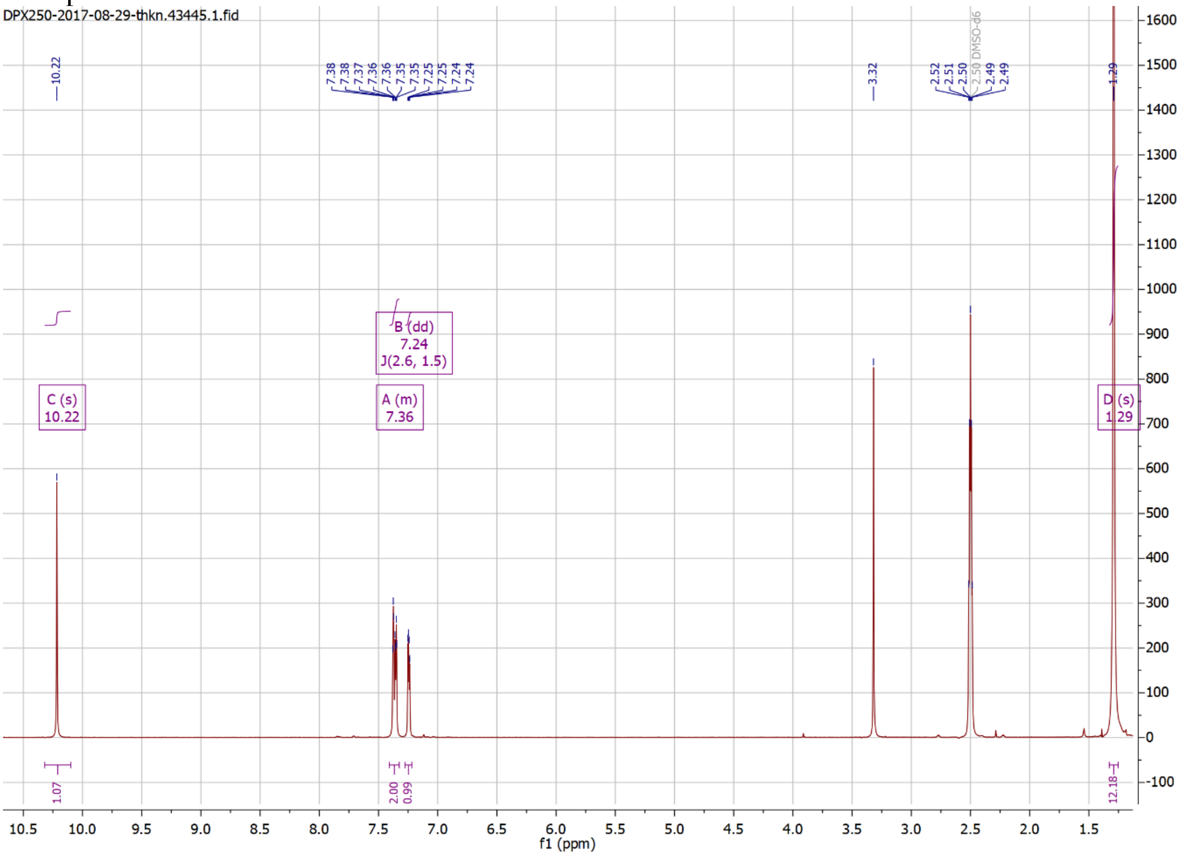

#### Compound 18

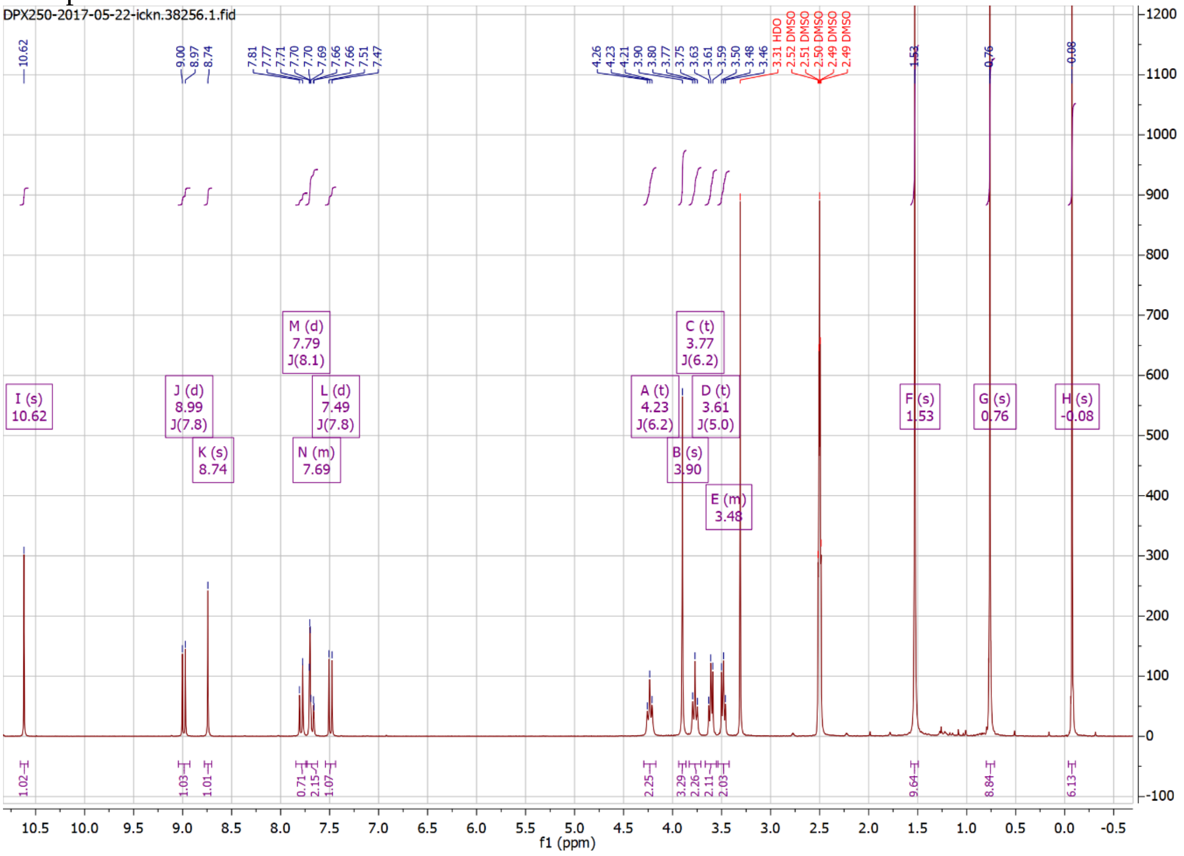

#### Compound 19

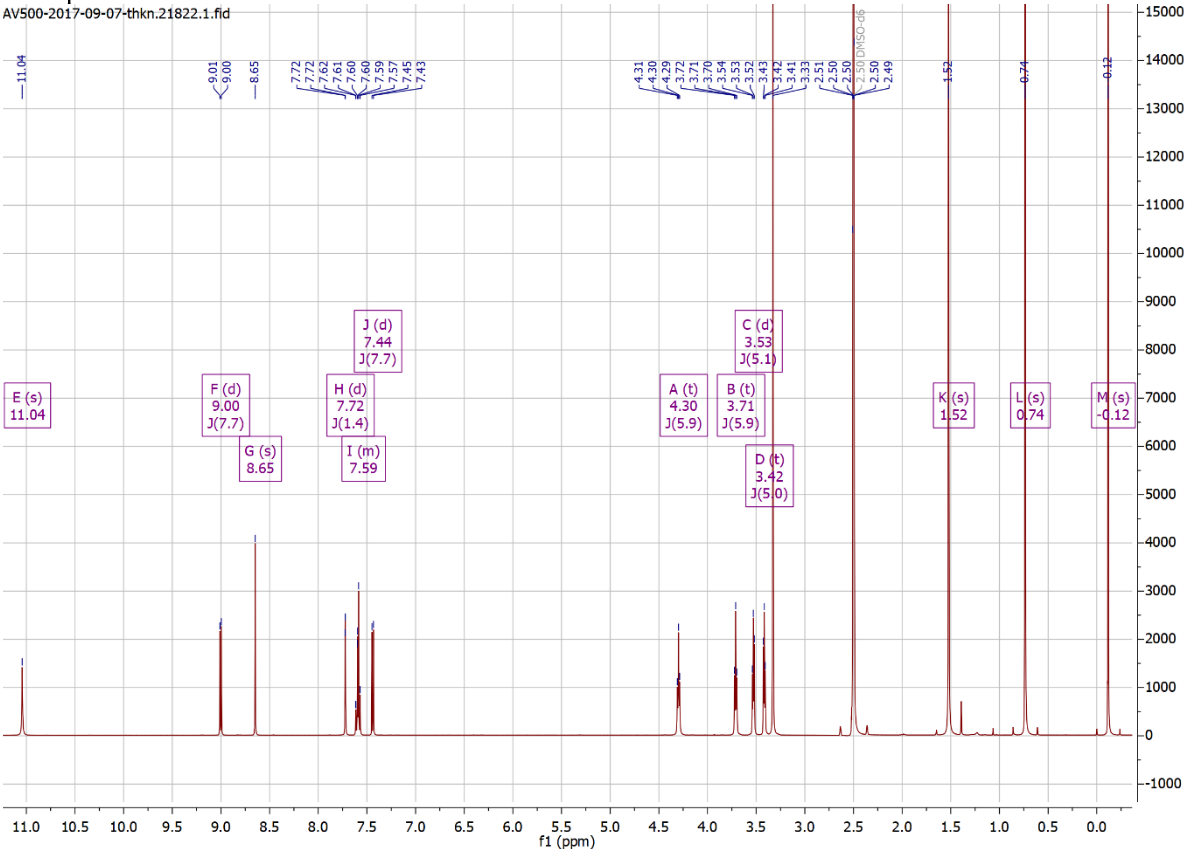

### Compound 20

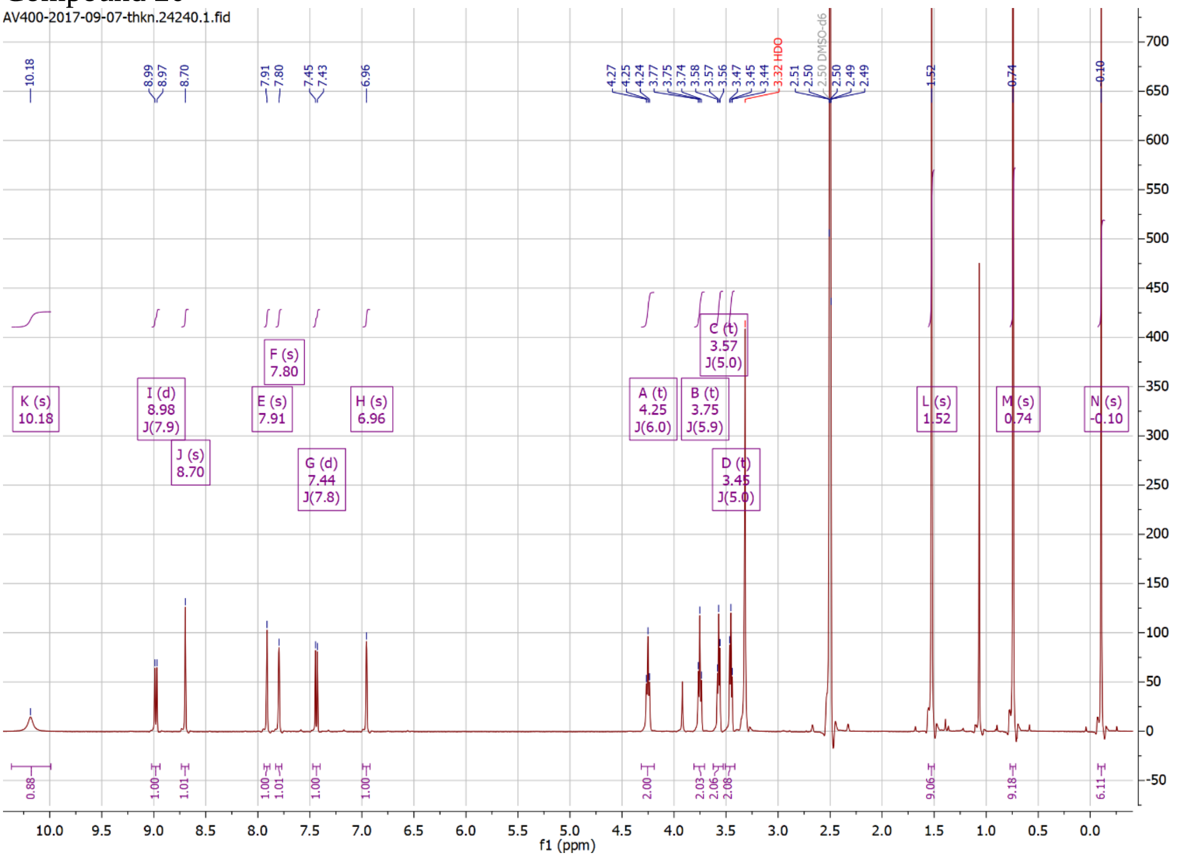

### Compound 21

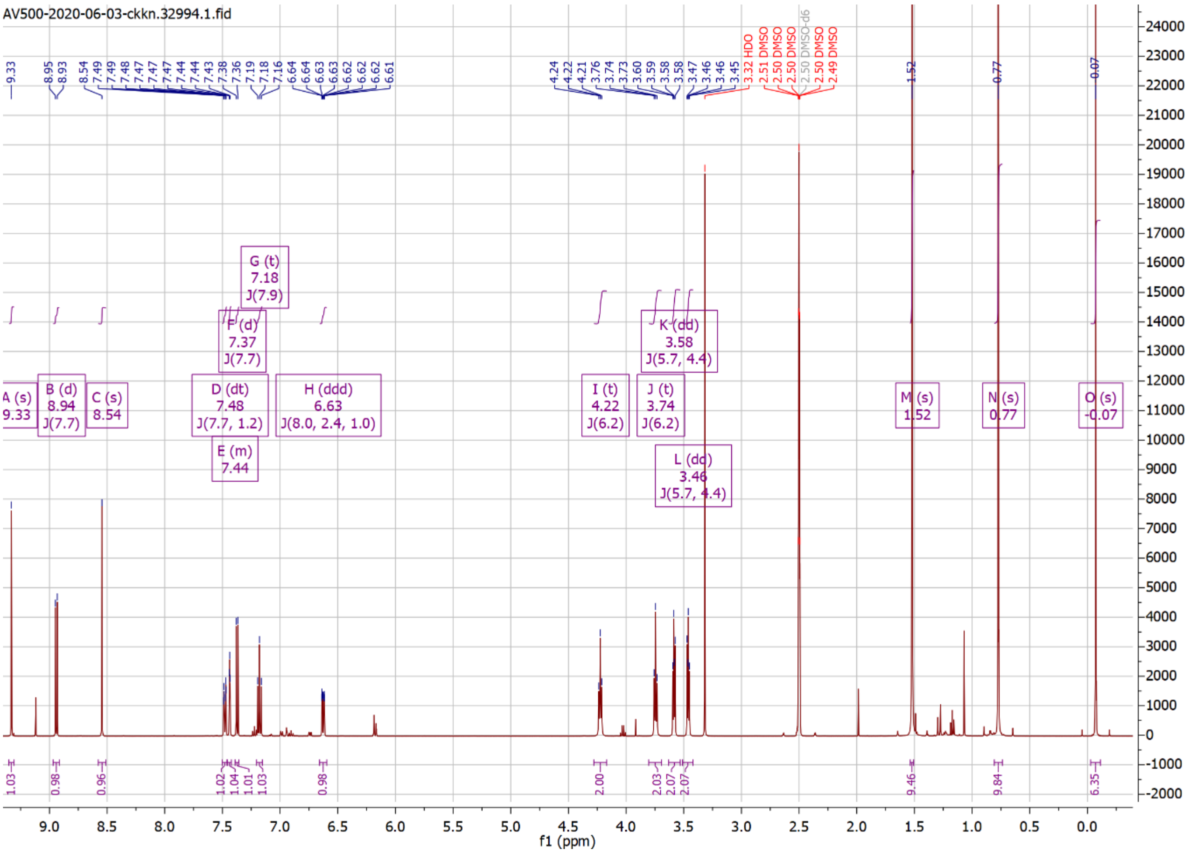

Compound 22

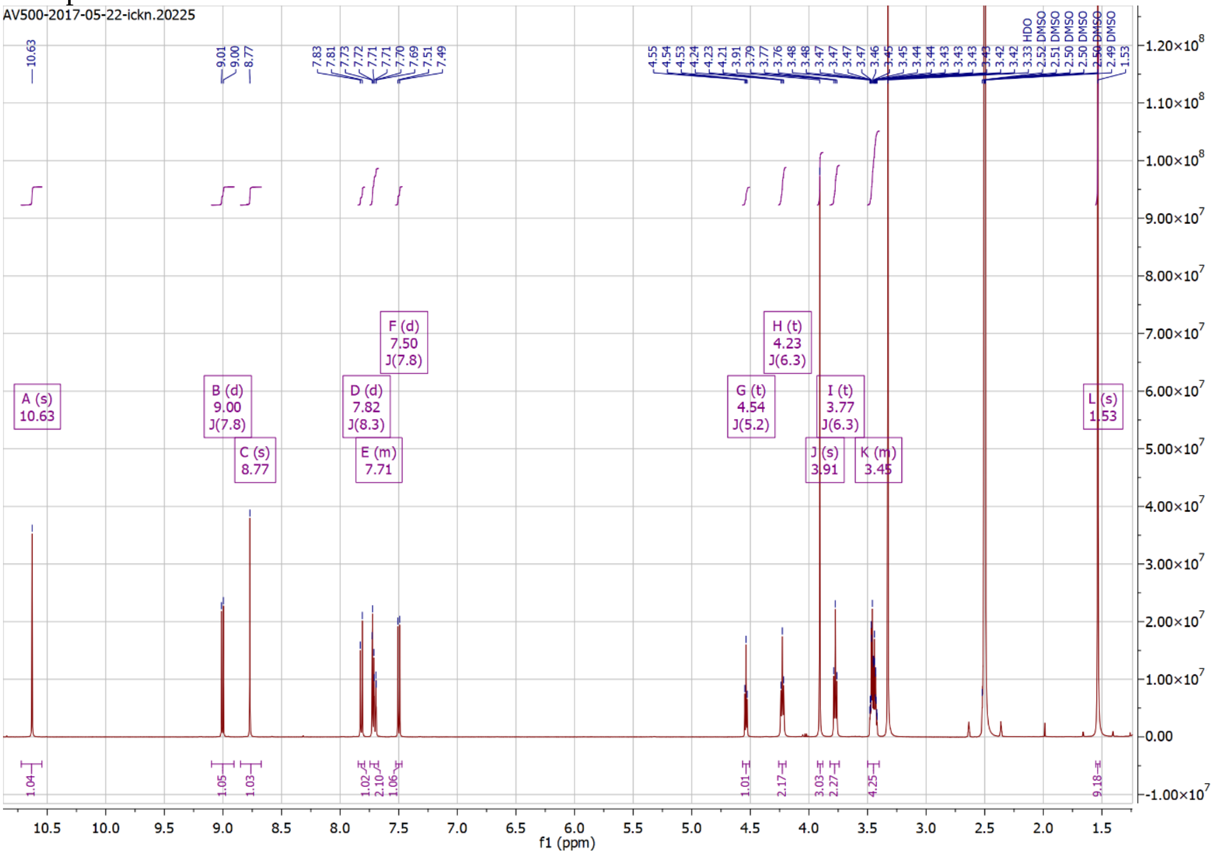
AV500-2017-05-22-ickn.20225
A (s) 10.63
B (d) 9.00 J(7.8)
C (s) 8.77
D (d) 7.82 J(8.3)
E (m) 7.71
F (d) 7.50 J(7.8)
G (t) 4.54 J(5.2)
H (t) 4.23 J(6.3)
I (t) 3.77 J(6.3)
J (s) 3.91
K (m) 3.45
L (s) 1.53
f1 (ppm)

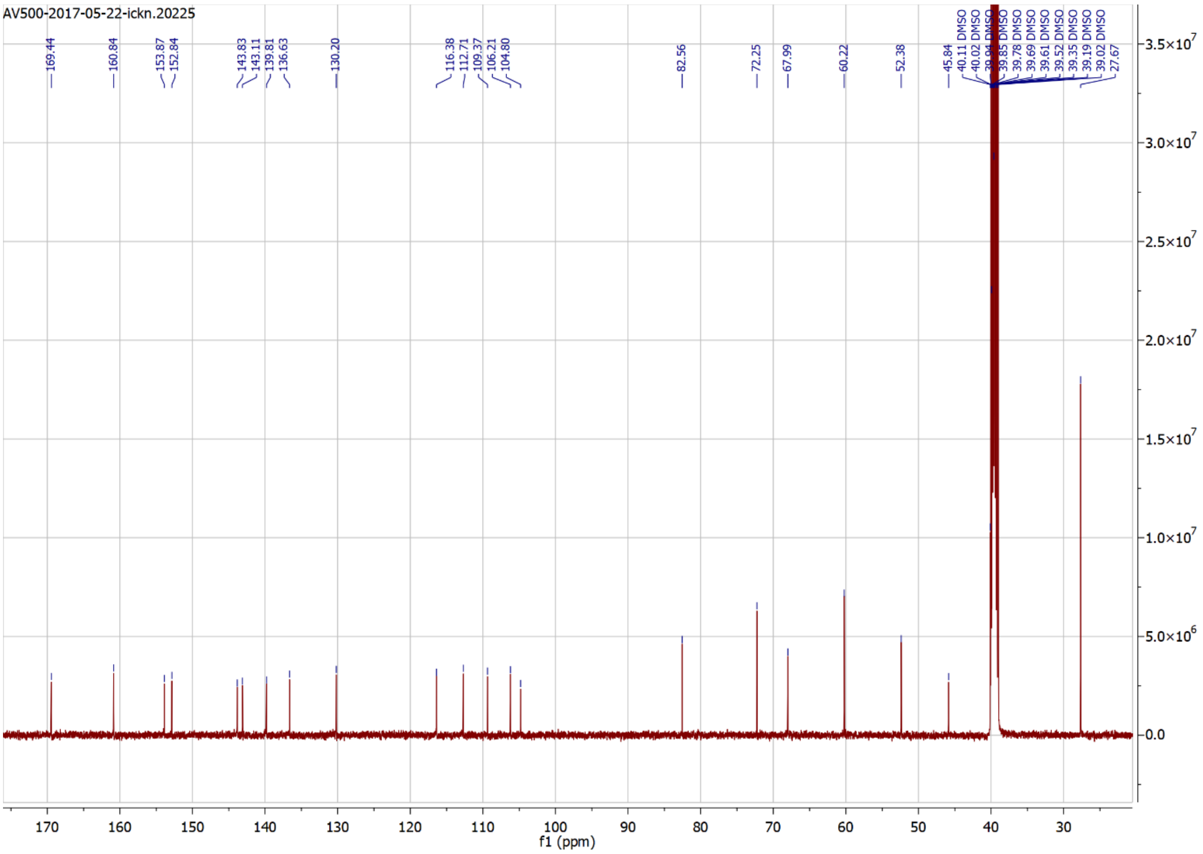
AV500-2017-05-22-ickn.20225
169.44
160.84
153.87
152.84
143.83
143.11
139.81
136.63
130.20
116.38
112.71
109.37
106.21
104.80
82.56
72.25
67.99
60.22
52.38
45.84
27.67
f1 (ppm)

### Compound 23

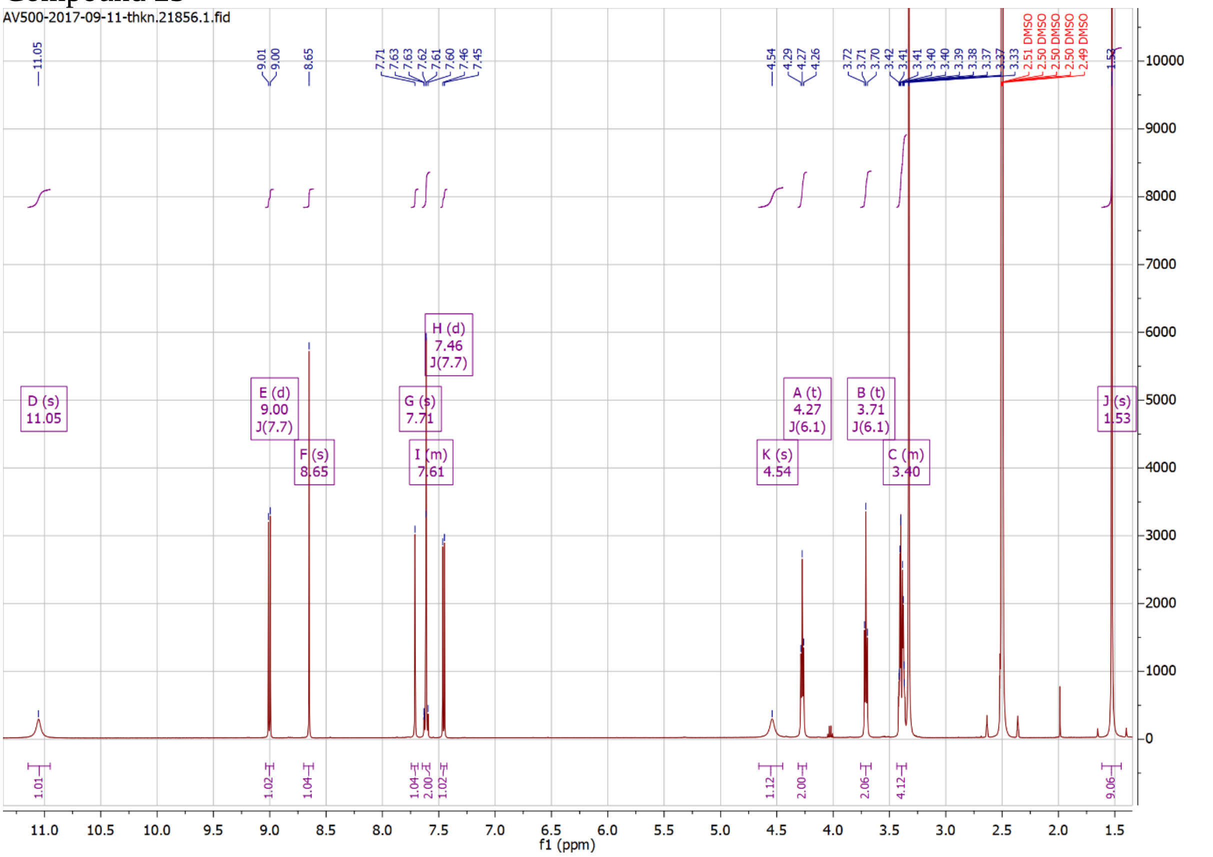
AV500-2017-09-11-thkn.21856.1.fid
f1 (ppm)

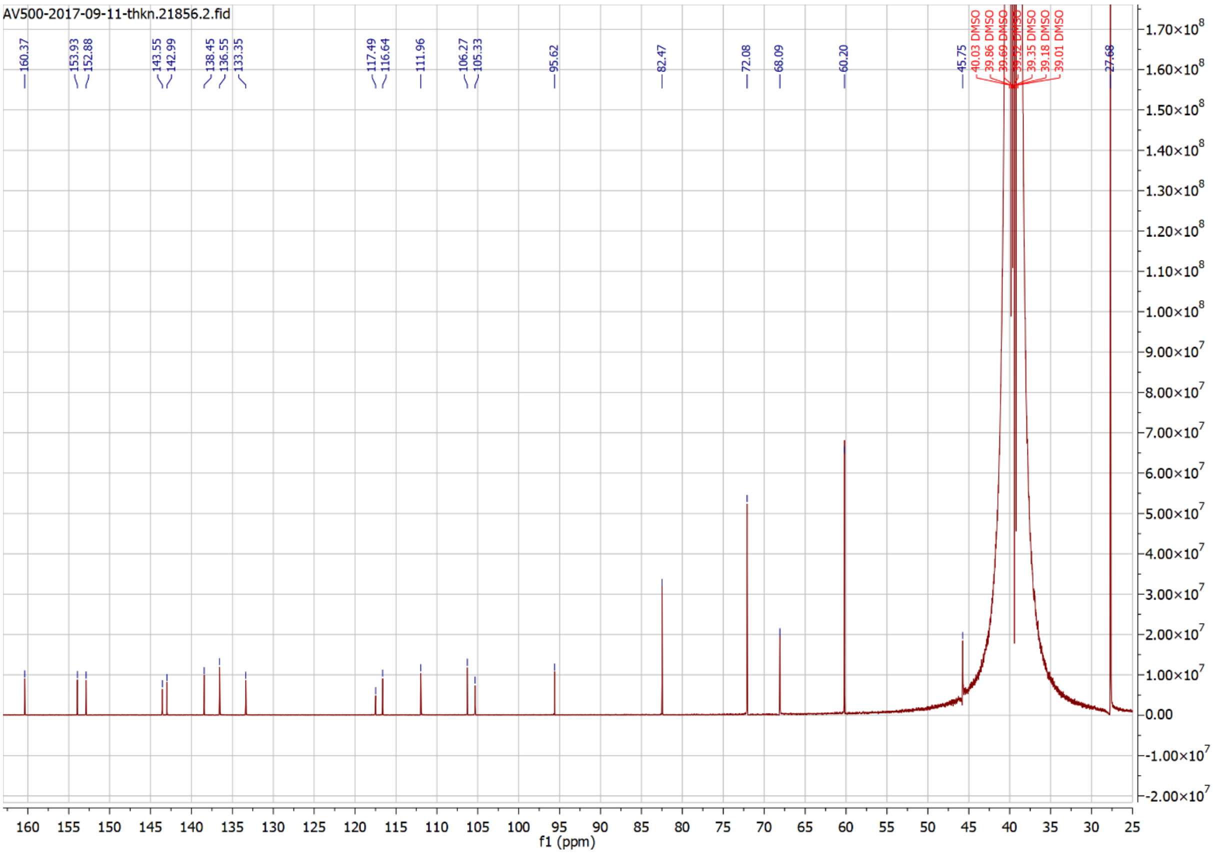
AV500-2017-09-11-thkn.21856.2.fid
f1 (ppm)

Compound 24

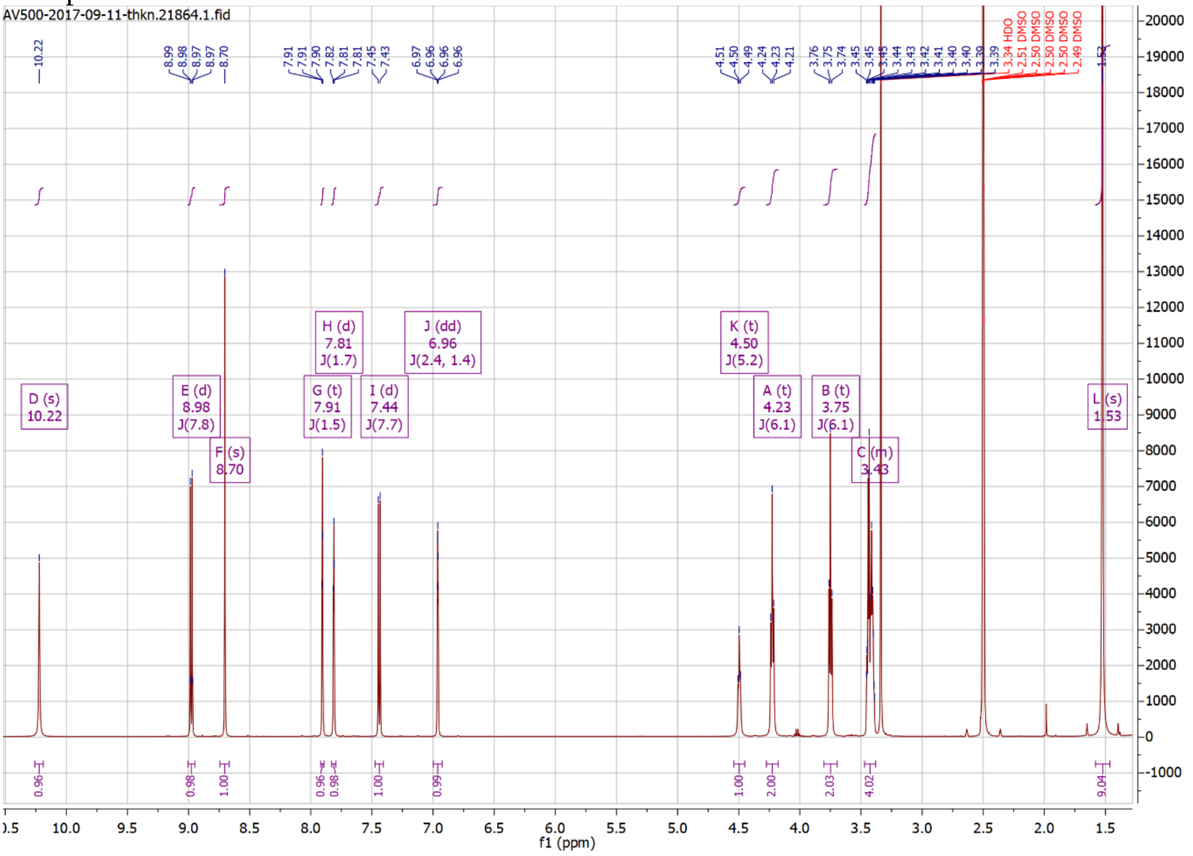
AV500-2017-09-11-thkn.21864.1.fid
D (s) 10.22
E (d) 8.98 J(7.8)
F (s) 8.70
G (t) 7.91 J(1.5)
H (d) 7.81 J(1.7)
I (d) 7.44 J(7.7)
J (dd) 6.96 J(2.4, 1.4)
K (t) 4.50 J(5.2)
A (t) 4.23 J(6.1)
B (t) 3.75 J(6.1)
C (m) 3.43
L (s) 1.53
3.34 HDO
2.51 DMSO
2.50 DMSO
2.50 DMSO
2.50 DMSO
2.49 DMSO
f1 (ppm)

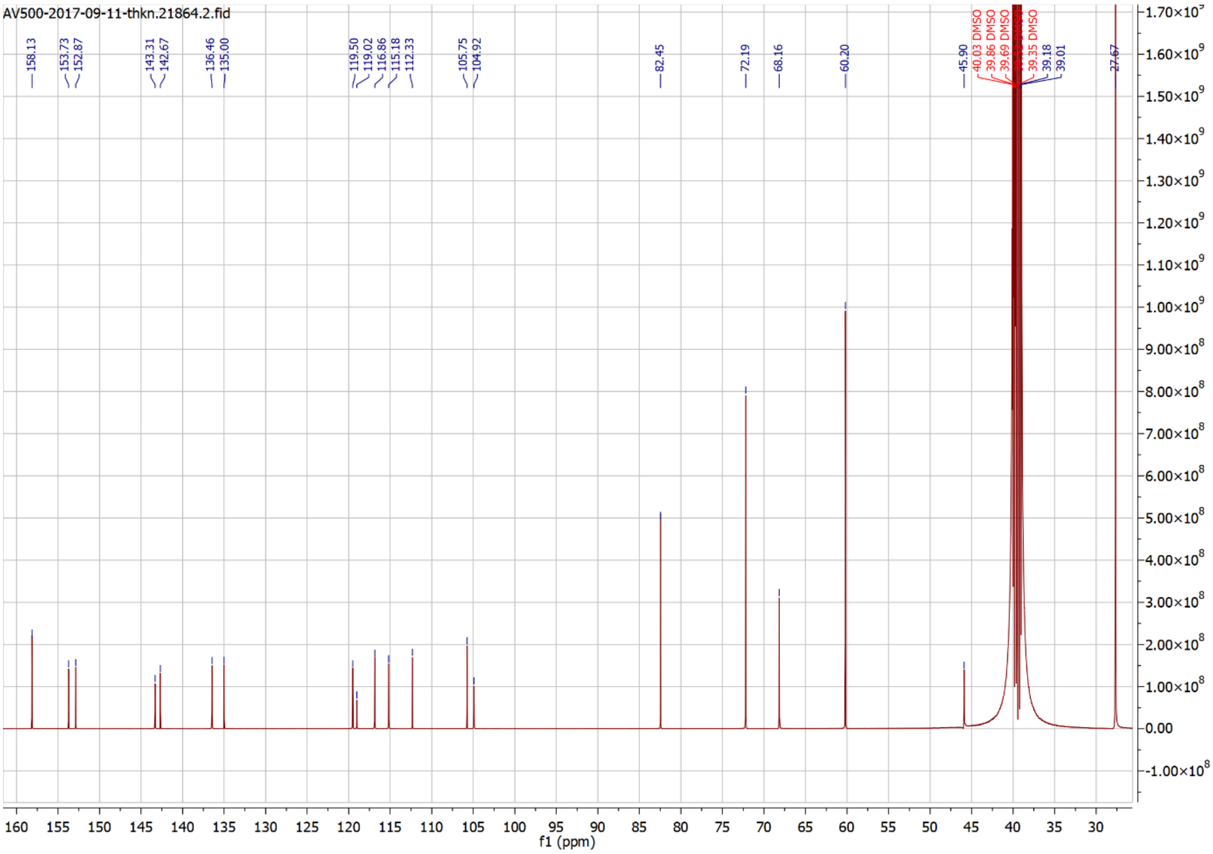
AV500-2017-09-11-thkn.21864.2.fid
158.13
153.73
152.87
143.31
142.67
136.46
135.00
119.50
119.02
116.86
115.18
112.33
105.75
104.92
82.45
72.19
68.16
60.20
45.90
40.03 DMSO
39.86 DMSO
39.69 DMSO
39.35 DMSO
39.18
39.01
27.67
f1 (ppm)

### Compound 25

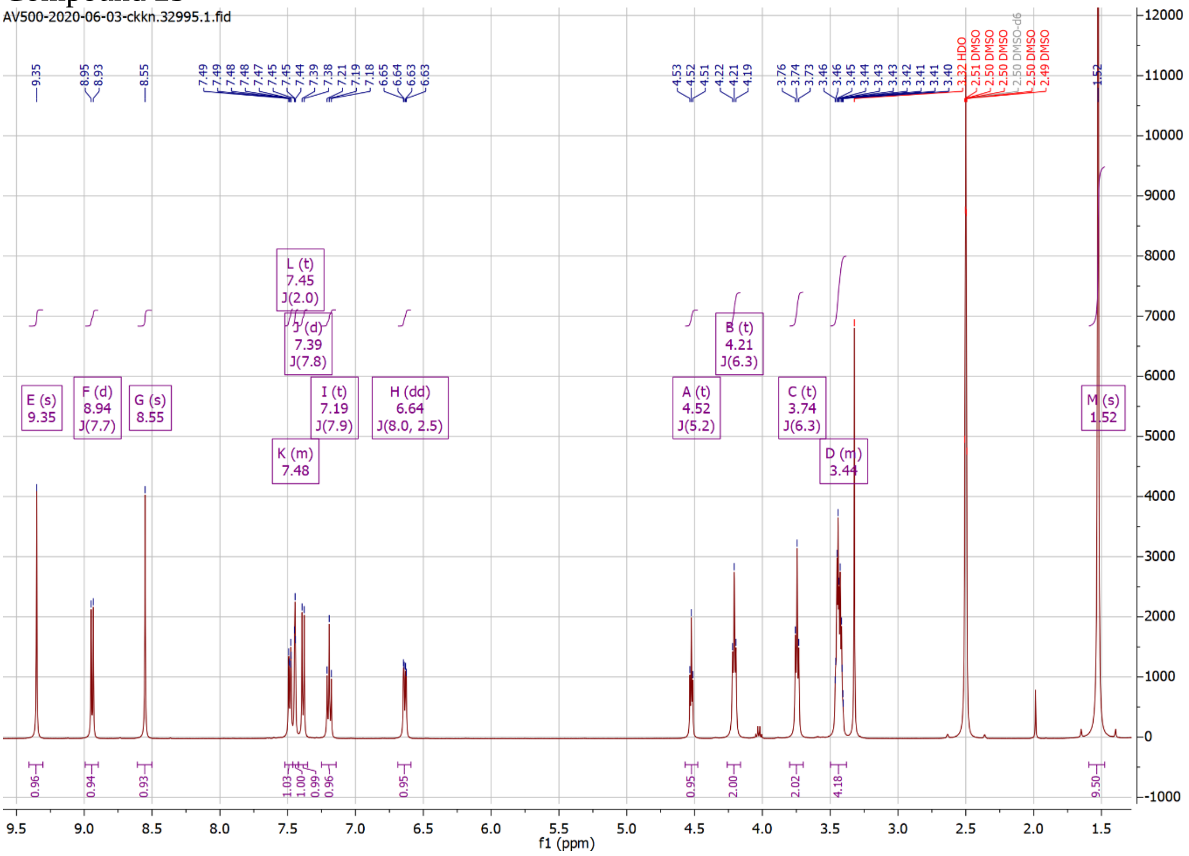
AV500-2020-06-03-ckkn.32995.1.fid
E (s) 9.35
F (d) 8.94 J(7.7)
G (s) 8.55
L (t) 7.45 J(2.0)
J (d) 7.39 J(7.8)
K (m) 7.48
I (t) 7.19 J(7.9)
H (dd) 6.64 J(8.0, 2.5)
A (t) 4.52 J(5.2)
B (t) 4.21 J(6.3)
C (t) 3.74 J(6.3)
D (m) 3.44
M (s) 1.52
f1 (ppm)

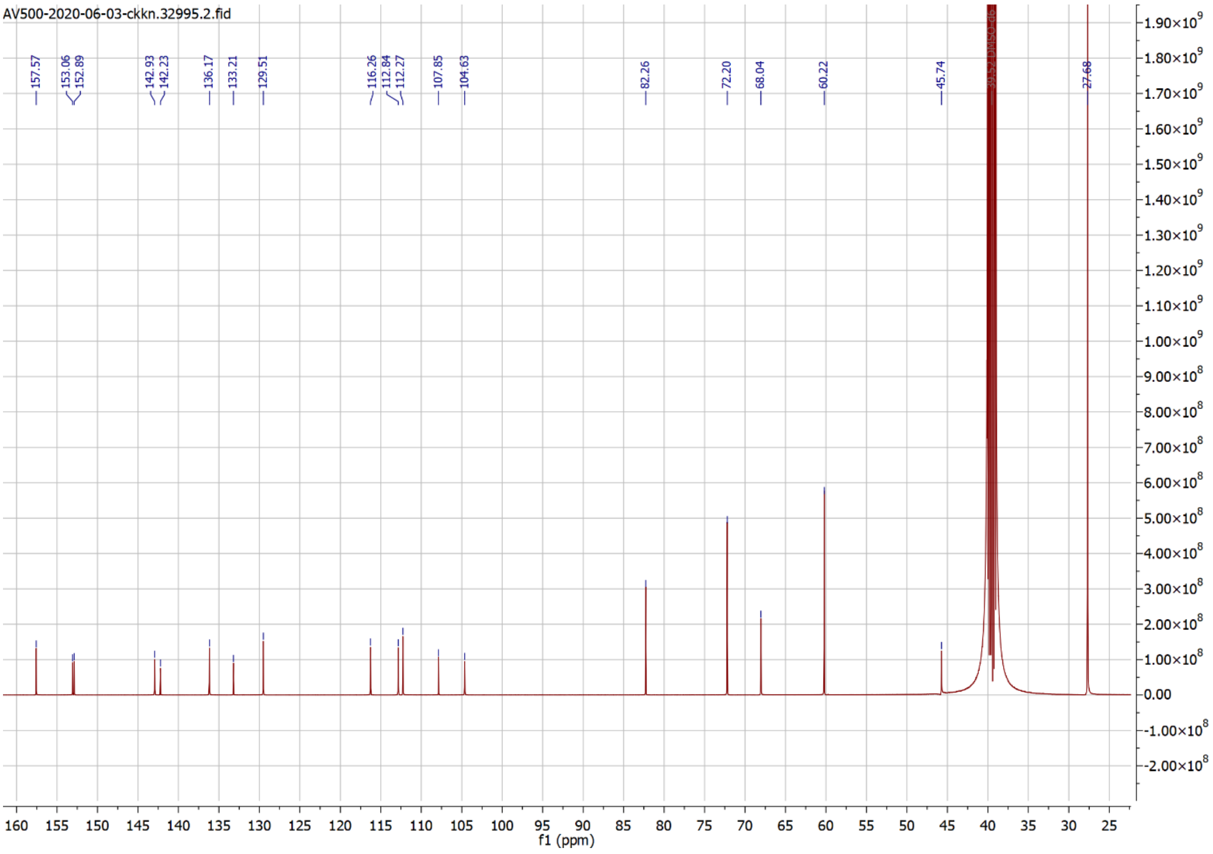
AV500-2020-06-03-ckkn.32995.2.fid
157.57
153.06
152.89
142.93
142.23
136.17
133.21
129.51
116.26
112.84
112.27
107.85
104.63
82.26
72.20
68.04
60.22
45.74
27.68
f1 (ppm)

### Compound 26

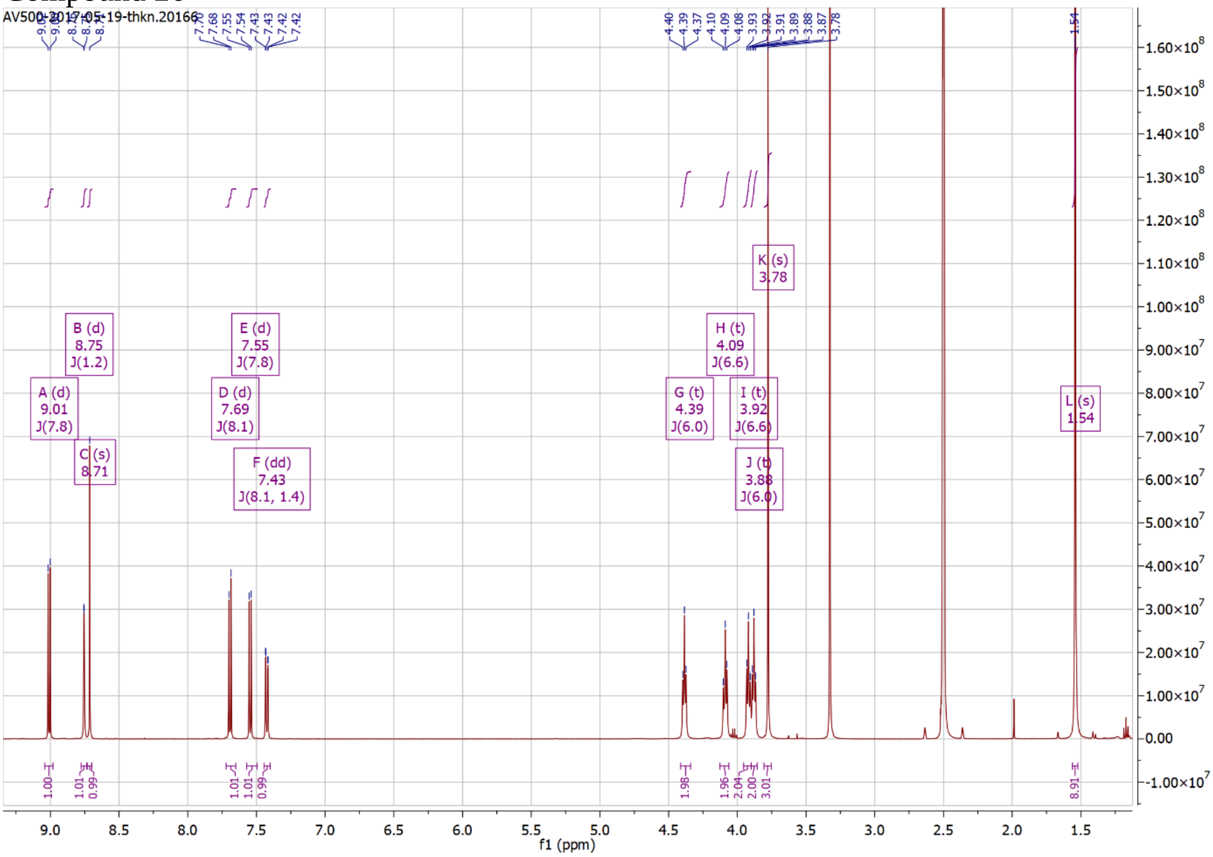
AV500-2017-05-19-thkn.20166
A (d) 9.01 J(7.8)
B (d) 8.75 J(1.2)
C (s) 8.71
D (d) 7.69 J(8.1)
E (d) 7.55 J(7.8)
F (dd) 7.43 J(8.1, 1.4)
G (t) 4.39 J(6.0)
H (t) 4.09 J(6.6)
I (t) 3.92 J(6.6)
J (t) 3.88 J(6.0)
K (s) 3.78
L (s) 1.54
f1 (ppm)

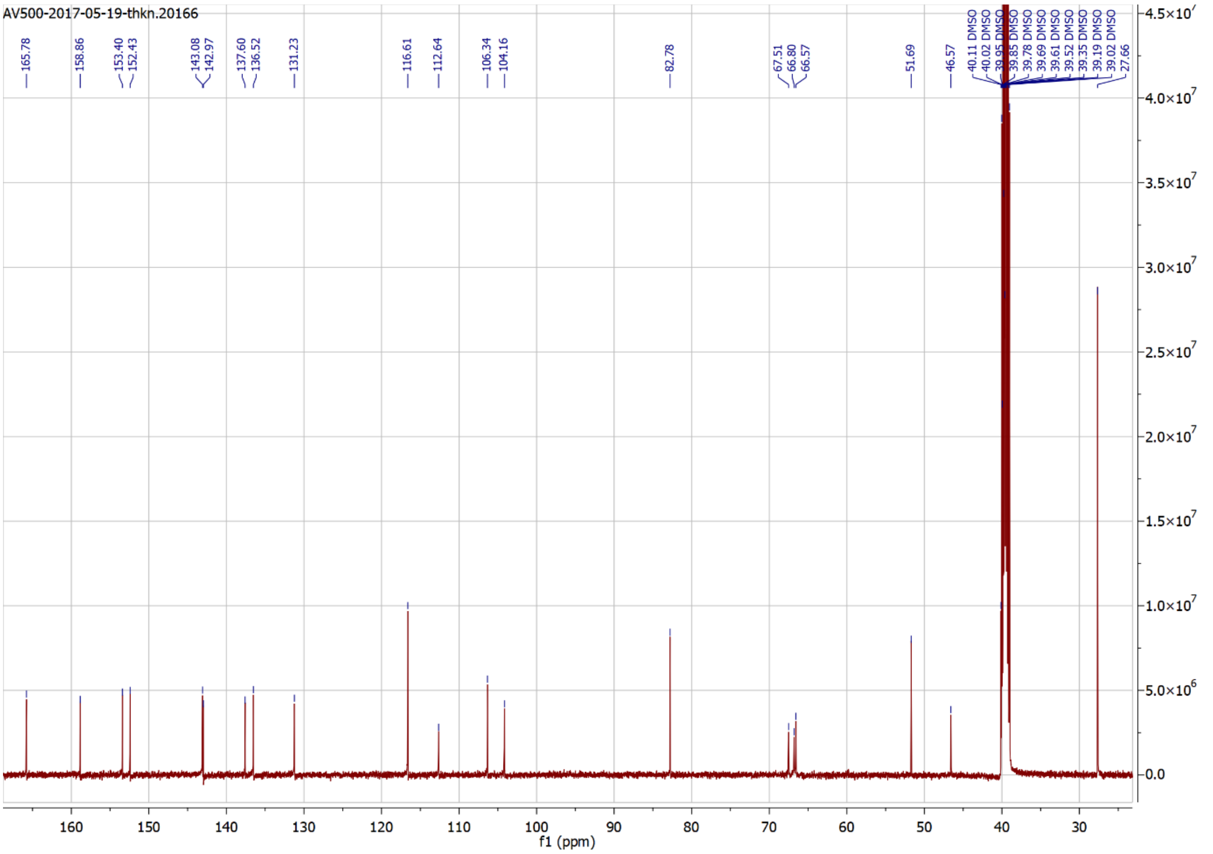
AV500-2017-05-19-thkn.20166
165.78
158.86
153.40
152.43
143.08
142.97
137.60
136.52
131.23
116.61
112.64
106.34
104.16
82.78
67.51
66.80
66.57
51.69
46.57
27.66
f1 (ppm)

### Compound 27

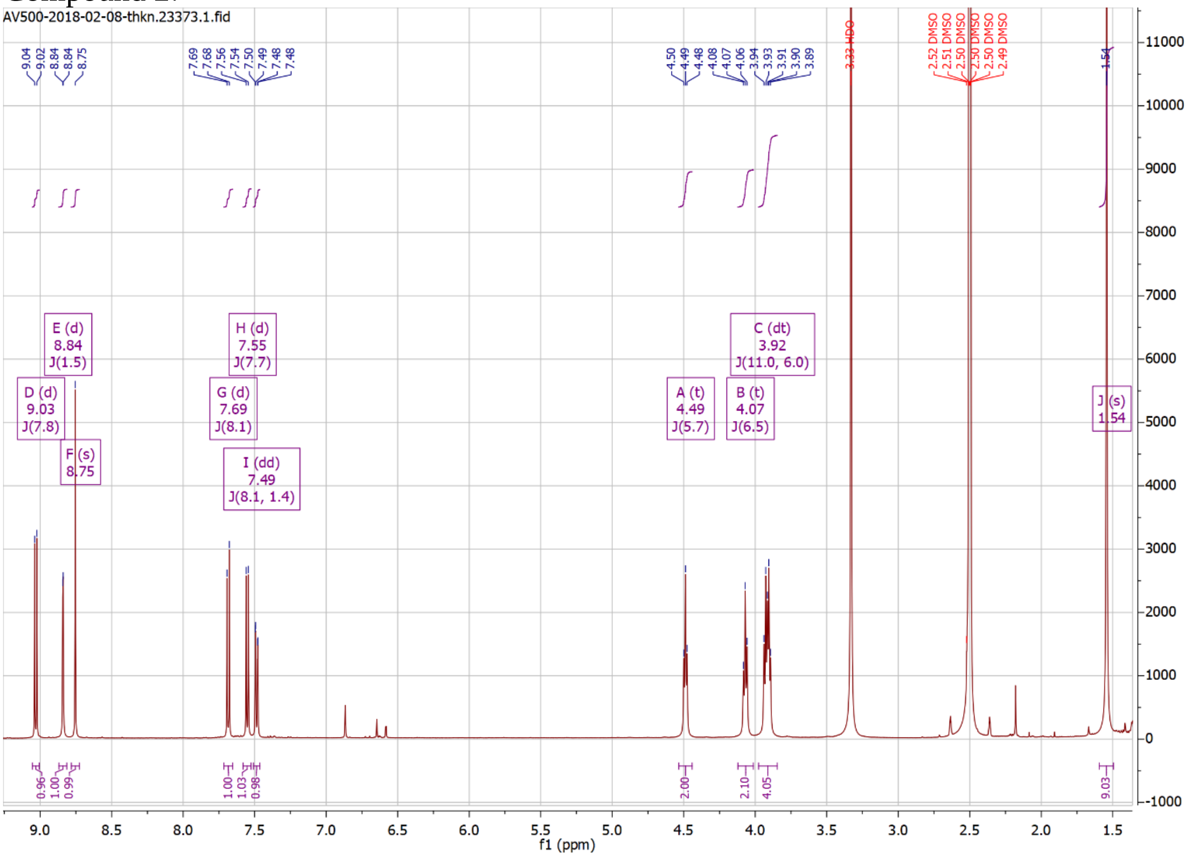
AV500-2018-02-08-thkn.23373.1.fid
A (t) 4.49 J(5.7)
B (t) 4.07 J(6.5)
C (dt) 3.92 J(11.0, 6.0)
D (d) 9.03 J(7.8)
E (d) 8.84 J(1.5)
F (s) 8.75
G (d) 7.69 J(8.1)
H (d) 7.55 J(7.7)
I (dd) 7.49 J(8.1, 1.4)
J (s) 1.54
3.33 HDO
f1 (ppm)

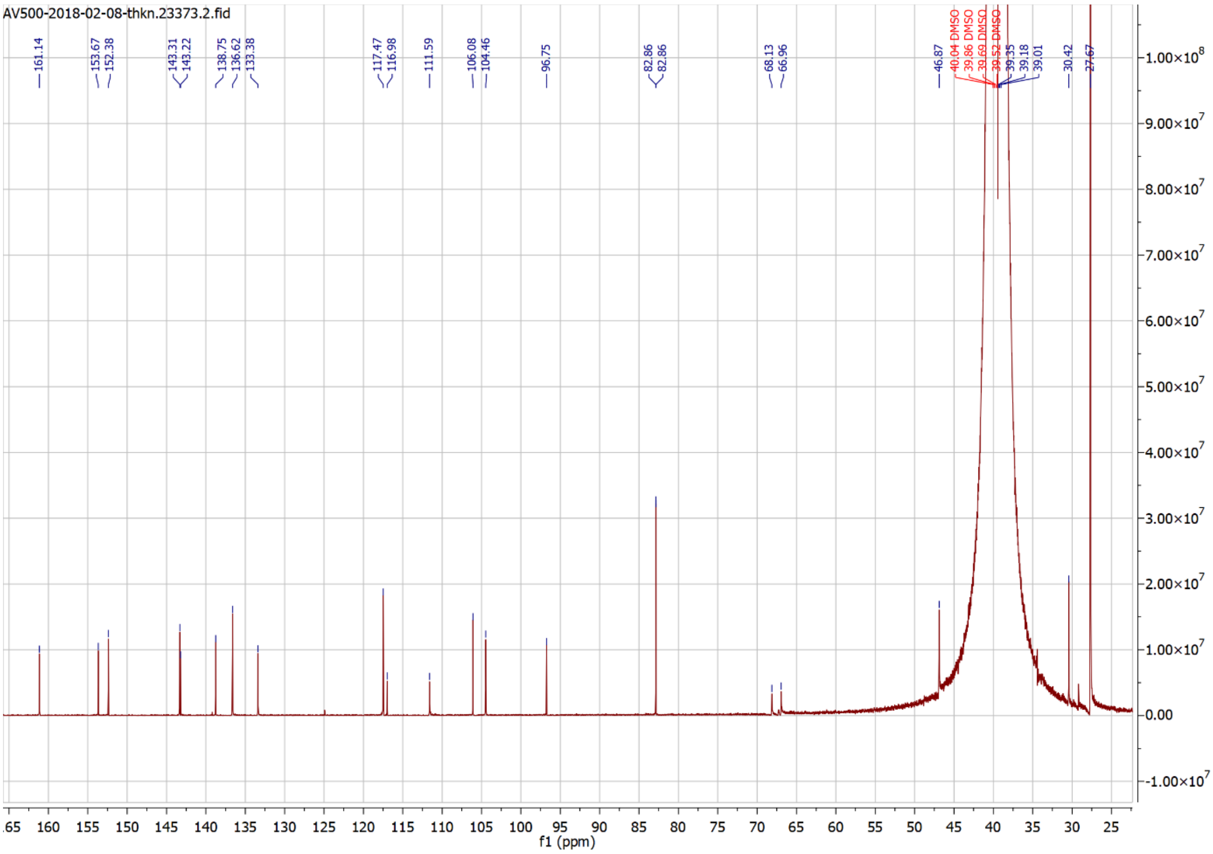
AV500-2018-02-08-thkn.23373.2.fid
161.14
153.67
152.38
143.31
143.22
138.75
136.62
133.38
117.47
116.98
111.59
106.08
104.46
96.75
82.86
82.86
68.13
66.96
46.87
30.42
27.67
f1 (ppm)

Compound 28

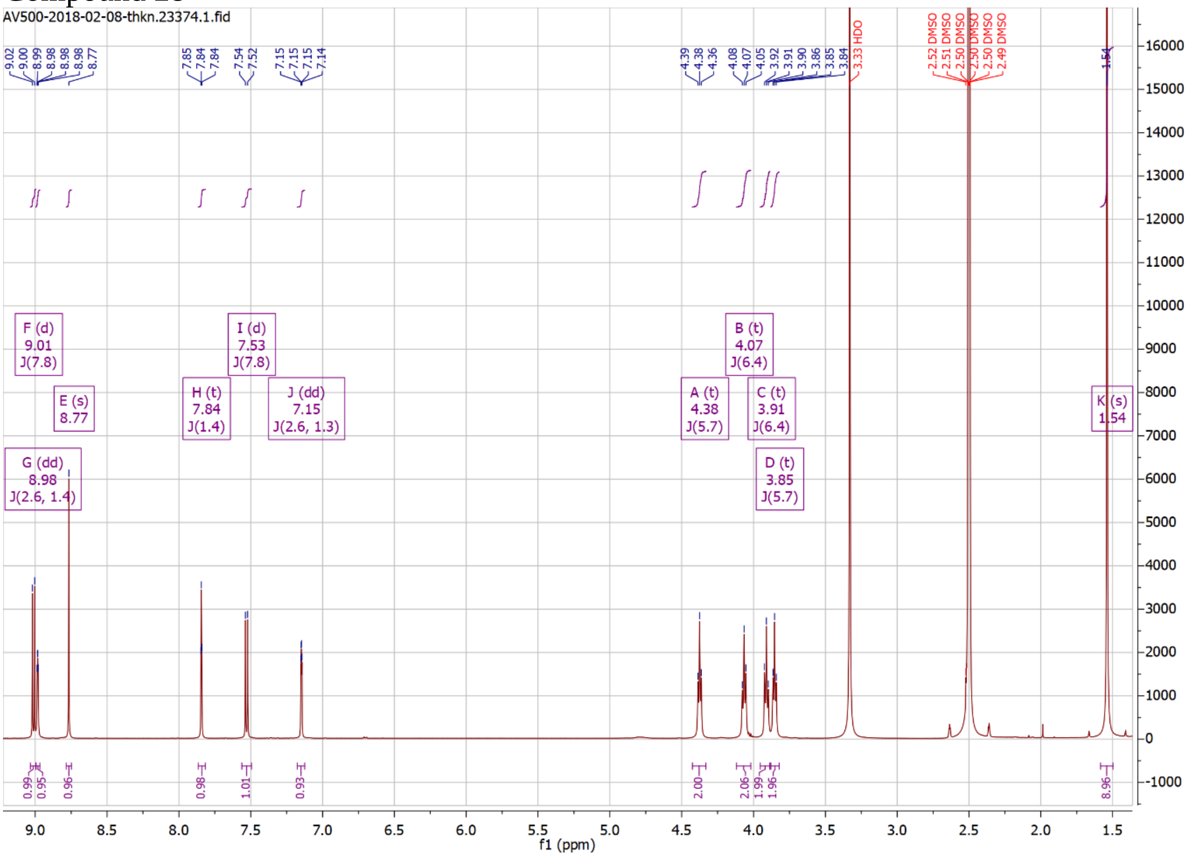
AV500-2018-02-08-thkn.23374.1.fid
F (d) 9.01 J(7.8)
E (s) 8.77
G (dd) 8.98 J(2.6, 1.4)
H (t) 7.84 J(1.4)
I (d) 7.53 J(7.8)
J (dd) 7.15 J(2.6, 1.3)
A (t) 4.38 J(5.7)
B (t) 4.07 J(6.4)
C (t) 3.91 J(6.4)
D (t) 3.85 J(5.7)
K (s) 1.54
3.33 HDO
f1 (ppm)

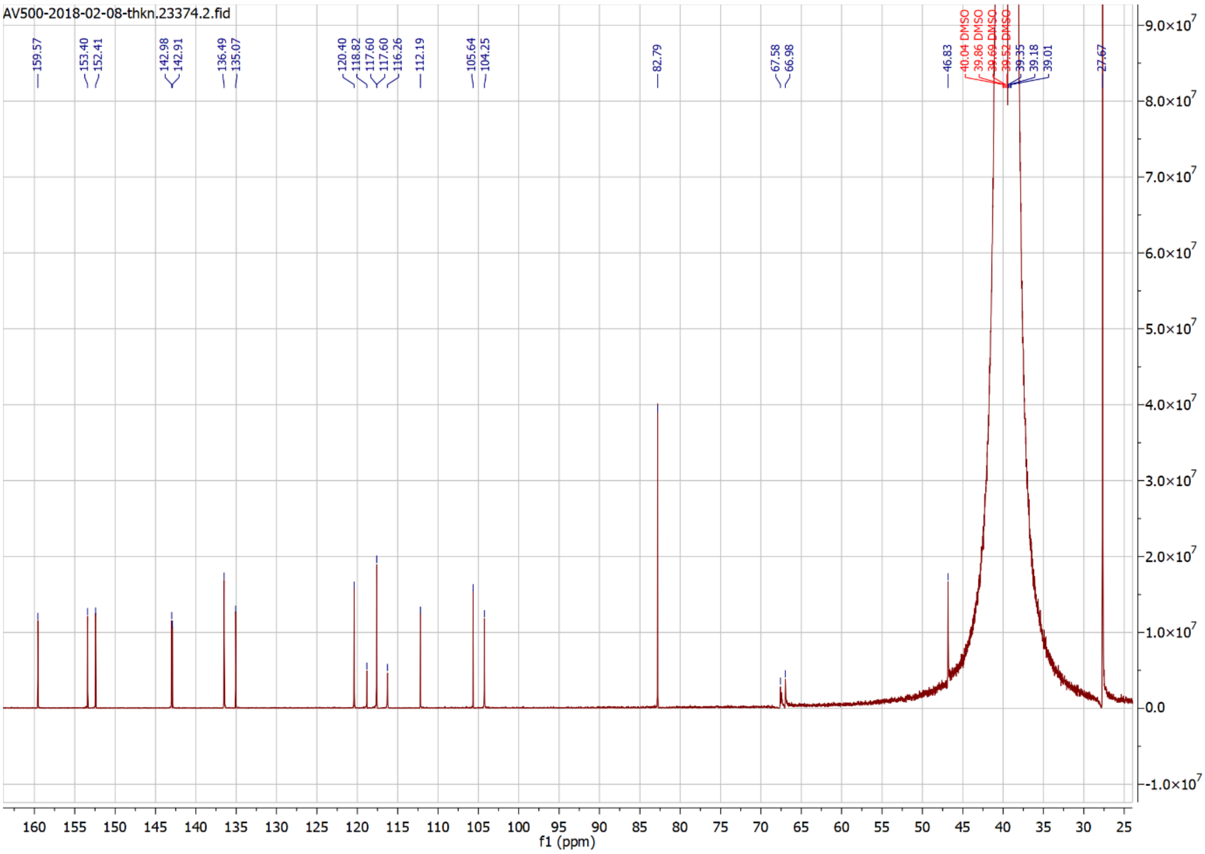
AV500-2018-02-08-thkn.23374.2.fid
159.57
153.40
152.41
142.98
142.91
136.49
135.07
120.40
118.82
117.60
117.60
116.26
112.19
105.64
104.25
82.79
67.58
66.98
46.83
27.67
f1 (ppm)

Compound 29

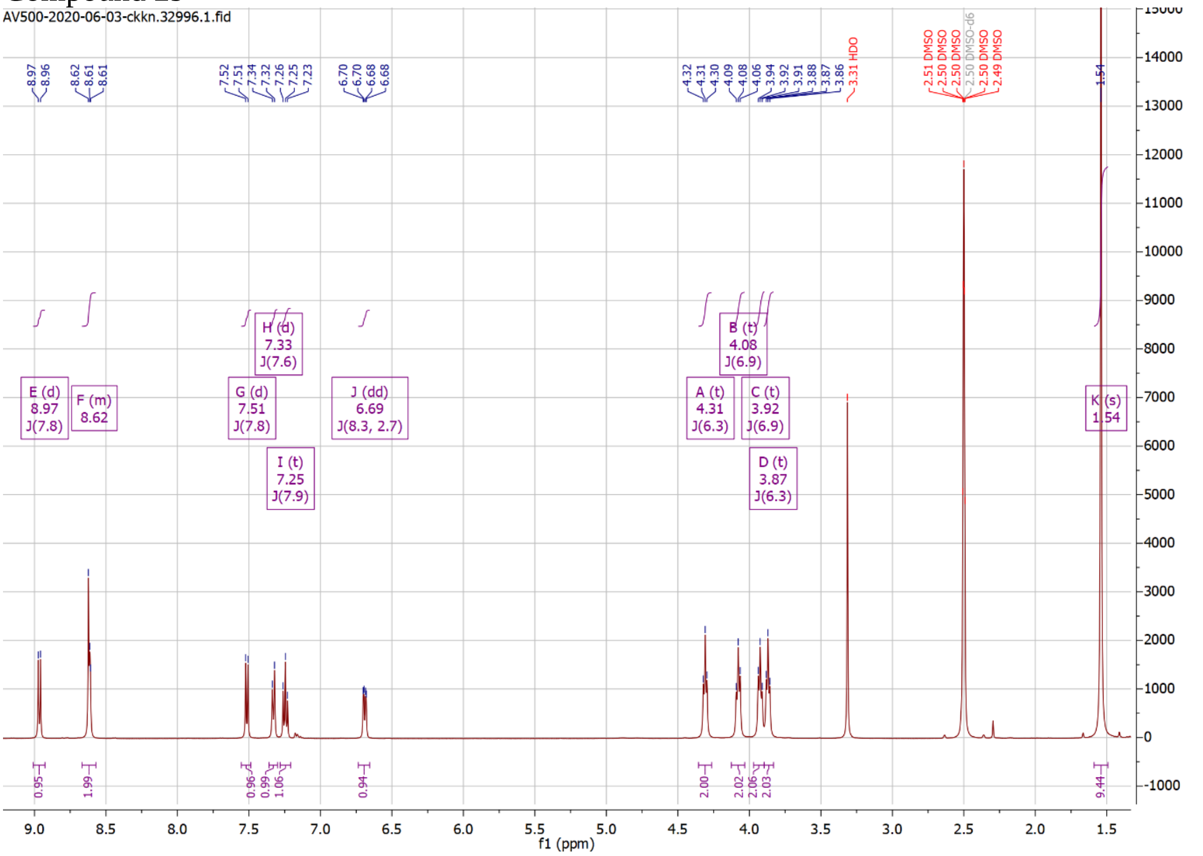
AV500-2020-06-03-ckkn.32996.1.fid
E (d) 8.97 J(7.8)
F (m) 8.62
G (d) 7.51 J(7.8)
H (d) 7.33 J(7.6)
I (t) 7.25 J(7.9)
J (dd) 6.69 J(8.3, 2.7)
A (t) 4.31 J(6.3)
B (t) 4.08 J(6.9)
C (t) 3.92 J(6.9)
D (t) 3.87 J(6.3)
K (s) 1.54
3.31 HDO
2.50 DMSO-d6
f1 (ppm)

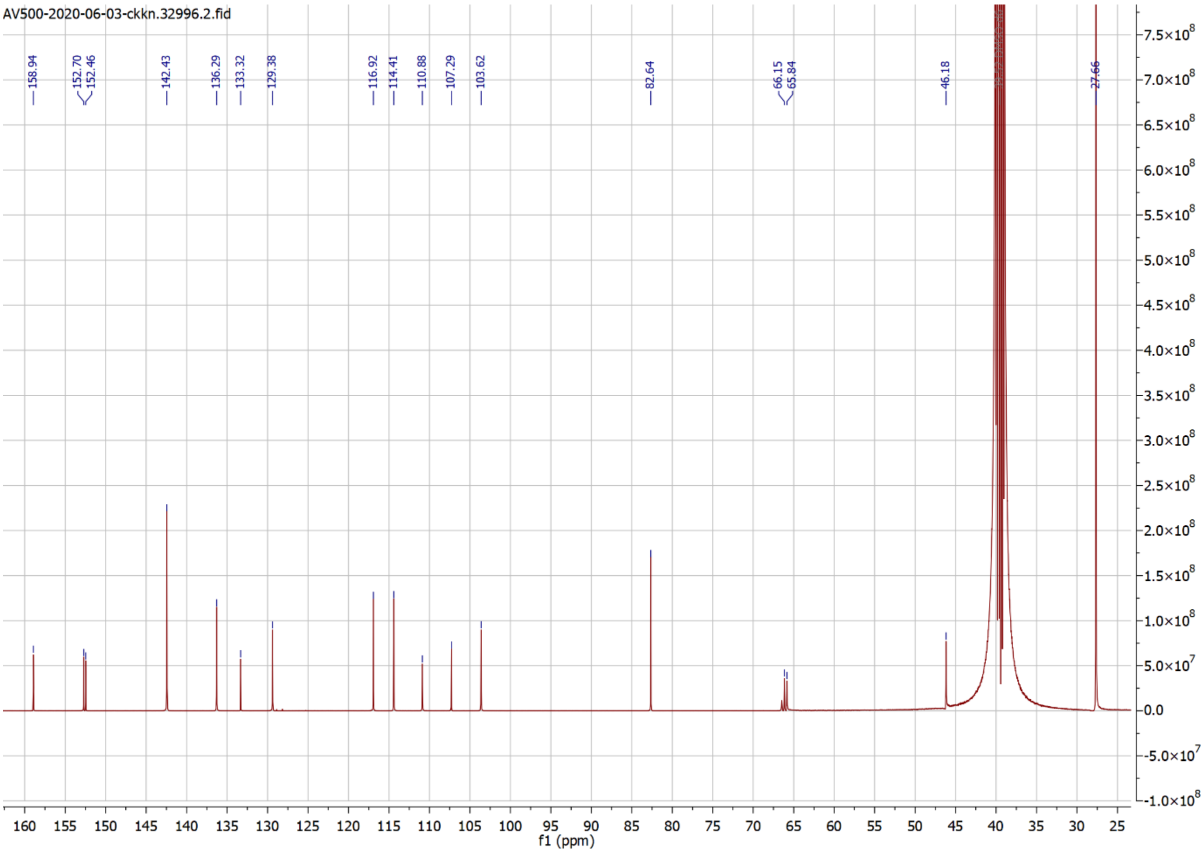
AV500-2020-06-03-ckkn.32996.2.fid
158.94
152.70
152.46
142.43
136.29
133.32
129.38
116.92
114.41
110.88
107.29
103.62
82.64
66.15
65.84
46.18
27.66
f1 (ppm)

Compound 30

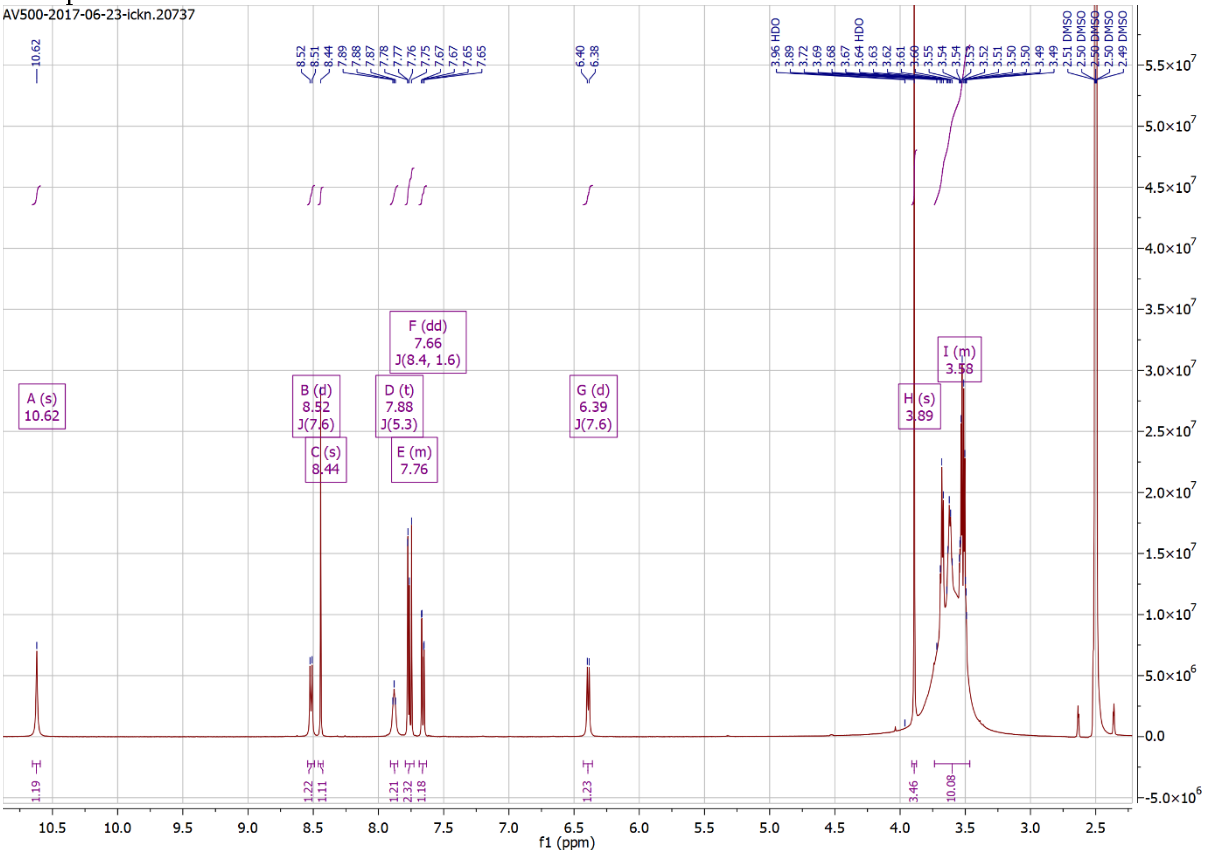

AV500-2017-06-23-ickn.20737
A (s) 10.62
B (d) 8.52 J(7.6)
C (s) 8.44
D (t) 7.88 J(5.3)
E (m) 7.76
F (dd) 7.66 J(8.4, 1.6)
G (d) 6.39 J(7.6)
H (s) 3.89
I (m) 3.58
f1 (ppm)

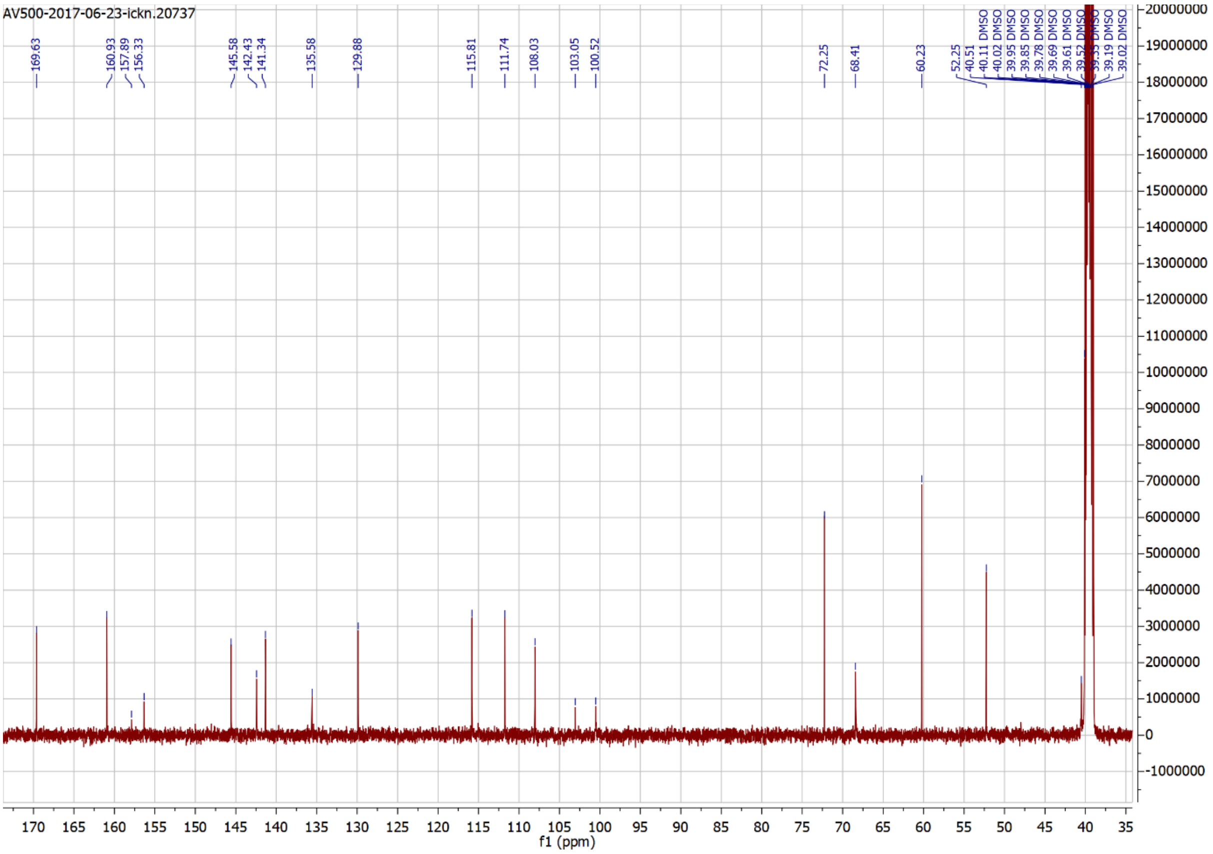

AV500-2017-06-23-ickn.20737
169.63
160.93
157.89
156.33
145.58
142.43
141.34
135.58
129.88
115.81
111.74
108.03
103.05
100.52
72.25
68.41
60.23
52.25
f1 (ppm)

### Compound 31

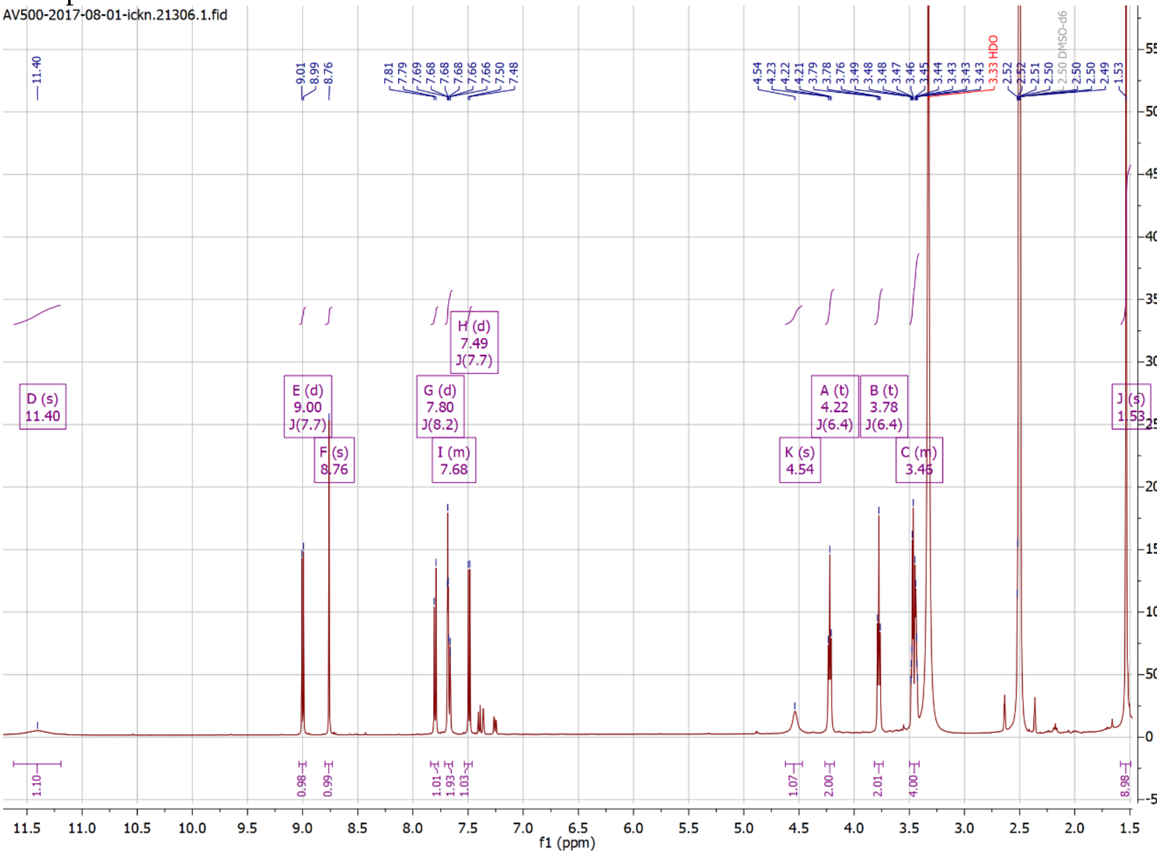
AV500-2017-08-01-ickn.21306.1.fid
D (s) 11.40
E (d) 9.00 J(7.7)
F (s) 8.76
G (d) 7.80 J(8.2)
H (d) 7.49 J(7.7)
I (m) 7.68
K (s) 4.54
A (t) 4.22 J(6.4)
B (t) 3.78 J(6.4)
C (m) 3.46
J (s) 1.53
3.33 HDO
2.50 DMSO-d6
f1 (ppm)

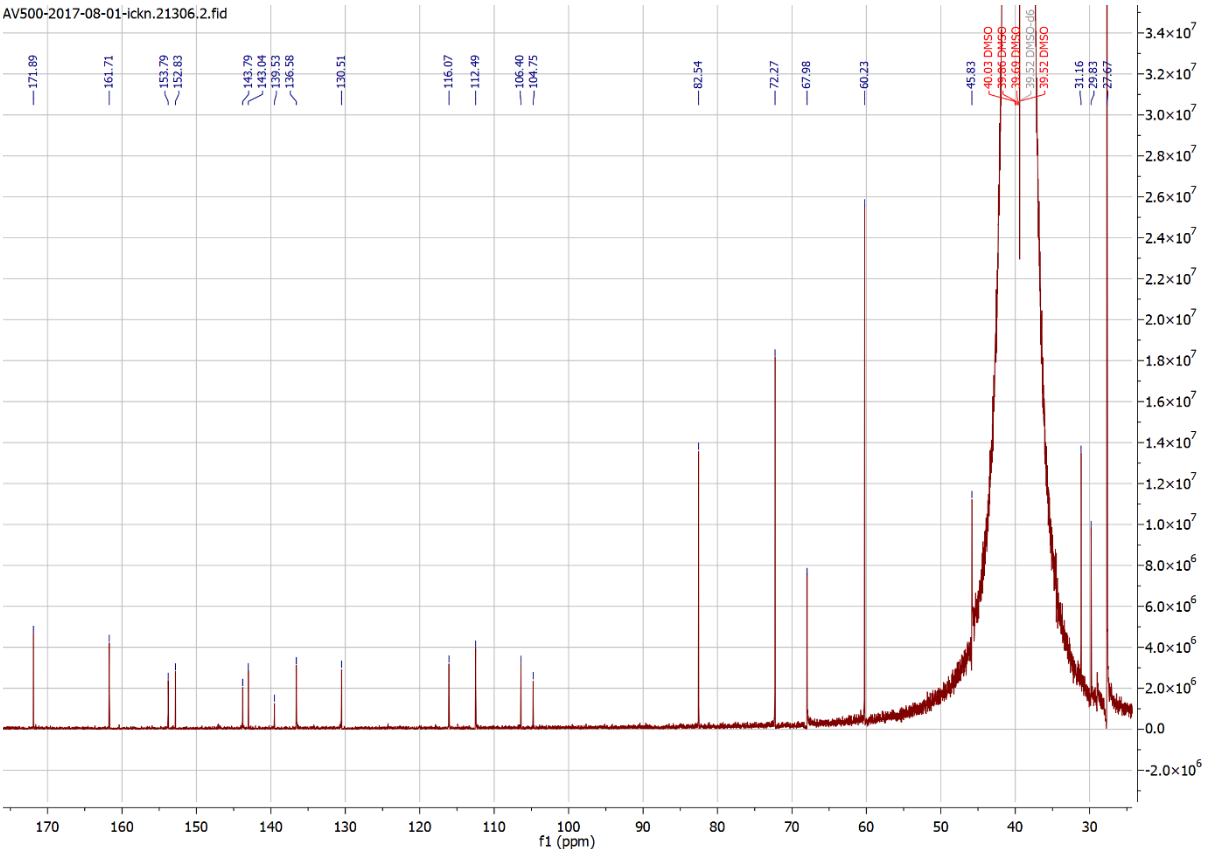
AV500-2017-08-01-ickn.21306.2.fid
171.89
161.71
153.79
152.83
143.79
143.04
139.53
136.58
130.51
116.07
112.49
106.40
104.75
82.54
72.27
67.98
60.23
45.83
40.03 DMSO
39.86 DMSO
39.69 DMSO
39.52 DMSO-d6
39.52 DMSO
31.16
29.83
27.67
f1 (ppm)

Compound 32

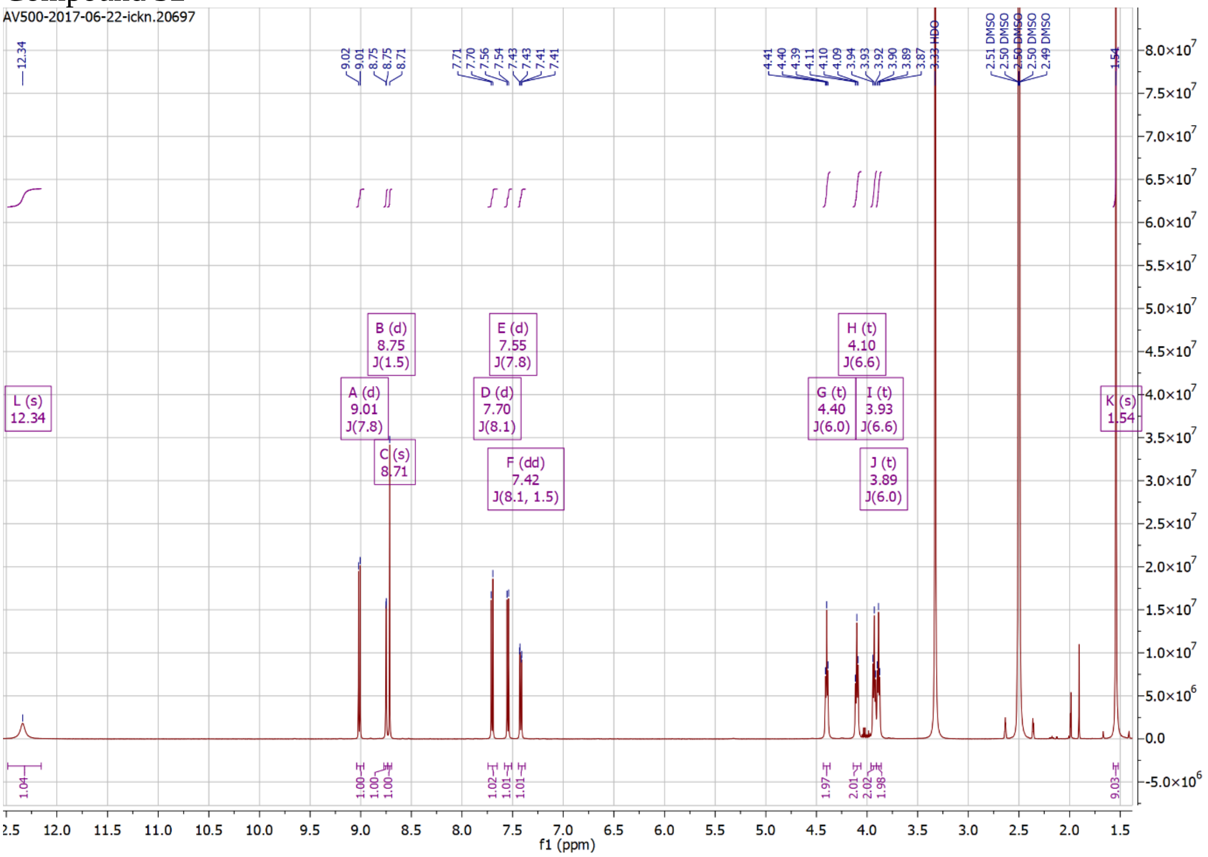
AV500-2017-06-22-ickn.20697
L (s) 12.34
A (d) 9.01 J(7.8)
B (d) 8.75 J(1.5)
C (s) 8.71
D (d) 7.70 J(8.1)
E (d) 7.55 J(7.8)
F (dd) 7.42 J(8.1, 1.5)
G (t) 4.40 J(6.0)
H (t) 4.10 J(6.6)
I (t) 3.93 J(6.6)
J (t) 3.89 J(6.0)
K (s) 1.54
3.33 HDO
2.50 DMSO
f1 (ppm)

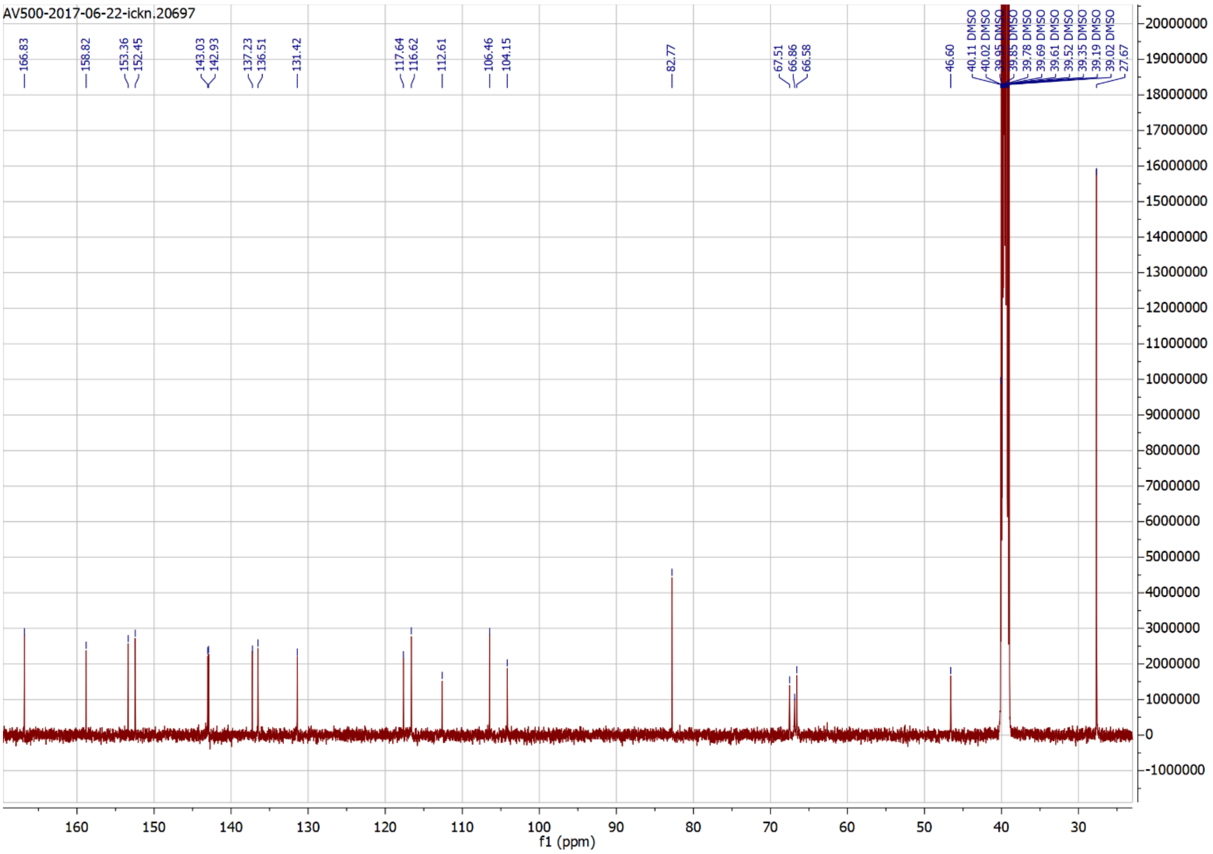
AV500-2017-06-22-ickn.20697
166.83
158.82
153.36
152.45
143.03
142.93
137.23
136.51
131.42
117.64
116.62
112.61
106.46
104.15
82.77
67.51
66.86
66.58
46.60
27.67
f1 (ppm)

### Compound 33

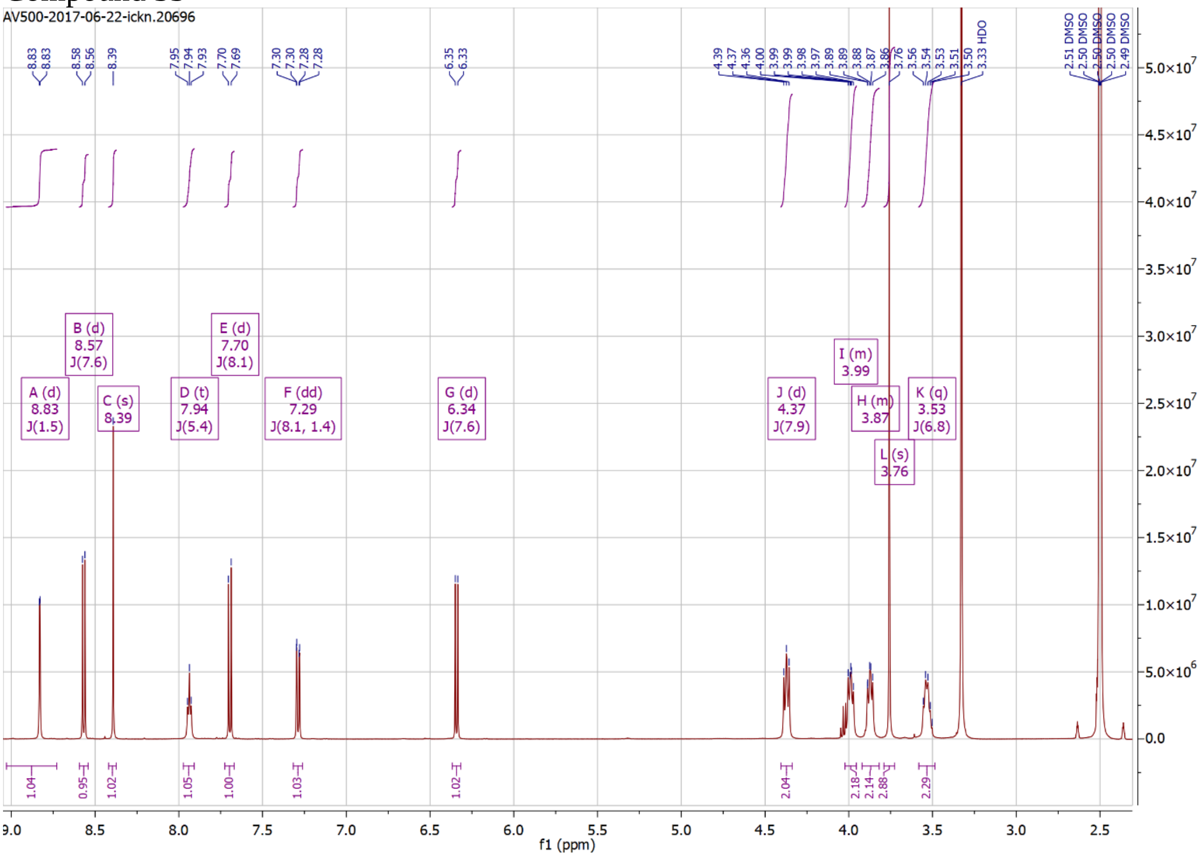
AV500-2017-06-22-ickn.20696
A (d) 8.83 J(1.5)
B (d) 8.57 J(7.6)
C (s) 8.39
D (t) 7.94 J(5.4)
E (d) 7.70 J(8.1)
F (dd) 7.29 J(8.1, 1.4)
G (d) 6.34 J(7.6)
J (d) 4.37 J(7.9)
I (m) 3.99
H (m) 3.87
L (s) 3.76
K (q) 3.53 J(6.8)
3.33 HDO
f1 (ppm)

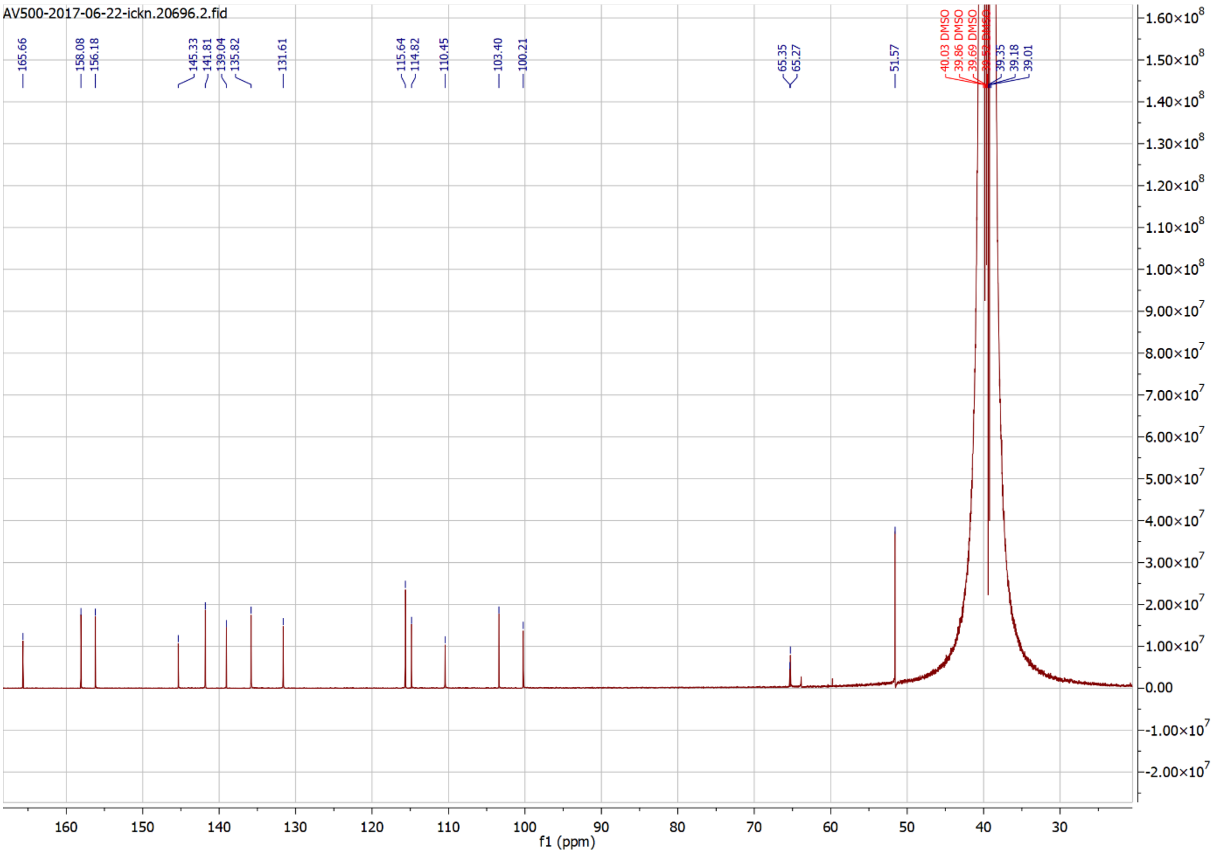
AV500-2017-06-22-ickn.20696.2.fid
165.66
158.08
156.18
145.33
141.81
139.04
135.82
131.61
115.64
114.82
110.45
103.40
100.21
65.35
65.27
51.57
39.35
39.18
39.01
f1 (ppm)

### Compound 34

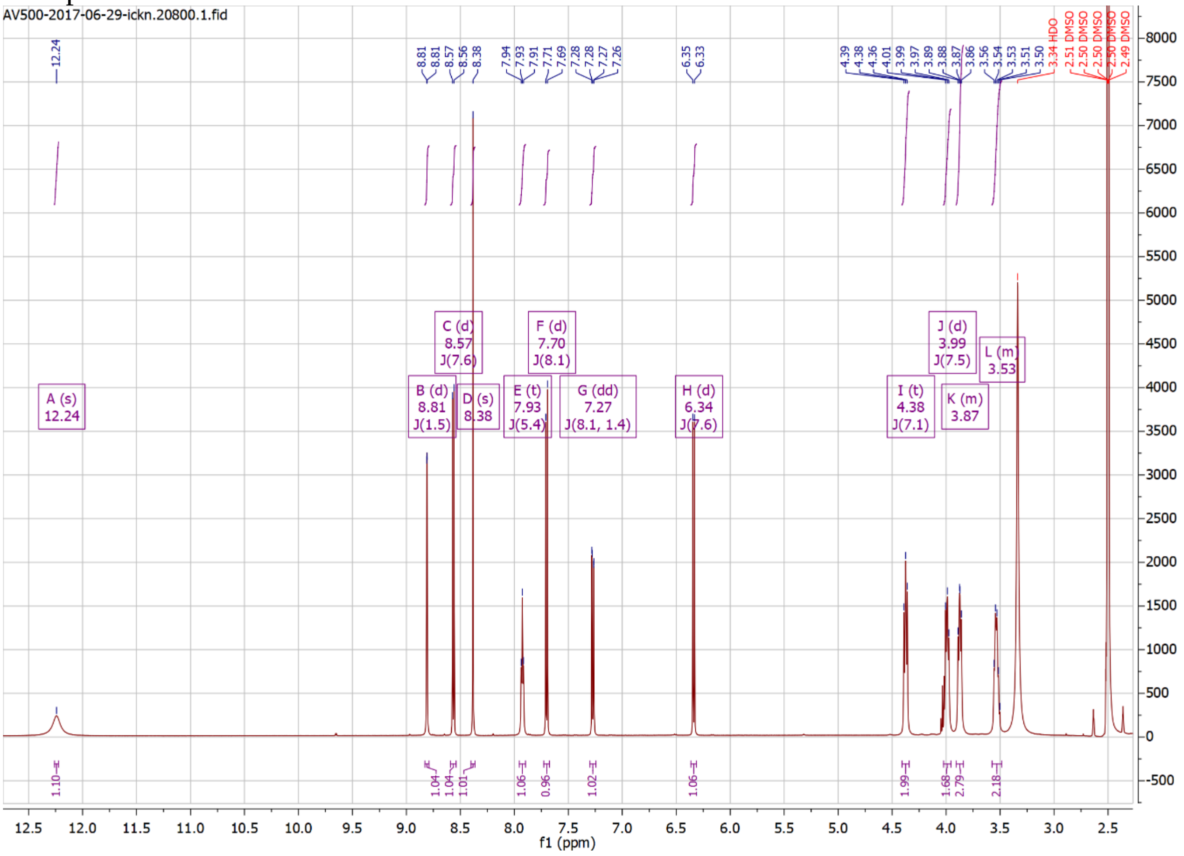
AV500-2017-06-29-ickn.20800.1.fid
A (s) 12.24
B (d) 8.81 J(1.5)
C (d) 8.57 J(7.6)
D (s) 8.38
E (t) 7.93 J(5.4)
F (d) 7.70 J(8.1)
G (dd) 7.27 J(8.1, 1.4)
H (d) 6.34 J(7.6)
I (t) 4.38 J(7.1)
J (d) 3.99 J(7.5)
K (m) 3.87
L (m) 3.53
f1 (ppm)

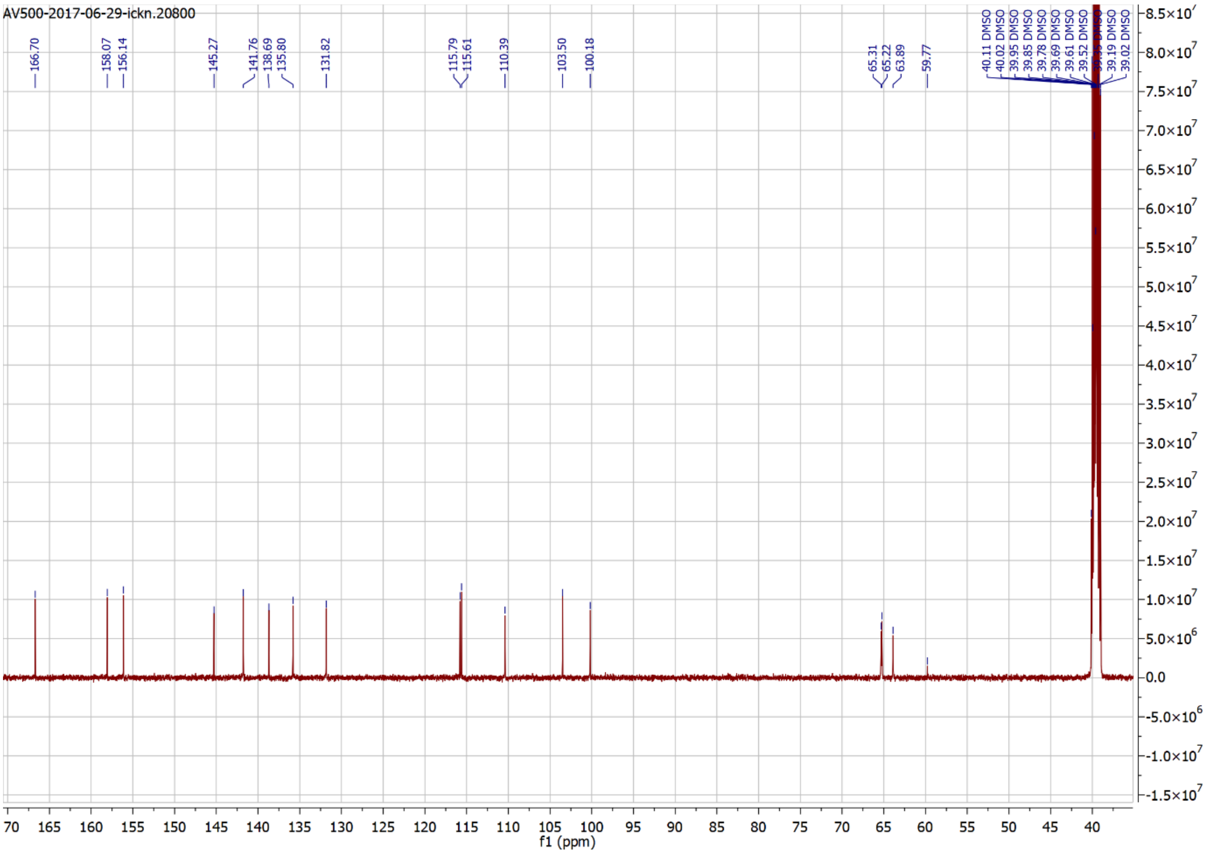
AV500-2017-06-29-ickn.20800
166.70
158.07
156.14
145.27
141.76
138.69
135.80
131.82
115.79
115.61
110.39
103.50
100.18
65.31
65.22
63.89
59.77
f1 (ppm)

Compound 35

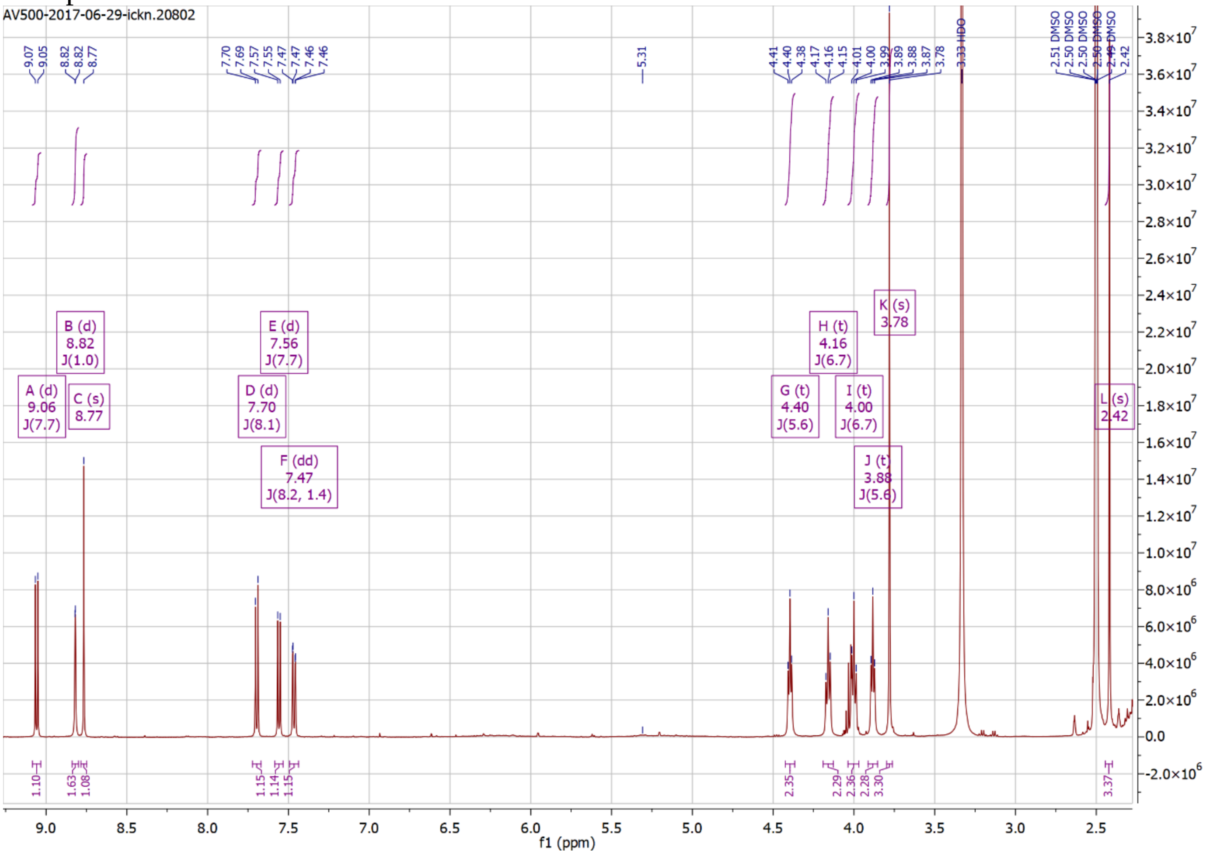

AV500-2017-06-29-ickn.20802
A (d) 9.06 J(7.7)
B (d) 8.82 J(1.0)
C (s) 8.77
D (d) 7.70 J(8.1)
E (d) 7.56 J(7.7)
F (dd) 7.47 J(8.2, 1.4)
G (t) 4.40 J(5.6)
H (t) 4.16 J(6.7)
I (t) 4.00 J(6.7)
J (t) 3.88 J(5.6)
K (s) 3.78
L (s) 2.42
3.33 HDO
f1 (ppm)

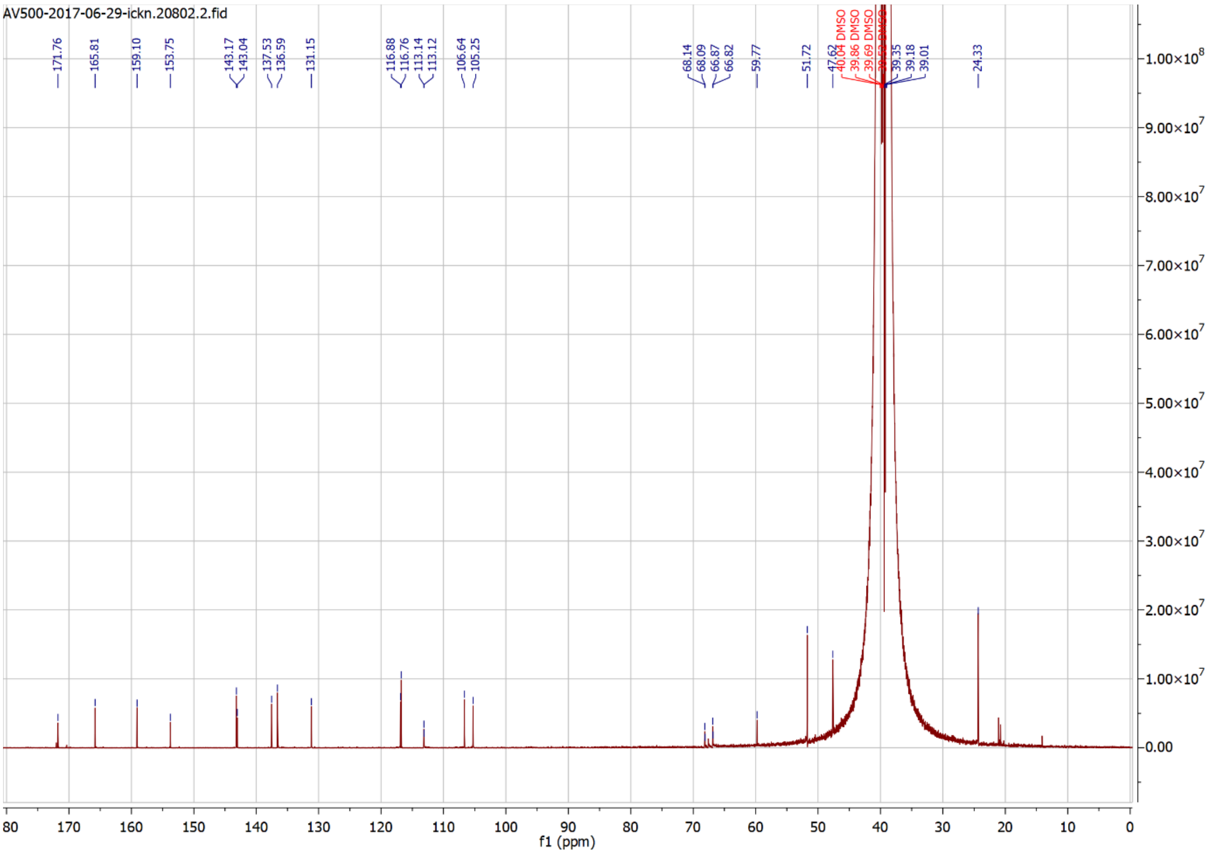

AV500-2017-06-29-ickn.20802.2.fid
171.76
165.81
159.10
153.75
143.17
143.04
137.53
136.59
131.15
116.88
116.76
113.14
113.12
106.64
105.25
68.14
68.09
66.87
66.82
59.77
51.72
47.62
24.33
f1 (ppm)

Compound 36

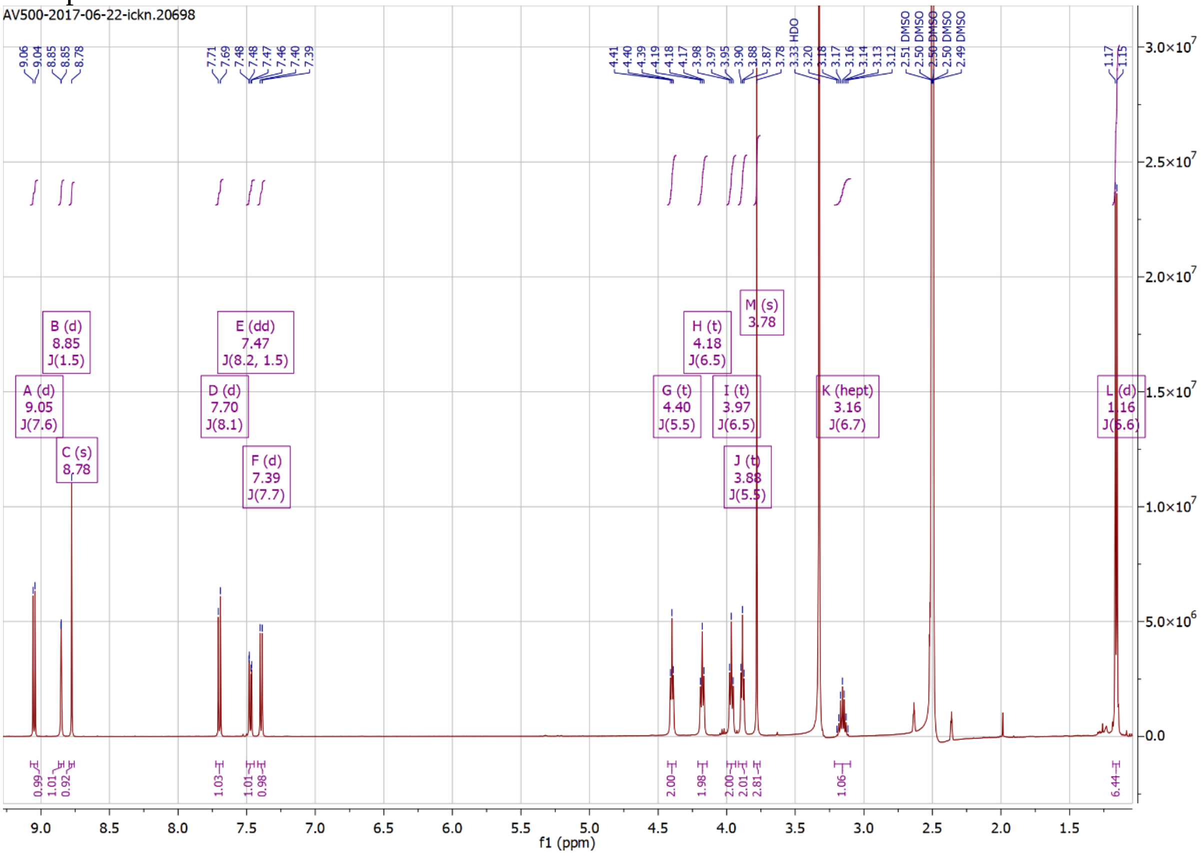
AV500-2017-06-22-ickn.20698
A (d) 9.05 J(7.6)
B (d) 8.85 J(1.5)
C (s) 8.78
D (d) 7.70 J(8.1)
E (dd) 7.47 J(8.2, 1.5)
F (d) 7.39 J(7.7)
G (t) 4.40 J(5.5)
H (t) 4.18 J(6.5)
I (t) 3.97 J(6.5)
J (t) 3.88 J(5.5)
M (s) 3.78
K (hept) 3.16 J(6.7)
L (d) 1.16 J(6.6)
f1 (ppm)

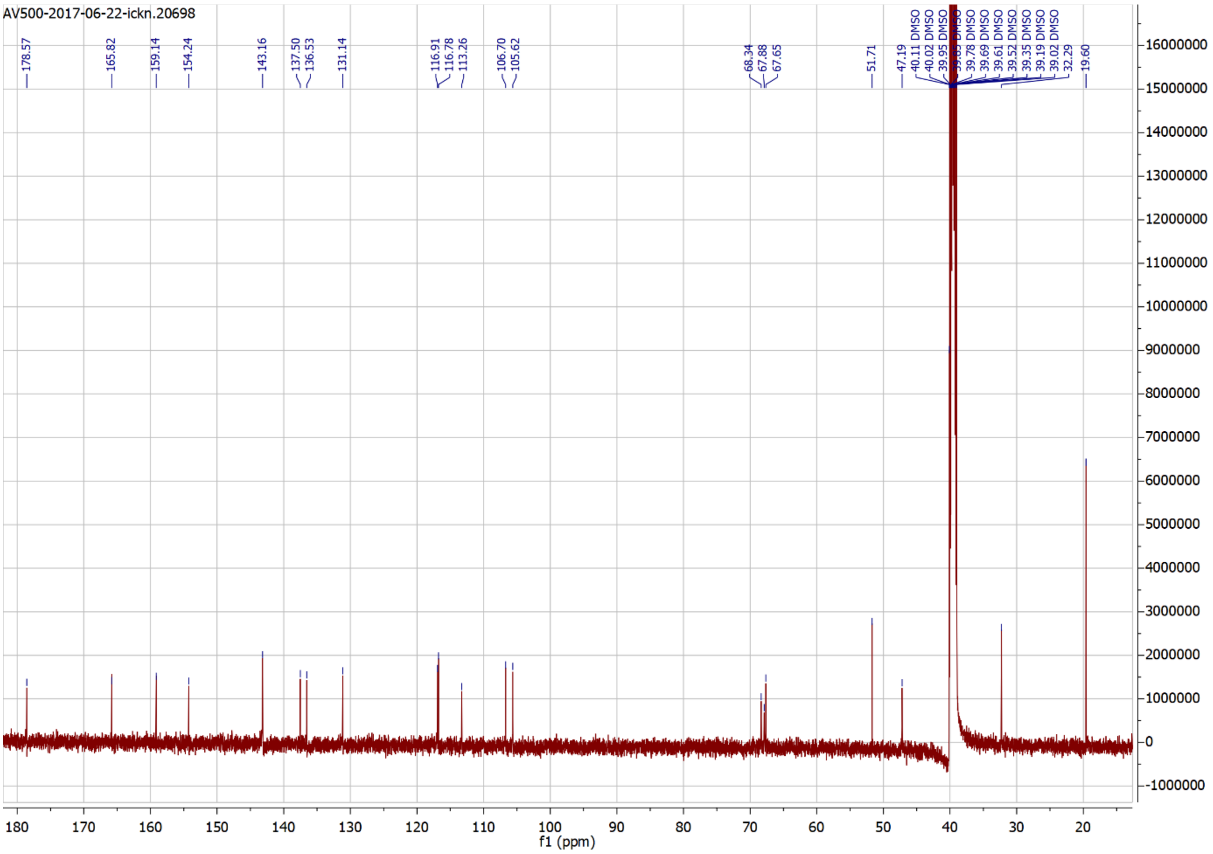
AV500-2017-06-22-ickn.20698
178.57
165.82
159.14
154.24
143.16
137.50
136.53
131.14
116.91
116.78
113.26
106.70
105.62
68.34
67.88
67.65
51.71
47.19
32.29
19.60
f1 (ppm)

Compound 37

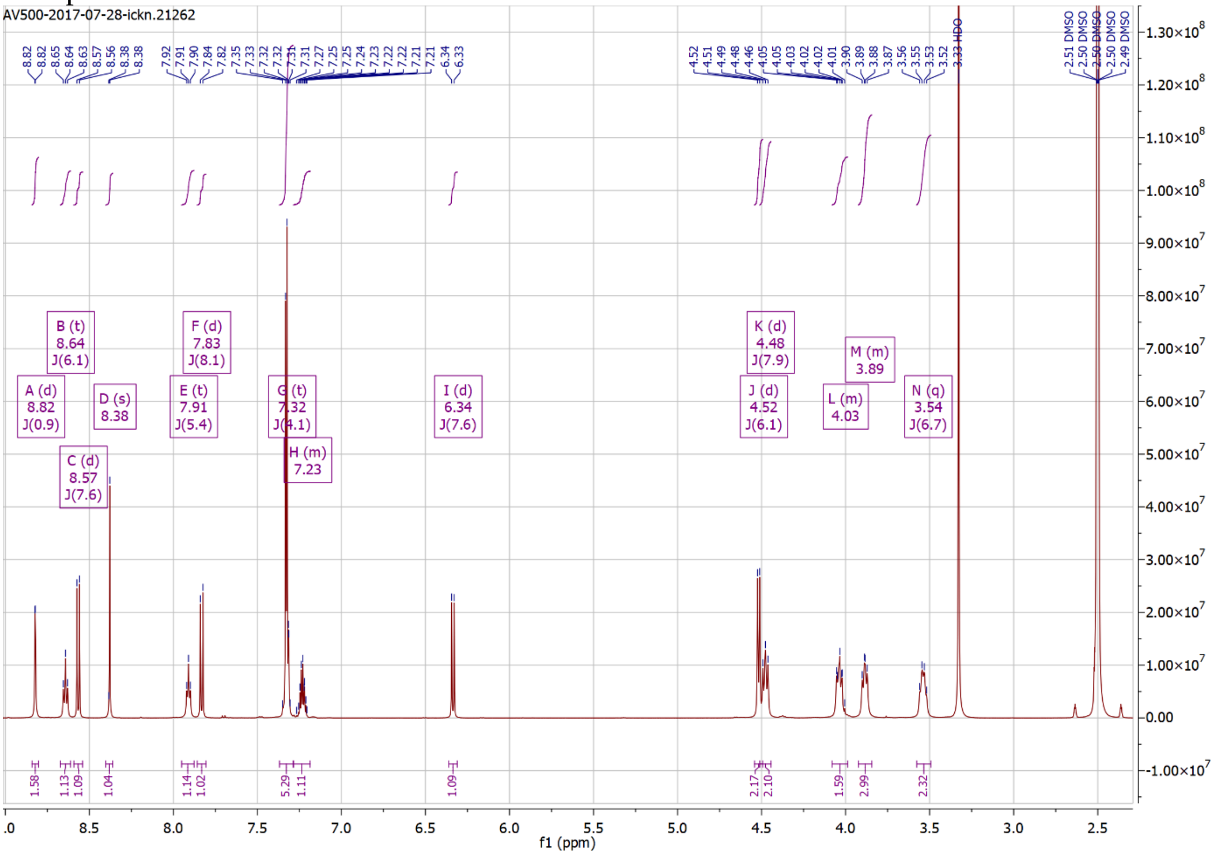
AV500-2017-07-28-ickn.21262
A (d) 8.82 J(0.9)
B (t) 8.64 J(6.1)
C (d) 8.57 J(7.6)
D (s) 8.38
E (t) 7.91 J(5.4)
F (d) 7.83 J(8.1)
G (t) 7.32 J(4.1)
H (m) 7.23
I (d) 6.34 J(7.6)
J (d) 4.52 J(6.1)
K (d) 4.48 J(7.9)
L (m) 4.03
M (m) 3.89
N (q) 3.54 J(6.7)
3.33 HDO
f1 (ppm)

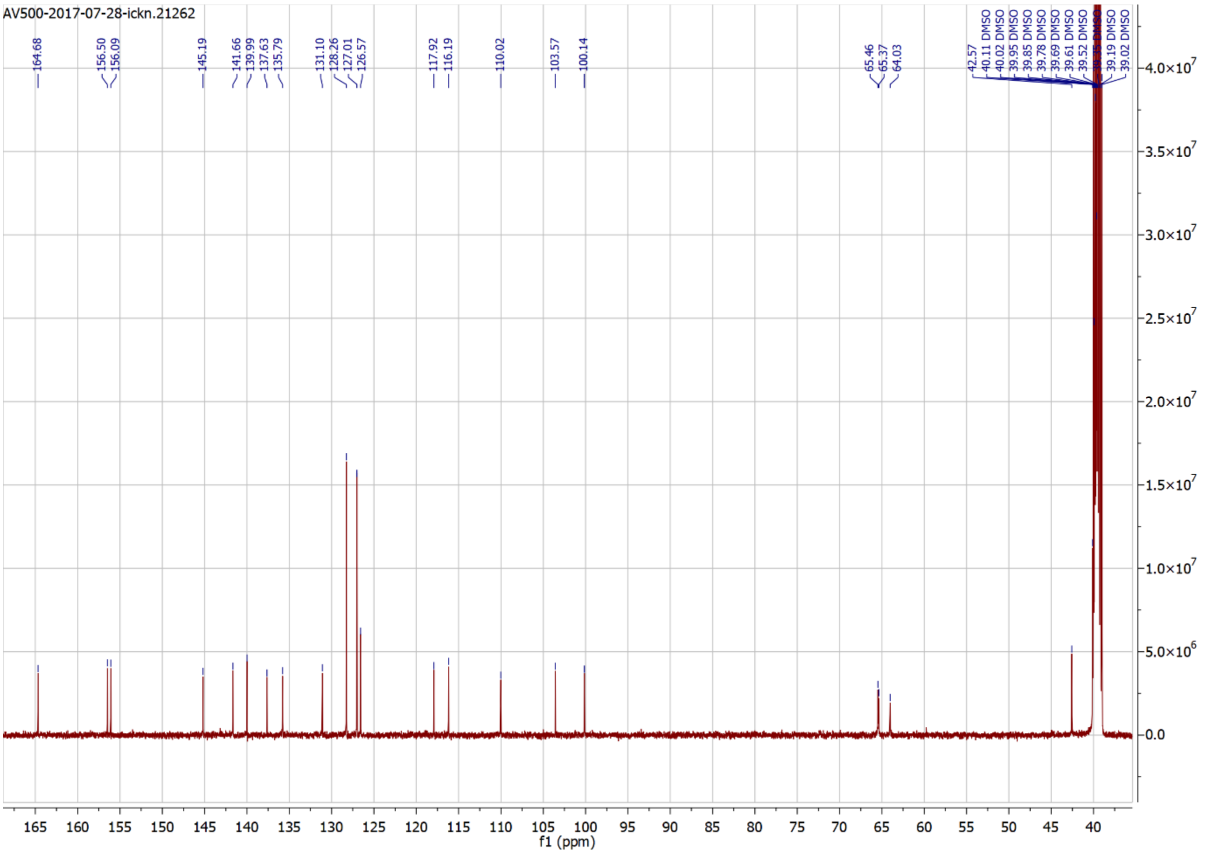
AV500-2017-07-28-ickn.21262
164.68
156.50
156.09
145.19
141.66
139.99
137.63
135.79
131.10
128.26
127.01
126.57
117.92
116.19
110.02
103.57
100.14
65.46
65.37
64.03
f1 (ppm)
